# Esr1+ hypothalamic-habenula neurons shape aversive states

**DOI:** 10.1101/2022.11.17.516965

**Authors:** Daniela Calvigioni, Janos Fuzik, Pierre Le Merre, Marina Slashcheva, Felix Jung, Cantin Ortiz, Antonio Lentini, Veronika Csillag, Marta Graziano, Ifigeneia Nikolakopoulou, Moritz Weglage, Iakovos Lazaridis, Hoseok Kim, Irene Lenzi, Hyunsoo Park, Björn Reinius, Marie Carlén, Konstantinos Meletis

**Affiliations:** Department of Neuroscience, Karolinska Institutet, Stockholm, Sweden; Department of Medical Biochemistry and Biophysics, Karolinska Institutet, Stockholm, Sweden

## Abstract

Excitatory projections from the lateral hypothalamic area (LHA) to the lateral habenula (LHb) drive aversive responses. We used Patch-seq guided multimodal classification to define the structural and functional heterogeneity of the LHA-LHb pathway. Our classification identified six glutamatergic neuron types with unique electrophysiological properties, molecular profiles, and projection patterns. We found that genetically-defined LHA-LHb neurons signal distinct aspects of emotional or naturalistic behaviors: Esr1+ LHA-LHb neurons induce aversion, whereas Npy+ LHA-LHb neurons control rearing behavior. Repeated optogenetic drive of Esr1+ LHA-LHb neurons induces a behaviorally persistent aversive state, and large-scale recordings showed a region-specific neural representation of the aversive state in the prelimbic region of the prefrontal cortex. We further found that exposure to unpredictable mild shocks induced a sex-specific sensitivity to develop a stress state in female mice, which was associated with a specific shift in the intrinsic properties of bursting-type Esr1+ LHA-LHb neurons. In summary, we describe the diversity of LHA-LHb neuron types, and provide evidence for the role of Esr1+ neurons in aversion and sexually dimorphic stress sensitivity.

## Main

Behaviors based on emotional processing (emotional behaviors) are regulated by internal and external value and state signals, where negative signals for example lead to avoidance (*1*). Emotional behaviors and affective disorders are thought to depend on subcortical as well as cortical circuitries, where the LHb and the PFC are key nodes in value and emotion processing (*2*). The prefrontal cortex (PFC) is a key network controlling cognitive and emotional behavior (*3, 4*) and the LHb is known to integrate value and emotion signals (*5*). Importantly, the main excitatory input to LHb originates from the lateral hypothalamic area (LHA), and this LHA-LHb pathway can signal avoidance and the expression of depression-like states (*6–9*). Affective disorders such as depression are more common in women (*10*) and the sensitivity to negative events show sex differences (*11, 12*). The hypothalamus is a highly heterogeneous region, composed of many subregions (*13*) containing different neuron types defined by differential gene expression (*14*) and projection pattern (*15, 16*), including sexually dimorphic circuits and behavior (*17, 18*) . In spite of the evident cellular and functional diversity in other hypothalamic regions, the LHA-LHb pathway has so far been studied as a homogeneous glutamatergic population (*6*–*8, 19, 20*).

To define the organization and function of the glutamatergic (Vglut2+) LHA-LHb pathway, we used a multimodal classification of neuron types based on Patch-seq to establish the electrophysiological, neuroanatomical, and molecular diversity of Vglut2+ LHA-LHb neurons. In contrast to the notion of a homogenous LHA-LHb pathway, we found that the LHA-LHb pathway is composed of several neuron types characterized by discrete intrinsic properties, molecular markers, and spatial projection patterns. Cell-type specific optogenetic manipulations revealed that negative signals and persistent negative states are mediated through estrogen receptor 1-expressing (Esr1+) LHA-LHb neurons, whereas neuropeptide Y-expressing (Npy+) LHA-LHb neurons control rearing behavior. Neuropixels recordings in the PFC highlighted that induction of negative signals through repeated activation of Esr1+ LHA-LHb neurons leads to persistent shifts in the PFC neural activity, revealing a central role for the prelimbic area (PL) in encoding the negative state. Further supporting the role of Esr1+ LHA-LHb neurons in persistent negative emotional states, we found that increased stress sensitivity in female mice was dependent on the activity of Esr1+ LHA-LHb neurons and associated with a persistent shift in the intrinsic properties of the bursting-type Esr1+ LHA-LHb neurons. Altogether, the excitatory LHA-LHb pathway is composed of several discretely organized neuron subtypes with unique functions, where Esr1+ LHA-LHb neurons are central in mediating negative signals and stress sensitivity.

### Electrophysiological diversity of LHA-LHb neurons

To define the possible heterogeneity of LHA-LHb neurons, we visualized single LHA-LHb neurons and used ex vivo cell-attached and whole-cell patch-clamp recordings to probe the intrinsic properties of 230 LHA-LHb neurons in 46 mice (Fig. 1a and Extended Data Fig. 1a-d). First, we measured ex vivo the spontaneous activity in cell-attached recording mode and found that all recorded LHA-LHb neurons were spontaneously discharging action potentials (APs) (Fig. 1b). We validated the tonic firing activity of LHA-LHb neurons in vivo using Neuropixels recordings of optotagged LHA-LHb in head-fixed awake mice (Extended Data Fig. 1e-j). Following cell-attached recordings, we proceeded to ex vivo whole-cell recordings and characterized a large number of intrinsic parameters to classify putative neuron subtypes. We found that the recorded LHA-LHb neurons could be classified according their firing types (e.g. bursting, regular spiking, fast spiking). We therefore performed an expert-based classification and named the different neuron subtypes according to their defining intrinsic properties: a) fast adapting with putative Bk-current type (FA-Bk), b) bursting type (Burst), c) regular spiking with narrow APs type (RS-N), d) late-spiking with narrow APs type (LS-N), e) late-spiking with wide APs type (LS-W), f) regular spiking with wide APs type (RS-W) (Fig. 1c, Extended Data Fig. 1k-n and Table 1). The key intrinsic properties were segregated anatomically in the LHA suggesting topographical organization of the neuron subtypes (Fig. 1d-f). Supporting the anatomical organization of LHA-LHb neuron types, we found a discrete topographical organization of soma localization along the anteroposterior axis, as well as distinct somatic and dendritic morphologies for each subtype (Fig. 1e-h, Extended Data Fig. 2a-i, and Tables 2-3).

**Figure 1.**
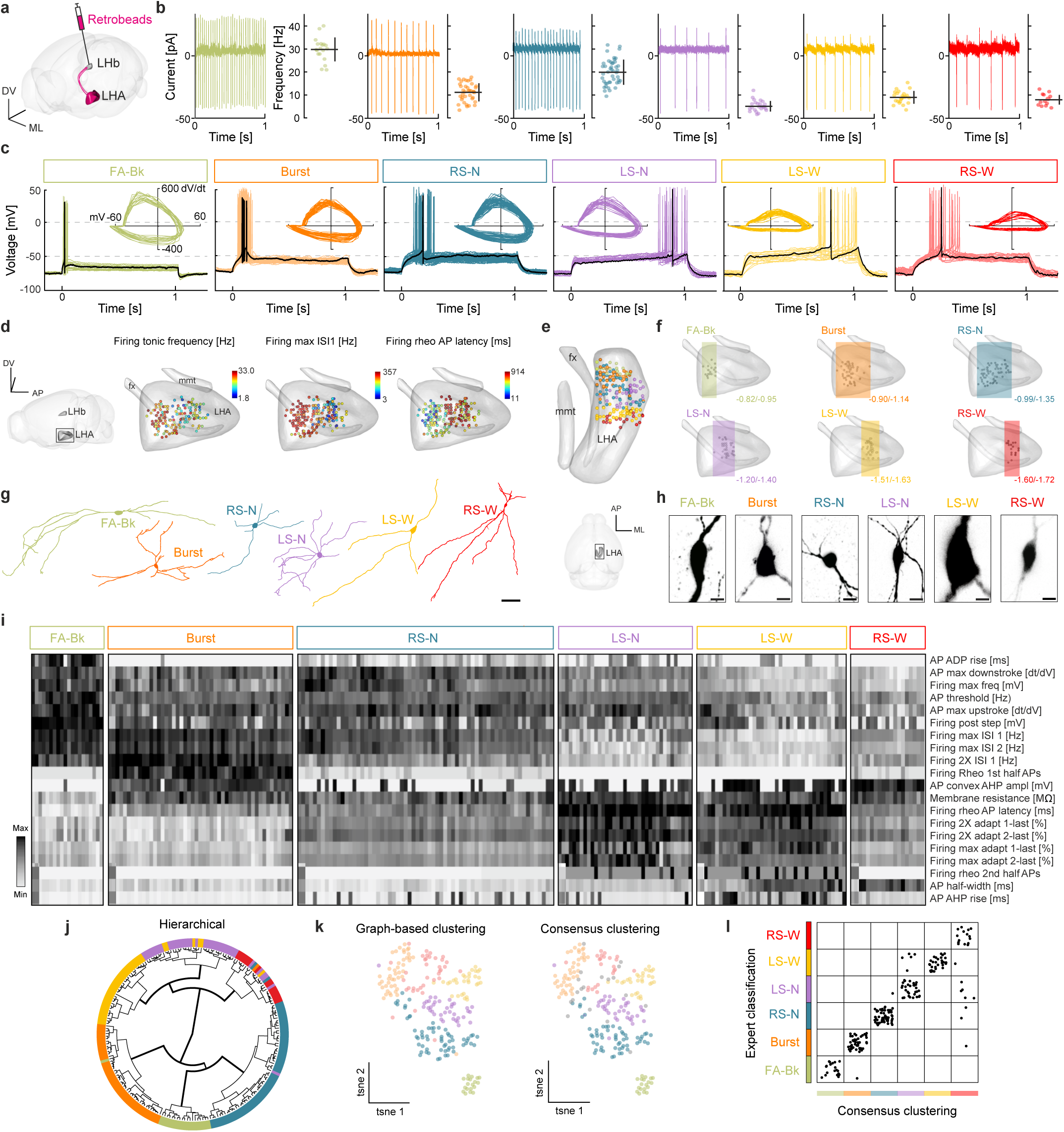
Electrophysiological diversity of LHA-LHb neurons. **a**, Strategy for retrograde labelling of LHA-LHb neurons. **b**, Cell-attached electrophysiological recordings reveal tonic firing of LHA-LHb neurons. Cell-attached traces from representative neurons (left), and the mean firing rate of the individual neurons (mean ± SD; right). n = 230 neurons, from left to right: n = 20, 49, 65, 37, 40, 19, N = 46 wt mice,). **c**, Overlay of the individual whole-cell traces of firing at rheobase (Rheo) of all recorded LHA-LHb neurons. Black: trace from a representative neuron. Inset: phase-plane plots of the first action potential at rheobase for all individual neurons. **d**, Electrophysiological properties reveal anatomical organization of LHA-LHb neurons. Dots: recorded LHA-LHb neurons color coded by different electrophysiological parameters. **e**, 3D position of all recorded LHA-LHb neurons (color coded by cell type). **f**, 3D visualization of the A-P distribution of electrophysiologically characterized LHA-LHb neuron types. Bregma coordinates show the most anterior and posterior coordinates for each subtype. **g**, Reconstruction of representative dendritic morphologies of LHA-LHb neuron types. **h**, Images of representative soma morphologies of the LHA-LHb neuron types. **i**, Heat map of 20 electrophysiological parameters selected based on principal component analysis (PCA). Classification of LHA-LHb neuron types by expert classification, as in (**c**). One column = one LHA-LHb neuron. **j**, Circular dendrogram for hierarchical clustering of LHA-LHb neurons. Color code: expert classification of LHA-LHb neurons. **k**, t-SNE plots of graph-based clustering (left), and consensus clustering (right). Color code: consensus clustering of LHA-LHb neurons. **l**, Agreement between the expert classification and the unsupervised consensus clustering of LHA-LHb neurons. Abbreviations: action potential (AP), anterior-posterior (A-P), medio-lateral (M-L), dorso-ventral (D-V), fast adapting with putative Bk-current (FA-Bk), bursting (Burst), regular spiking with narrow APs (RS-N), late-spiking with narrow APs (LS-N), late-spiking with wide APs (LS-W) and regular spiking with wide APs (RS-W), 2-times the rheobase current injection (2XRheo), maximal firing frequency (Max), principal component analysis (PCA) and membrane intrinsic property (Mem). All data acquired in male mice. Scale bar: 50 µm (**g**), 10 µm (**h**). **See also Extended Data Fig.1-2, Table 1-3.**

To validate the expert-based classification, we performed an independent clustering of the electrophysiological parameters using hierarchical clustering and graph-based clustering with 20 PCA-selected parameters from each neuron (Fig. 1i-k, Extended Data Fig. 1o). We found that the two unsupervised clustering methods resulted in a similar classification of LHA-LHb neurons into six clusters (Fig. 1j-k). To test the agreement between clustering methods, the two unsupervised approaches were first summed into a consensus clustering, which we then compared to the expert-based classification. We found that 214/230 neurons were classified into the same six neuron types independent of the approach used (Fig. 1l and Extended Data Fig. 1p-q). In summary, we uncovered a surprising diversity in the glutamatergic LHA-LHb pathway, establishing six discrete neuron types with characteristic tonic frequencies and intrinsic electrophysiological features.

### Molecular diversity of LHA-LHb neurons

We then asked whether the electrophysiological properties of the six LHA-LHb neuron types could also be captured by other modalities, for example their gene expression pattern. We therefore used Patch-seq (*21*) to obtain a combined view of intrinsic physiology, morphology, position, and gene expression pattern for the whole-cell-recorded neurons (Fig. 2a). We performed Patch-seq on 163 neurons (46 mice), resulting in ∼6200 detected genes per cell on average (Extended Data Fig. 2j-k). We used unbiased clustering of the single-cell RNA profiles to overlay the extracted molecular identity onto the expert-based electrophysiological classification (Fig. 2b). Clustering based on gene expression data alone was not sufficient to identify the putative six LHA-LHb types, and the three molecular clusters consisted of multiple electrophysiological types (Fig. 2c and Extended Data Fig. 2l). We therefore used the electrophysiological data to identify cell-type-specific markers for the six neuron types. We found that the LHA-LHb neurons as expected expressed the glutamatergic marker Vglut2 (gene name Slc17a6), and could be separated based on expression of a palette of molecular markers: FA-Bk neurons were Pv+/Nppc+; Burst neurons were Esr1+/Plpp4+; RS-N neurons were Esr1+/Glpr1+; LS-N neurons were Npy+/Pax6+; LS-W neurons were Gal+/Hcrt+; RS-W neurons were Avpr1a+/Hcrt+ (Fig. 2d). In conclusion, the multimodal classification of Vglut2+ LHA-LHb neurons based on electrophysiological, morphological, and molecular definitions, together established six discrete neuron subtypes.

**Figure 2.**
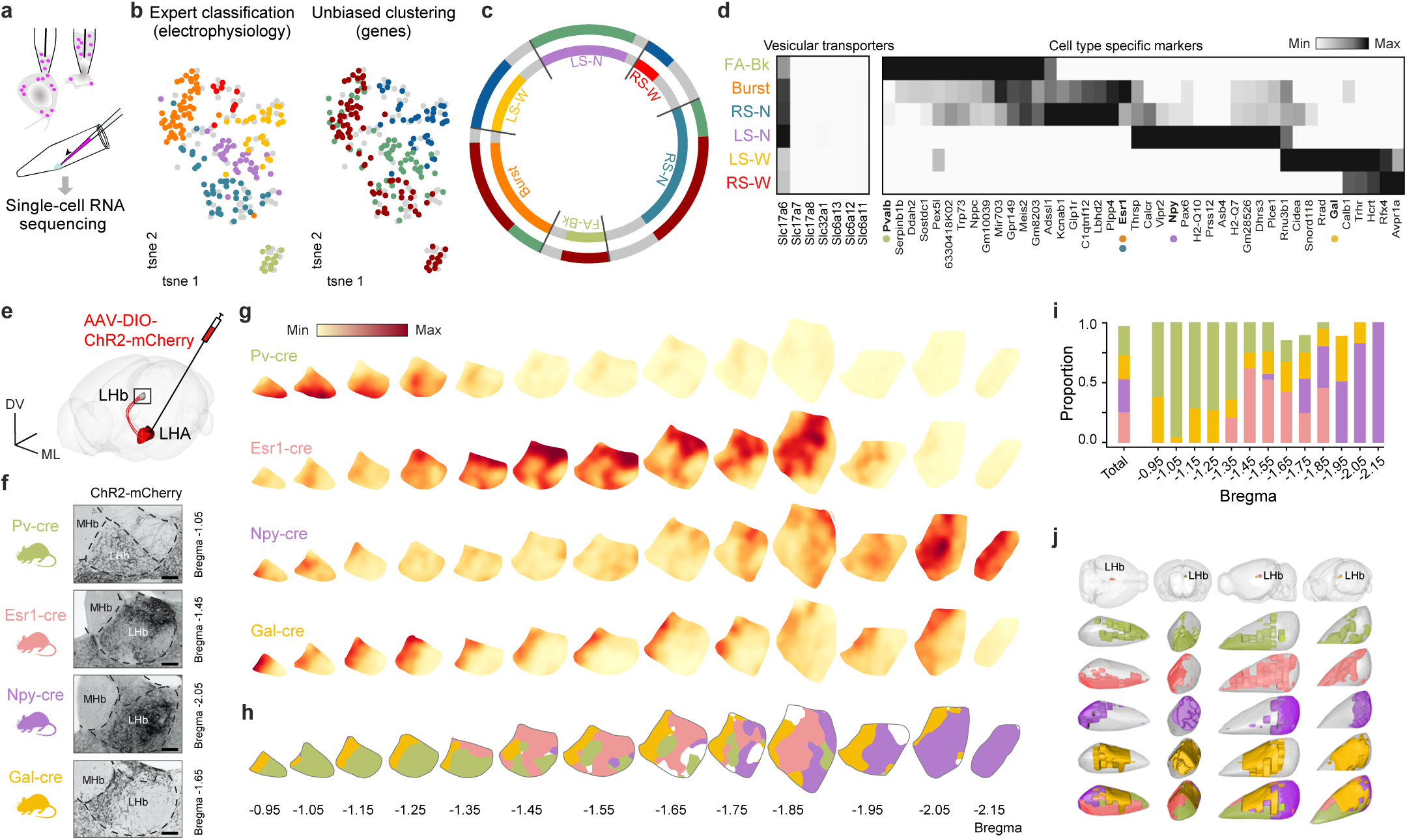
Discrete organization of genetically targeted LHA-LHb pathways. **a**, Schematic of Patch-seq showing somatic harvesting of retrobead-labeled LHA-LHb neurons. **b**, t-SNE plots of all recorded LHA-LHb neurons (n = 230 neurons, N = 46 wt mice, same neurons as in Fig. 1). The cell type identify of neurons harvested for Patch-seq are color coded based on electrophysiology (expert classification; left), or gene expression (unbiased clustering; right). n = 163 harvested neurons, FA-Bk: n = 12, Burst: n = 42, RS-N: n = 35, LS-N: n = 35, LS-W: n = 28, RS-W: n = 11, N = 46 wt mice. Gray neurons: recorded but not harvested. **c**, Comparison of electrophysiological (expert classification as in (**b**), left) versus gene expression classification (unbiased clustering as in (**b**), right) of LHA-LHb neurons (colors as in (**b**)). **d**, Heat-map of genes with differential expression in the electrophysiologically defined LHA-LHb neuron types (right). Expression of vesicular transporters in the LHA-LHb neuron types (left). Colored dots: genetic markers employed for subsequent cell-type specific targeting. **e**, Experimental strategy for anterograde labelling of LHA-LHb axon terminals. **f**, Representative images of virally labelled LHA-LHb axon terminals in the mouse cre lines used for targeting of specific LHA-LHb pathways (PV-, Esr1-, Npy-, and Gal-cre mice, respectively). Brain section with peak terminal density is shown. **g**, Heatmaps of the axon terminal density in the LHb for the four genetically targeted LHA-LHb pathways. **h**, Visualization of the topographical organization of the pathway-specific projection fields in the LHb. Colors as in g, white: not assigned to a specific pathway (Methods). **i**, Proportion of the LHb area targeted by the distinct LHA-LHb pathways, plotted along the A-P axis. Left bar; cumulative targeting of LHb by the four LHA-LHb pathways. **j**, 3D reconstructions (four different orientations) of the LHb projection fields of the four LHA-LHb pathways. Abbreviations: Medial habenula (MHb). All data acquired in male mice. Scale bar: 100 µm (**f**). **See also Extended Data 3-4.**

### Genetic targeting of LHA-LHb neuron types and neuroanatomical organization

Based on the molecular profile of the LHA-LHb neuron types revealed by Patch-seq, we used mice with cell-type specific cre recombinase expression to target the putative neuron types. We used Pv-cre, Esr1-cre, Npy-cre, and Gal-cre mice to target the main LHA-LHb neuron types, and injected cre-dependent adeno-associated viruses (AAV-DIO) into the LHA. We found that this strategy resulted in the genetic labeling of the distinct neuron types and their projections to the LHb (Fig. 2e-f and Extended Data Fig. 3). To validate the cell-type specific labeling strategy, we first established the specificity of our genetic targeting in Esr1-cre mice by confirming the presence of Esr1 protein in LHA-LHb neurons after cre-dependent viral labeling (Extended Data Fig. 4a-b). In addition, we used whole-cell recordings to confirm that the retrograde cell-type specific viral targeting approach in Esr1-cre mice labeled specifically the Burst and the RS-N neuron types, in Npy-cre mice LS-N neurons types, as well as in Pv-cre mice specifically the FA-BK LHA neuron type (Extended Data Fig. 4c). We also confirmed that retrograde targeting in Esr1-cre mice did not label neighboring Esr1+ nuclei (e.g. VMH), and that axons from Npy+ and Gal+ LHA-LHb neurons expressed the corresponding peptide (Extended Data Fig. 4d-g).

Taking advantage of the cell-type specific labeling of the different LHA-LHb neuron types, we aimed to further determine the topographical organization of neurons types and map their projection pattern. Using two different retrograde viral labeling approaches, we mapped the number and location of genetically labeled LHA-LHb subtypes. In summary, we confirmed that LHA-LHb subtypes were discretely organized in the anteroposterior axis, and that the Esr1+ and Gal+ subtypes were comparable in population size, whereas Pv+ and Npy+ subtypes were a smaller population (Extended Data Fig. 3d-l). Importantly, we found a specific organization of axon terminals for each LHA-LHb pathway, where axon terminals from Pv+ LHA-LHb neurons showed preferential targeting of anterior LHb domains, whereas the Esr1+ LHA-LHb neurons targeted the intermediate LHb domain with inputs to specific LHb subregions (e.g. avoiding the oval-LHbLO and the parvocellular-LHbMPc subnucleus of the LHb; Fig. 2f-h and Extended Data Fig. 3b-c). In contrast, the Npy+ LHA-LHb neurons targeted the posterior LHb domains, whereas Gal+ LHA-LHb neurons targeted a medial LHb domain along the entire anteroposterior axis. Importantly, the different LHA-LHb pathways targeted comparable cumulative areas but that were organized topographically across the LHb (Fig. 2i-j). Together, the cell-type specific targeting of the LHA-LHb neuron subtypes showed that the molecular and physiological profile also reflected a specialized neuroanatomical organization.

### Specialized role of LHA-LHb neuron types in behavior

To address whether the observed heterogeneity in the physiology and organization of LHA-LHb neuron types reflects functional specialization, we applied different strategies to first probe the contribution of each pathway to aversion. Based on candidate markers from the Patch-seq data, we used Pv-cre, Esr1-cre, Npy-cre, Gal-cre mice to target and manipulate four pathways independently, and compared these to manipulation of the entire LHA-LHb projection using Vglut2-cre mice. We used two different viral strategies to achieve cell-type specific bilateral optogenetic activation of cell bodies versus terminals: a) retrograde labeling using AAVretro-DIO-ChR2-mCherry injection into the LHb with somatic stimulation in LHA, and b) AAV-DIO-ChR2-mCherry injection into to the LHA in combination with axon terminal stimulation (Fig. 3a-c and Extended Data Fig. 5). To specifically probe the distinct role of Esr1+ LHA-LHb neurons, we further established a strategy for cell-type specific silencing of Esr1+ neurons using expression of the tetanus light chain (TeLC), which we also combined with optogenetic activation of the entire LHA-LHb pathway (Fig. 3a, d, and Extended Data Fig. 6a-r).

**Figure 3.**
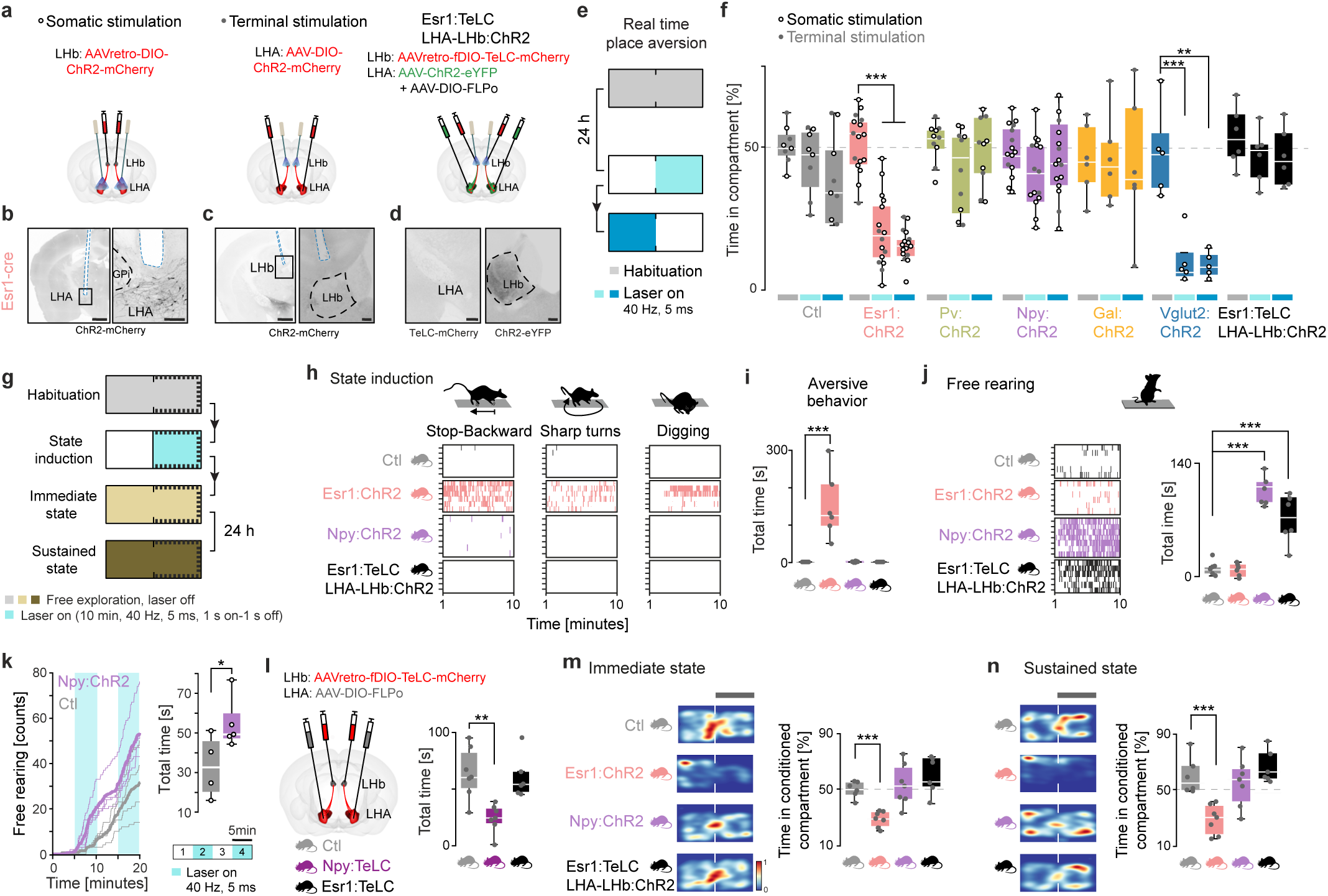
The Esr1+ LHA-LHb pathway drives aversion. **a**, Experimental strategy for somatic optogenetic manipulation of LHA-LHb neurons, optogenetic manipulation of LHA-LHb axon terminals in the LHb (middle), and optogenetic manipulation of the axon terminals in the LHb of the entire LHA-LHb pathway in combination with TeLC silencing of the Esr1+ LHA-LHb neurons (right). **b**-**d**, Representative images of the optic fiber path and/or viral expression for the three experimental strategies. b, ChR2-mCherry expression and somatic optogenetic manipulation of Esr1+ LHA-LHb neurons. Black box: location of right panel. c, ChR2-mCherry expression and optogenetic stimulation of Esr1+ LHA-LHb axon terminals in the LHb. Black box: location of right panel. d, Nuclear TeLC-mCherry expression in Esr1+ LHA-LHb neurons in the LHA (right), and ChR2-eYFP expression in LHA-LHb terminals in the LHb (left). **e**, Schematic outline of the real-time place aversion test (rtPA). 10 min habituation was 24 h later followed by optogenetic stimulation paired to the right compartment (ON, light blue, 10 min) and immediate switch of the optogenetic stimulation to the left compartment (ON SWITCH, blue, 10 min). **f**, Behavior in the rtPA. Optogenetic activation of the Esr1+ LHA-LHb and the VGlut2+LHA-LHb pathway, respectively, significantly reduced the time spent in the compartment paired with optogenetic stimulation (Esr1:ChR2 habituation vs ON or ON SWITCH: p < 0.001; Vglut2:ChR2 habituation vs ON: p < 0.001, habituation vs ON SWITCH: p = 0.0025, unpaired *t*-test). Horizontal bars: light blue, ON; blue, ON SWITCH; gray, right compartment during habitation. Empty circles: somatic stimulation, gray dots: axon terminal stimulation. Ctl: N = 8, Esr1:ChR2: N = 16, PV:ChR2: N = 10, Npy:ChR2: N = 15, Gal:ChR2: N = 6, Vglut2:ChR2: N = 5, Esr1:TeLC LHA-LHb: ChR2: N = 6. **g**, Schematic outline of the strategy for induction of an immediate (beige) and sustained aversive state (brown), respectively. State induction (light blue): 10 min optogenetic stimulation of LHA-LHb axon terminals in the LHb. **h**, Scoring of aversive behaviors (stop-backward movement; sharp turns; digging) during state induction, 1 colored coded vertical bar = 1 s in a specific aversive behavior. Ctl, Esr1:ChR2, and Esr1:TeLC LHA-LHb:ChR2 mice: N = 6 mice, respectively, Npy:ChR2: N = 7 mice, same animals in (**i**-**j**, **m**-**n**). **i**, The optogenetic stimulation of Esr1+ LHA-LHb axon terminals significantly increased the total time spent in aversive behaviors during state induction (Esr1:ChR2 vs Ctl: p = 0,0022, unpaired *t*-test). **j**, Scoring of free rearing during state induction, 1 colored coded vertical bar = 1 s free rearing (left). Optogenetic stimulation of Npy+ LHA-LHb axon terminals significantly increased the time spent free rearing (Npy:ChR2 vs Ctl : p < 0.001, unpaired *t*-test). The same effect was seen in repones to axon terminal stimulation of the compound LHA-LHb pathway in combination with TeLC silencing of the Esr1+ LHA-LHb neurons (Esr1:TeLC LHA-LHb:ChR2 vs Ctl: p < 0.001, unpaired t-test; right). **k**, Free rearing in the open field. Somatic activation of Npy+ LHA-LHb neurons significantly increased the total time spent free rearing (right; Npy:ChR2: (N = 5) vs Ctl (N = 4) mice: p = 0.0487, unpaired t-test). Cumulative number of free rearing events; thick line: mean (left). **l**, Experimental strategy for TeLC mediated silencing of Npy+ LHA-LHb and Esr1+ LHA-LHb neurons, respectively (left). TeLC silencing of Npy+ LHA-LHb neurons significantly decreased the total time spent free rearing in the open field (right; Npy:TeLC (N = 7) vs Ctl (N = 7): p = 0.0016, unpaired t-test). Esr1:TeLC: N = 6 mice. **m**-**n**, Free exploration directly after (j) and 24 h after (k) the state induction. Optogenetic stimulation of Esr1+ LHA-LHb axon terminals significantly increased conditioned place aversion (right; immediate state: Esr1:ChR2 vs Ctl mice: p < 0.001; sustained state: Esr1:ChR2 vs Ctl mice: p < 0.001, unpaired t-test). Representative heatmaps of locomotion (left). All data acquired in male mice. For boxplots (**f**, **i-n**), data shown as median, box (25th and 75th percentiles), and whiskers (data points that are not outliers). Scale bars: 1 mm, boxes 100 µm (**b**-**d**). \**p* < 0.05, \*\**p* < 0.01, \*\*\**p* < 0.001. **See also Extended Data 5-6, Video 1 and 2, and Table 4.**

To first evaluate aversion signaling, we used optogenetic activation in a two-compartment real-time place aversion task, and confirming previous findings (*6*), we found that somatic and terminal optogenetic activation of the entire glutamatergic (i.e.Vglut2+) LHA-LHb population mice induced significant avoidance of the opto-stimulated compartment. Importantly, somatic or axon terminal optogenetic stimulation of Esr1+ LHA-LHb neurons induced significant avoidance of the opto-stimulated compartment, whereas activation of Pv+, or Gal+, or Npy+ neuron subtypes failed to induce any avoidance response (Fig. 3e-f and Extended Data Fig. 6n, y, Video 1 and Table 4). To determine whether activation of Esr1+ LHA-LHb neurons is necessary for the aversive response, we performed the real-time place aversion task using optogenetic activation of all LHA-LHb neurons in combination with TeLC silencing of Esr1+ LHA-LHb neurons. We found that mice with silenced Esr1+ LHA-LHb neurons showed no aversion during optogenetic activation of the remaining LHA-LHb pathway (Fig. 3f), demonstrating their necessary role in mediating the aversive signals in the LHA-LHb pathway. Interestingly, although Pv+, or Gal+, or Npy+ neuron subtypes failed to induce real-time place aversion, we found that specifically the activation of Npy+ LHA-LHb neurons induced significant free rearing behavior in the open field (Extended Data Fig. 6s-x, Video 2). In addition, we explored the possible role of Pv+, or Gal+, or Npy+ in appetitive behaviors, and performed optogenetic activation during sucrose consumption. We did not observe any significant effects on sucrose consumption after activation of the Pv+, Npy+, or Gal+ LHA-LHb neurons arguing against a role in reward or food consumption (Extended Data Fig. 6z-aa).

Since we observed distinct and significant behavioral modulation by optogenetic activation of the Esr1+ and Npy+ LHA-LHb neurons, we proceeded to investigate their role in inducing persistent behavioral states. We established a state induction protocol based on optogenetic activation of each LHA-LHb pathway for 10 min followed by testing of the immediate and persistent behavioral response (Fig. 3g). We found that continuous activation of the Esr1+ LHA-LHb neurons induced specific behaviors associated with aversion (e.g. stop and backwards movement, digging behavior) during the stimulation period (Fig. 3h-i). In contrast, we did not observe any aversive behaviors even during continuous activation of the Npy+ LHA-LHb neurons (Fig. 3h-i). Importantly, continuous as well as block wise activation of Npy+ LHA-LHb neurons induced free rearing behavior during the entire stimulation period (Fig. 3j-k). The increased rearing was not associated with an increased anxiety-like state or induction of long-term negative signals, since optogenetic activation of Npy+ LHA-LHb pathway did not affect time spent in the center of the open field nor did it result in avoidance in the conditioned place aversion assay (Extended Data Fig. 6r, y). In summary, we found that activation of Npy+ LHA-LHb neurons can induce rearing, and that cell-type specific silencing of their activity significantly reduced free rearing under naturalistic conditions (Fig. 3l and Extended Data Fig. 6ab).

To directly address whether the continuous activation of Esr1+ or Npy+ LHA-LHb neurons could induce a persistent negative signal, we tested the immediate and sustained (24 hours later) behavioral response after the optogenetic state induction protocol (10 min). We found significant conditioned place aversion immediately after the activation of the Esr1+ LHA-LHb neurons and not after activation of Npy+ LHA-LHb neurons (Fig. 3m). Supporting a central role for Esr1+ LHA-LHb neurons in generating a sustained aversive state, we found that the conditioned place aversion persisted for 24 hours (Fig. 3n). Overall, these results show the central role of Esr1+ LHA-LHb neurons in mediating immediate aversive signals and promoting the persistent aversive state.

### Pathway-specific aversive and state signals in the PFC

We next asked whether we could identify shifts in brain-wide neural dynamics associated with the persistent aversive state induced by the Esr1+ LHA-LHb pathway. We focused our study on recording the PFC neural dynamics, based on the key role of the PFC in controlling emotional state through the integration of cortical and subcortical information (*22, 23*). Our aim was to identify brain-wide network effects linked to the aversive state, rather than direct synaptic modulation upon LHA-LHb optogenetic modulation, since the PFC is not known to have monosynaptic inputs from LHb (*24*). We used high-density Neuropixels recordings in head-fixed Esr1-cre and Vglut2-cre mice to map mPFC dynamics and behavioral responses to internally generated (i.e pathway-specific optogenetic activation) and externally derived (i.e. air puffs) aversive signals (Fig. 4a-b). To validate the aversive component of the PFC neural dynamics we performed the same experiments in Npy-cre and control mice. In block 1 (50 trials), we recorded responses to a pure tone (200 ms, 10 kHz), and in block 2 (100 trials) the response to the same tone followed by optogenetic stimulation (500 ms, 40 Hz pulse train) of the respective LHA-LHb pathway. In block 3 (50 trials) we recorded responses to the tone without optogenetic activation (same as in block 1), while we introduced in block 4 (50 trials) a new auditory stimulus (200 ms blue noise) followed by a mild air puff to the eye (Fig. 4b and Extended data Video 3).

**Figure 4.**
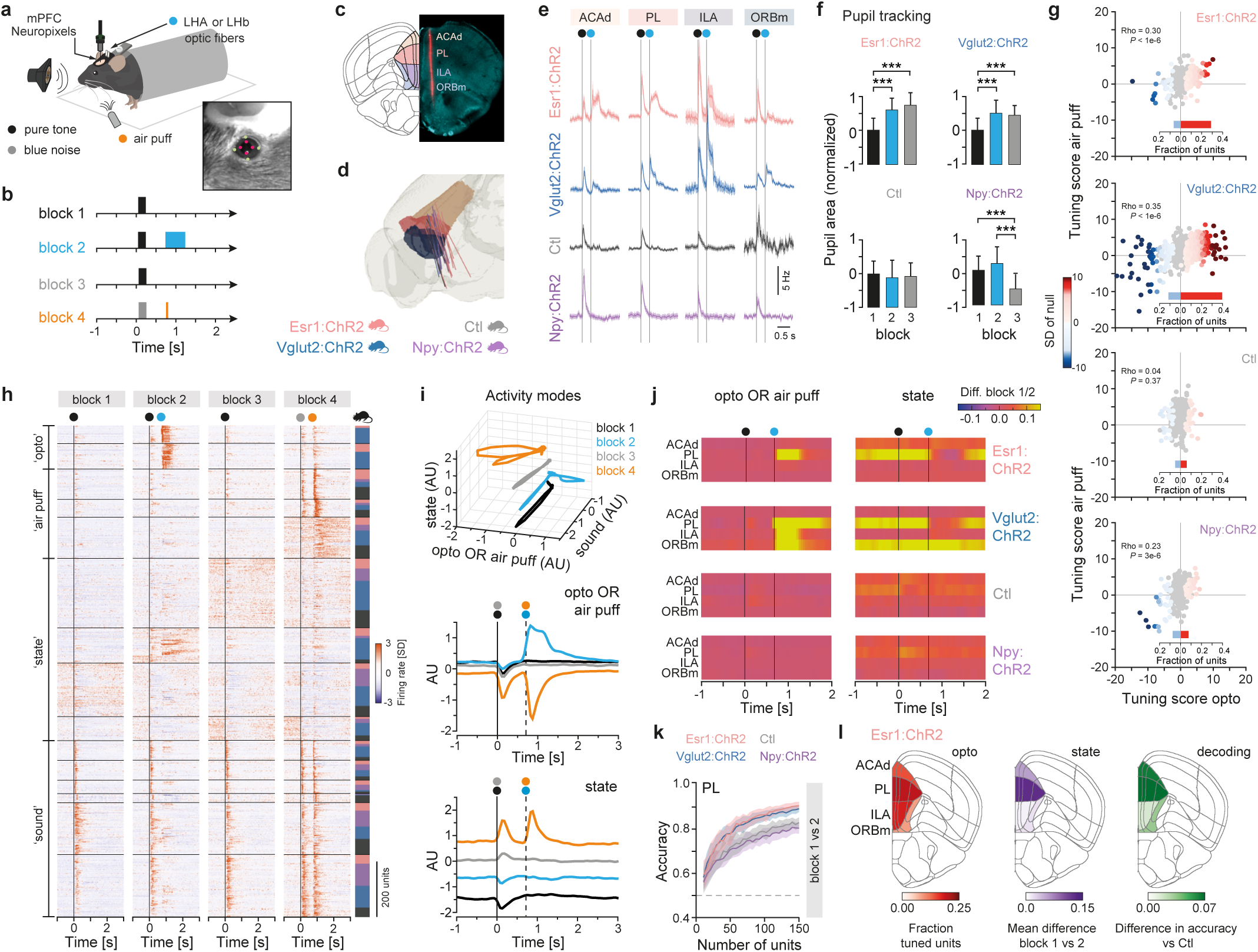
The Esr1+ LHA-LHb pathway shapes aversive signals and activity states in the prefrontal cortex. **a**, Schematic illustration of the experimental setup. **b**, Behavioral protocol, colors as in (**a**). Block 1: presentation of a pure tone (200 ms) at trial start. Block 2: presentation of a pure tone (200 ms) at trial start followed by optogenetic stimulation (500 ms) of LHA-LHb neurons 500 ms after end of the tone. Block 3: identical to block 1. Block 4: presentation of blue noise (200 ms) at trial start followed by mild air puffs (50 ms) to the eye 500 ms after end of sound presentation. **c**, Representative example of CM-DiI labeled (red) probe track in DAPI stained (turquoise) brain section. AP = 1.90 mm, Allen reference atlas CCFv3. **d**, 3D rendering of the tracked anatomical position of 25 Neuropixels probes in the PFC. Brain regions color coded as in (c). Bottom: the animal cohorts: Ctl, Npy:ChR2, and Vglut2-:ChR2 mice: N = 5, respectively, Esr1:ChR2: N = 10 mice for all panels. **e**, Peri-stimulus time histograms (PSTH; bin size 10 ms) of the firing rate modulation in block 2 for all units (n = 4135 neurons). Vertical lines: onset of auditory stimulus (black dot), and optogenetic stimulation (blue dot). Mean ± s.e.m **f**, Difference between the mean pupil area in single trails (block 1-3), and the mean pupil area in block 1 (block 1: n = 50 trials, 2: n = 100 trials, 3: n = 50 trials). Mean ± SD (Mann-Whitney U test with Bonferroni correction). **g**, Color coded tuning score for all units (dots; n = 1945 units) in response to optogenetic stimulation vs air puffs. Gray units: non-significant tuning. Bar graphs: fraction of units with significantly negative (light blue) and positive (red) tuning. **h**, Activity-based hierarchical clustering of all units (n = 1788 neurons; mouse cre lines color coded as in (**d**); vertical lines: onset of auditory stimulus (black or gray), optogenetic stimulation (blue), and air puffs (orange)). **i**, mPFC neuronal population activity in the four blocks projected in 3D onto three activity modes (‘state’, ‘sound’, and ‘opto OR air puff’; N = 25 mice, all genotypes; top). mPFC population activity projected onto the ‘opto OR air puff’’ mode (middle), and the ‘state’ mode (bottom). Vertical lines: onset of auditory stimulus (black or gray), optogenetic stimulation (blue), and air puffs (orange)). **j**, Mapping of the ‘opto OR air puff’ mode and the ‘state’ mode in discrete mPFC subregions. The difference between the projection of block1 vs block 2 shown as heatmap for each mPFC subregion. Vertical lines: onset of auditory stimulus (black) and optogenetic stimulation (blue). **k**, Decoding (average prediction accuracy) of block identity (block 1 vs 2) (mean ± SD, 50 repeated cross-validations; dashed line: 50% chance performance). **l**, Fraction of significantly tuned units across the mPFC subregions in response to optogenetic stimulation in Esr1:ChR2 mice (N = 10; left). Mean difference of the projection of the ‘state mode’ between block 1 vs block 2 across the mPFC subregions (middle). Difference in decoding (average prediction accuracy) of block identity (block 1 vs block 2) in Esr1:ChR2 vs Ctl mice (right). AP = 1.90 mm, Allen reference atlas CCFv3. Schematic in **a** adapted from scidraw (https://scidraw.io/). All data acquired in male mice. \*\*\**p* < 0.001. **See also Extended Data 7-9, Video 3, Video 4, Tables 5-9.**

We focused our analysis on recorded single units in four mPFC subregions – the anterior cingulate area, dorsal part (ACAd), the prelimbic area (PL), the infralimbic area (ILA), and the orbitofrontal area, medial part (ORBm) (Fig. 4c-d), resulting in 4135 well isolated units from 25 mice (Extended Data Fig. 7a). We found strong modulation of mPFC units in response to optogenetic activation only in Vglut2-cre and Esr1-cre and not in Npy-cre mice, while air puffs and auditory stimuli modulated mPFC activity in all four mouse groups (Fig. 4e and Extended Data Fig. 7a). The latency of the mPFC responses to auditory stimulus and air puff were similar across the different mice (Extended Data Fig. 7b-d). Importantly, similar to the aversive state induced in freely behaving mice, we found that head-fixed mice with activation of the Esr1+ or Vglut2+ LHA-LHb pathways developed a persistent behavioral state in block 2 and block 3 measured by a significant increase in pupil size (Fig. 4f and Extended Data Fig. 7e-h, Video 4, and Table 5).

To quantify the tuning of single mPFC units, we used a generalized linear model (GLM) to fit the unit firing rate (binned) with the events in the four blocks (two auditory stimuli, optogenetic activation, air puff,) as regressors (*25*). Tuning scores were computed for all recorded units fitted with the GLM (n = 1954 units) and significantly tuned units were identified (Extended Data Fig. 8a-e). This revealed a significant mean tuning score to the optogenetic activation of the Esr1+ and Vglut2+ LHA-LHb pathways (Fig. 4g and Extended Data Fig. 8f-g and Table 6), and spike waveform classification showed significant tuning of both narrow-spiking putative inhibitory interneurons (NS; n = 564), and wide-spiking putative pyramidal neurons units (WS; n = 3418) (Extended Data Fig. 8h-m, and Table 7-8). We found that opto-modulated mPFC units in block 2 trials showed a number of conditioned units (*26*) in Vglut2-cre and Esr1-cre mice (Extended Data Fig. 8n-q).

We next analyzed how aversive signals (internal optogenetic vs external air puff) shifted the mPFC population dynamics. We used principal component analysis (PCA) to visualize how neural activity evolved over the four blocks. We found similar activity trajectories during blocks with auditory stimuli (block 1 and 3) and in the block with air puffs across all four mouse groups (block 4; Extended Data Fig. 9a). In contrast, optogenetic drive of the Esr1+ and Vglut2+ LHA-LHb pathways resulted in dramatically divergent trajectories compared to Npy-Cre and control mice (i.e. in block 2), indicating prominent modulation of neural activities across many units and mPFC subregions. To better understand the contribution of single units to the divergent population dynamics, we performed hierarchical clustering of the activity profile of all GLM-fitted mPFC units (n = 1788 units). This unbiased classification identified two distinct clusters containing opto-modulated units (‘opto’ clusters), showing time-locked or sustained opto-modulation, and these units belonged to Vglut2-cre and Esr1-cre mice (block 2; Fig. 4h). Interestingly, we observed five clusters (‘state’ clusters) with units showing a significant increase in baseline firing rate in one block or ramping baseline activity across blocks (Extended Data Fig. 9b-g and Table 9).

Modulation of the spontaneous baseline firing could be a reflection of block-specific population activities associated with distinct internal states (*27*). We therefore next adapted a neural activity mode decomposition analysis to compare the main activity modes (‘sound’, ‘opto OR air puff’, ‘opto AND air puff’, ‘state’) revealed by the clustering, across genotypes and mPFC subregions. For this analysis, the high-dimensional neuronal activity space (dimensions = the number of recorded neurons) was reduced to a low-dimensional space (3D or 1D) where the axes were designed to maximally separate the two task epochs used to define the activity modes. The population activity of specific blocks, genotypes, and/or mPFC subregions was projected onto the activity modes, which showed accurate representation of block-specific events (sound, optogenetic, and air puff related activity) by their respective modes (Fig. 4i and Extended Data Fig. 9h-m).

We found specific differences in the projected neural activity between block 1 and block 2 onto the activity modes for the ‘opto OR air puff’, for the ‘opto AND air puff’, and for the ‘state’. For example, we found that the modulation along the activity mode capturing the optogenetic effect (‘opto OR air puff’) was specific to activation of Esr1+ or Vglut2+ LHA-LHb neurons in block 2. Interestingly, the ‘state’ mode (baseline activity in block 4 vs block 1) captured distinct population activity modes in each of the four blocks, with complete separation of the block projections across time (Fig. 4i). When we analyzed the region distribution of these activity modes (‘opto OR air puff’; ‘state’), we found that they were detected primarily in the PL in Esr1-cre mice, whereas they were widely distributed in the mPFC in Vglut2-cre mice (Fig. 4j, Extended Data Fig. 9j-k). This supports the notion that the ‘state’ activity mode reflects block-specific population activities associated with distinct behavioral and internal aversive states in mPFC.

To further address the region-specific encoding of the behavioral aversive state in mPFC, we used a logistic regression to decode block identity from the baseline activity (activity during 2 s preceding auditory stimulus). We found that baseline activity in PL allowed for increased accuracy in decoding block 1 vs block 2 identity in Esr1-cre and Vglut2-cre mice (Fig. 4k). No differences in accuracy between the genotypes were found when repeating the decoding of block identity (block 1 vs 2, 1 vs 3, 1 vs 4) using the baseline activity of the other mPFC subregions (Extended Data Fig. 9n-o). It is likely that the observed increase in decoding accuracy (i.e. block 1 vs block 2) based on the baseline activity of PL units reflects the induction of a distinct neural activity state that reflects some aspects of the aversive behavioral state induced by activation of the Esr1+ LHA-LHb pathway. The opto-modulation, negative state signature, and region decoding, together reinforce that activation of Esr1+ LHA-LHb neurons has profound and specific effects on the unit and population activities in PL (Fig. 4l and Extended Data Fig. 9p-v), revealing a region-specific signature of the aversive state in mPFC.

### Sexually dimorphic stress state is encoded in Esr1+ LHA-LHb subtype neurons

The sexual dimorphism in Esr1+ circuits and sex-selective gene expression (*17, 18*) in combination with our finding on the aversive state induction prompted us to investigate the role of Esr1+ LHA-LHb neurons in mediating the sensitivity to aversive stimuli and to develop a sustained stress state. We found that Esr1+ LHA-LHb neurons were necessary for fear conditioning in male and female mice, however in the case of milder aversive stimuli (novel auditory stimulus, looming stimulus) we found that silencing Esr1+ LHA-LHb neurons primarily reduced aversive responses in female mice (Extended Data Fig. 10a-g). To follow up the sensitivity to mild aversive stimuli in female mice, we established a stress paradigm consisting of repeated exposure to unpredictable mild shocks to induce a persistent shift in the stress response, followed by a battery of behavioral tests performed to quantify sex-specific behavioral differences (Fig. 5a-b and Extended Data Fig. 10h). We synchronized the start of the shock protocol and the behavioral phenotyping in female mice based on cytological criteria (metestrus phase) to minimize possible effects of the cycling of sex hormones. We found that female mice developed a persistent stress state in response to the unpredictable stressors, including significantly increased marble burying, aversive behaviors in the looming test, and immobility in the forced swim test 24 hours after the last unpredictable shock exposure (Fig. 5c-d, Extended Data Fig. 10i-k, and Video 5). We found that cell-type specific silencing (TeLC) of Esr1+ LHA-LHb neurons reduced the negative impact of the unpredictable stressors on behavior in female mice (Fig. 5e). To generate a metric of the stress state, we combined data from the three behavioral tests to establish a stress index (SI). This quantification showed that only female mice developed a significant and persistent stress state, and that this state required the activity of Esr1+ LHA-LHb neurons as shown by the cell-type specific silencing (Fig. 5f).

**Figure 5.**
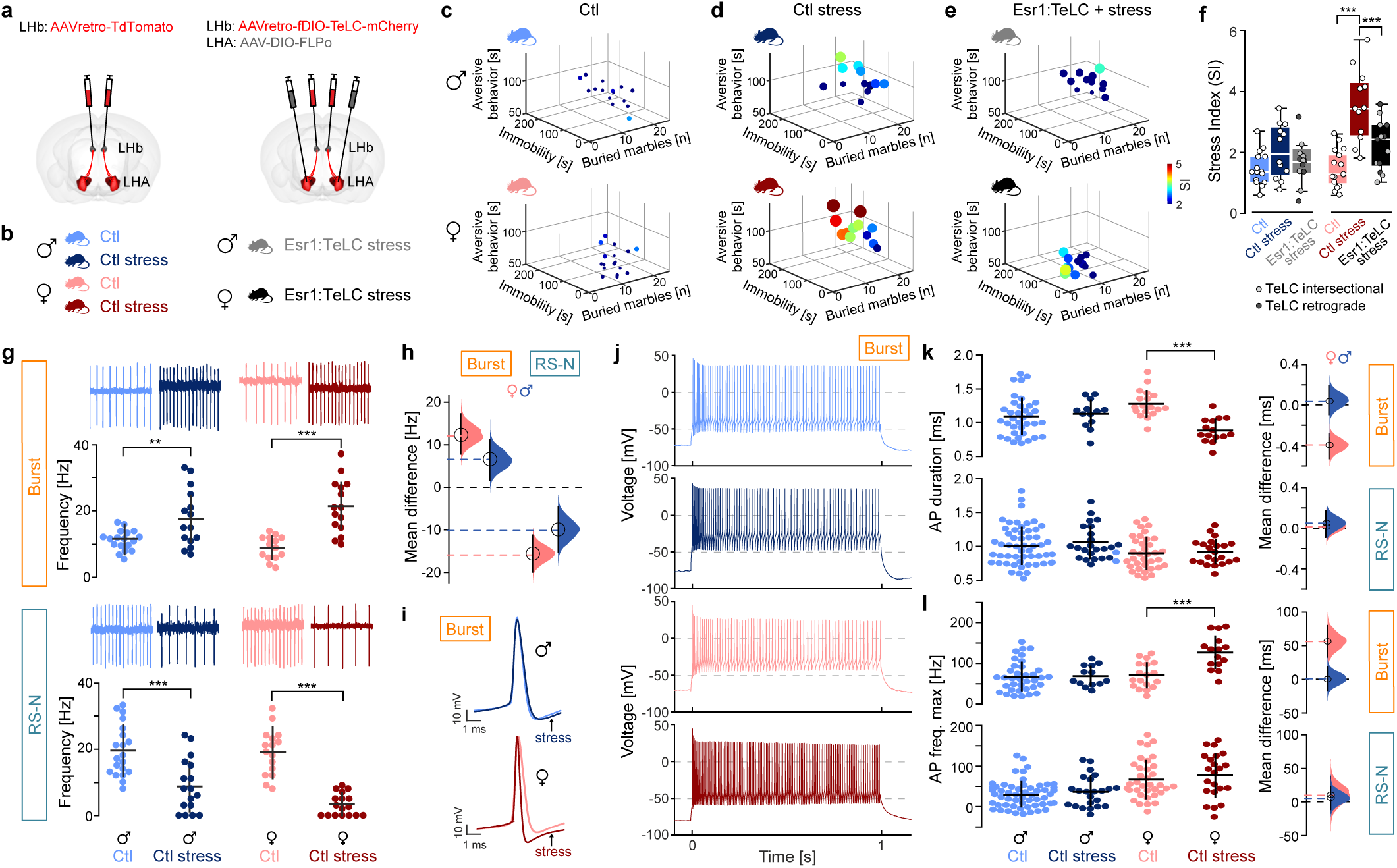
Sexually dimorphic stress sensitivity in Esr1+ LHA-LHb neurons. **a,** Experimental design for retrograde viral labelling of LHA-LHb neurons (left), and TeLC silencing of Esr1+ LHA-LHb neurons using intersectional strategy (right). See Extended Data 6c for alternative retrograde TeLC strategy. **b**, Animal cohorts used in behavioral phenotyping. Ctl mice (both sexes) were subjected to viral injection (a, left). Ctl stress mice (both sexes) were subjected to the same viral injection (a, left) and thereafter subjected to the stress paradigm. Mice (both sexes) with TeLC silencing of Esr1+ LHA-LHb neurons (Esr1:TeLC stress) were subjected to the stress paradigm. (e,f) shows combined data of TeLC silencing experiments with the intersectional (a, right) and with the retrograde strategy (Extended data 6c). **c-e**, Behavioral phenotyping 24 h after ending of the stress paradigm. 3D scatter plots summarizing: total time aversive behavior in the looming stimuli test (y axis), immobility in the FST (x axis), and marble burying (z axis). Data points: colored and sized by the stress index (SI). Ctl: N = 15 male, 16 female; Ctl stress: N = 12 male, 12 female; Esr1:TeLC stress: N = 12 male, 13 female. **f**, Quantification of the SI. The stress paradigm significantly increased the SI in female mice (Ctl vs Ctl stress female mice: p < 0.001, one-way ANOVA with Tukey’s Multiple comparisons test). Female mice with TeCL silencing of Esr1+ LHA-LHb neurons displayed significantly reduced SI (Ctl stress females vs Esr1:TeLC stress females: p < 0.001, one-way ANOVA with Tukey’s Multiple comparisons test). Same mice as in (**e**). **g-l**, Electrophysiological characterization of intrinsic properties of Esr1+ LHA-LHb neurons in stressed and control female and male mice (retrogradely labeled LHA-LHb neurons as in Extended Data 10l). **g**, Representative cell-attached traces of significantly increased tonic firing of Esr1+ Burst type neurons and decreased tonic firing of Esr1+ RS-N type neurons in stressed female and male mice (top). Quantification of the mean firing rate: Burst type neurons: Ctl stress female (n = 15) vs Ctl female (n = 14): p < 0.001, Ctl stress male (n = 15) vs Ctl male (n = 18): p = 0.0044; RS-N type neurons: Ctl stress female (n = 17) vs Ctl female (n = 16): p < 0.001, Ctl stress male (n = 17) vs Ctl male (n = 20): p < 0.001, DABEST test, mean ± SD; bottom. **h,** Quantification of the stress-induced change in the tonic firing frequency of Esr1+ Burst type (left) and Esr1+ RS-N type (right) neurons in female and male mice. The difference in firing between Ctl and Ctl stress females (pink) and Ctl and Ctl stress males (blue) is plotted. **i,** Representative rheobase APs of Burst type Esr1+ neurons. Quantified in k, colors as in g. **j,** Representative firing patterns of Burst type Esr1+ neurons at maximal depolarization. Quantified in l, colors as in (**g**). **k,** The AP duration of Esr1+ Burst type neurons is significantly decreased in stressed females. Quantification of the AP duration: Burst type neurons: Ctl stress female (n = 15) vs Ctl female (n = 18): p < 0.001, DABEST test, mean ± SD. RS-N type neurons: Ctl stress male: n = 22, Ctl male: n = 56, Ctl stress female: n = 22, Ctl female: n = 36. **l**, The maximal firing frequency of Esr1+ Burst type neurons is significantly increased in stressed females. Quantification of the maximal firing frequency: Ctl stress vs Ctl female mice: p < 0.001, DABEST test, mean ± SD, same neurons as i (**k**). n = number of neurons, N = number of mice. Data acquired in male vs female mice as stated in respective panel. For boxplots (**f**), data shown as median, box (25th and 75th percentiles), and whiskers (data points that are not outliers). Modified Gardner-Altman plots (**h, k-l**): distribution of bootstrap-sampled mean differences (area; pink: female Ctl stress vs female Ctl, blue: male Ctl stress vs male Ctl), mean of bootstrap sampling distribution of (circle) and the 95% confidence interval (CI; vertical black line). \*\**p* < 0.01, \*\*\**p* < 0.001. **See also Extended Data 10 and Extended Data Video 5.**

We then asked whether the Esr1+ Burst and the RS-N neuron types showed cell-type as well as sex-specific shifts in their intrinsic properties in response to the stress state. We followed the same stress induction protocol, exposing a separate cohort of male and synchronized female mice to the unpredictable mild shock protocol. To capture the sustained differences in intrinsic properties, we performed ex vivo cell-attached and whole-cell recordings from genetically labeled Esr1+ LHA-LHb neurons 24 hours after the last unpredictable shock as in the behavioral phenotyping (Extended Data Fig. 10h). We first confirmed that the electrophysiological parameters allowed us to classify Esr1+ LHA-LHb neuron types also in female mice under baseline conditions (Extended Data Fig. 10l-t and Table 10). We then recorded in the cell-attached mode to define the putative adaptation of the spontaneous firing frequencies in the stress state. Unexpectedly, we found that Burst and RS-N types were inversely affected by the stress state as Burst type neurons increased and RS-N type neurons decreased their firing in both sexes (Fig. 5g-h). The increased firing of the Burst neuron type supported their role as the primary candidate in mediating aversive stimuli. We also noticed that the increased firing of Burst type neurons was more prominent in female compared to male mice. We therefore next quantified the intrinsic properties from whole-cell recordings to determine whether the observed excitability adaptation could reflect the observed behavioral stress sensitivity in female mice. We found a significant decrease in the AP duration and a significant increase in the maximum firing frequency specifically in Burst type Esr1+ neurons from stressed female mice (Fig. 5i-k). Interestingly, we only found significant differences in the Burst type Esr1+ neurons in female mice, and no statistically significant changes in Burst type or RS-N type Esr1+ neurons in male mice. These results suggest that Esr1+ Burst type neurons, through shorter AP duration and increased maximum firing, became persistently more excitable upon induction of the sustained stress state in female mice. In summary, our findings show that the activity of Esr1+ LHA-LHb neurons is necessary for the sex-specific sensitivity to develop the stress state, and this state is associated with a specific shift in the intrinsic properties of Esr1+ LHA-LHb Burst type neurons in female mice.

## Discussion

We have uncovered an extensive diversity in the LHA-LHb pathway, establishing six neurons types with discrete physiology, molecular profile, and projection pattern. Using candidate molecular markers from Patch-seq data, we were able to visualize the cell-type specific organization and determine the role of LHA-LHb neuron subtypes in the control of immediate as well as sustained emotional behaviors. Our classification revealed a specialized organization of the glutamatergic LHA-LHb pathway and showed the discrete functional role for Esr1+ versus Npy+ LHA-LHb neuron subtypes. We found that the aversive signals induced by activation of the entire glutamatergic (Vglut2+) LHA-LHb pathway are mediated by the Esr1+ subpopulation, which are both necessary and sufficient to signal immediate negative signals as well as induce persistent states of aversion or stress. In contrast, we found that the Npy+ LHA-LHb pathway does not impact on avoidance or negative signals and instead controls rearing, since optogenetic activation of the Npy+ LHA-LHb pathway induced free rearing, and the cell-type specific silencing resulted in decreased rearing in the open field. Rearing is observed in naturalistic behaviors linked to exploration and has been proposed to signal novelty and risk assessment, for example associated with escape or defensive responses, and it is interesting to note that stimulation of escape or fear-related centers (e.g. PAG) can increase rearing behavior (*28, 29*). Npy+ LHA-LHb neurons mediate signals linked to exploration, although the relationship to uncertainty or escape-related states remains to be established.

The distinct behavioral effects we observed (i.e. aversion in Esr1+ vs. rearing in Npy+) could result from the topographically organized targeting of different LHb domains leading to the recruitment of divergent subcortical or cortical circuits. In combination with our findings on the role of LHA-LHb neuron subtypes in persistent aversive behavioral states, this prompted us to probe how internally generated negative signals though the LHA-LHb pathway indirectly shape brain-wide neural dynamics. The PFC is proposed to be a major hub for representing and integrating emotional signals, and we chose to therefore perform Neuropixels recordings in PFC in combination with optogenetic activation of the genetically distinct LHA-LHb pathways. We were able to generate a persistent negative state by optogenetic stimulation of the entire Vglut2+ LHA-LHb pathway in head-fixed mice, which was reflected in a strong modulation of the neural signal across several PFC subregions. In comparison, optogenetic activation of the Esr1+ LHA-LHb pathway induced a behavioral response of immediate as well as persistent aversion that primarily modulated the neural dynamics in the PL region. Our results support a model where negative emotional states are represented by specific ensembles in mPFC (*30, 31*), which are under the influence of specific LHA-LHb subpopulations.

These observations prompted us to further investigate a possible link between the Esr1+ LHA-LHb pathway and the behavioral manifestation of persistent negative states and stress. It is known that exposure to unpredictable negative events can precipitate a chronic stress response and shift the circuit dynamics into a maladaptive state (*32*). To test the role of Esr1+ LHA-LHb neuron subtypes in stress related behavior, we established a behavioral paradigm to specifically capture sex-specific differences in the sensitivity to develop a persistent stress-related state at the behavioral as well as the cellular level. We found that a paradigm based on repeated exposure to unpredictable mild foot shocks revealed a significantly increased sensitivity in female mice to develop behavioral correlates of stress. Importantly, the behavioral stress state was dependent on the activity of Esr1+ LHA-LHb neurons. While we have identified that Esr1-expression marks two electrophysiologically distinct LHA-LHb neuron types (i.e. the Burst and the RS-N type), we found that the stress state was associated with altered intrinsic properties specifically in the Burst-type Esr1+ neurons.

We found that Burst-type Esr1+ LHA-LHb neurons show persistent stress-induced shifts, suggesting that these play a role in mediating stress-related signals to downstream circuits. In a broader context, it is possible that Esr1+ neurons in several subcortical circuits can share a similar function in the regulation of behavioral states, and specifically through serving as nodes of sexually dimorphic gene expression and neuron function (*17, 18*). Further studies using intersectional and viral strategies based on the candidate markers we have presented, and that allow for differential targeting of Burst versus RS-N type Esr1+ LHA-LHb neurons, will be required to establish causal links between the cell-type specific state and the behavioral manifestation of stress.

It is interesting to note that negative emotional states, such as the chronic expression of stress or mood disorders, can emerge from similar patterns of persistent or burst-like activity in several nodes in the LHA-LHb-PFC circuitry. For example, persistent activity in hypothalamic neurons is found in fear and threat situations (*33*) and persistent activity in the PL region could underlie some of the sex differences in fear (*34*). Similarly, increased bursting activity in the LHb is a marker of depression-like behavior in animal models (*9, 35*). How negative signals in LHA-LHb produce acute versus persistent network effects in the mPFC that control the emotional state remains to be determined, and the role of region-specific signals needs to be further investigated, particularly since the functional identity of mPFC subregions is still debated (*36*).

Taken together, we propose that Ers1+ LHA-LHb neurons are central in signaling negative value, are necessary and sufficient to induce a persistent negative state, and mediate the sex-dependent sensitivity to develop behavioral and cellular maladaptive responses to unpredictable stressors. Our multimodal classification of the LHA-LHb pathway establishes a road map for understanding how diverse neuron types contribute to the complex dimensions of the emotional state and to sex differences in circuit function linked to stress.

## METHODS

### Animals

All procedures and experiments on animals were performed according to the guidelines of the Stockholm Municipal Committee (approval number N166/15, 155440-2020 and 7362-2019). Adult 2-5 months old mice were used: Esr1-cre (B6N.129S6(Cg)-Esr1^tm1.1(cre)And^/J; Jackson Laboratory Stock No: 017911), Npy-cre (B6.Cg-Npy^tm1(cre)Zman^/J; Jackson Laboratory Stock No: 027851), Pv-cre (B6;129P2-*Pvalb^tm1(cre)Arbr^*/J; Jackson Laboratory Stock No: 008069), Gal-cre (Tg(Gal-cre)KI87Gsat), Vglut2-cre (STOCK *Slc17a6^tm2(cre)Lowl^*/J; Jax Stock No: 016963) and wild-type mice (C57BL/6J; Charles River). Animals were group housed, up to five per cage, in a temperature (23°C) and humidity (55%) controlled environment in standard cages on a 12:12 hours light/dark cycle with *ad libitum* access to food and water, unless placed on a food restriction schedule. All food-restricted mice were restricted to 85–90% of their initial body weight by administering one feeding of 2.0–2.5 g of standard grain-based chow per day. All strains used were backcrossed with the C57BL/6J strain. Male mice were used for electrophysiological characterization of LHA-LHb neurons, Patch-seq, optogenetic manipulation and Neuropixels *in vivo* recordings (Figures 1-4). Once the cell types were electrophysiologically and molecularly established, as well as *in vivo* optogenetic manipulation and electrophysiological recordings where performed, we extended our analysis to mice of both sexes. Male and female mice, when possible aged matched and littermates, were used for baseline behavioral testing, and behavioral and electrophysiological characterization upon stress induction (Figure 5 and Extended data Figure 10).

### Estrous cycle staging identification

Four stages of the estrous cycle were determined by collection and analysis of predominant cell typology in vaginal smears as reported (*37*). Samples were collected daily throughout the duration of the experiment, consistently at the same time of the day (9-10 AM). In brief, the sample was collected with ddH2O-filled tip at the opening of the vaginal canal. The smear was placed on a glass slide and left at room temperature until completely dried. Cytological staining was performed by placing the dry slide in a coplin jar containing the crystal violet stain for 1 min, followed by repeated washes in ddH2O. The smear was examined under light microscopy to determine cell-types present. Once mice exhibited at least two regular four days estrous cycles, shock protocol was started on the day of metestrus, identified as small darkly stained leukocytes predominate. Electrophysiological recordings and behavioral phenotyping were performed on the metestrus day of the following cycle.

### Tracers and viral constructs

We used red retrobeads (Lumafluor) to visualize LHA-LHb neurons for the characterization of electrophysiological diversity in order to minimize potential cell type labeling bias due to viral tropism. Soma harvesting for Patch-seq of LHA-LHb neurons was performed after labelling with retrobeads in order to prevent potential effects of viral transduction on gene expression profile. Purified and concentrated adeno-associated viruses (AAV) were purchased from Addgene. Cre-inducible ChR2-mCherry was used (AAV-EF1α-DIO-hChR2(H134R)-mCherry; http://www.addgene.org/20297/) both for anterograde (Addgene viral prep #20297-AAV5) and for retrograde (Addgene viral prep #20297-AAVrg) cre-dependent optogenetic manipulation of LHA-LHb neurons. Control groups were injected with pAAV-hSyn-mCherry (a gift from Karl Deisseroth; Addgene viral prep # 114472-AAV5; Addgene viral prep # 114472-AAVrg; http://www.addgene.org/114472/). For TeLC silencing we used either a retrograde approach via cre-inducible TeLC (hSyn1-dlox-TeTxLC_2A_NLS_dTomato(rev)-dlox-WPRE-hGHp(A); VVF UZH ID # v620) or an intersectional approach combining a retrograde FLPo-dependent TeLC in LHb (ssAAV-retro/2-hSyn-fDIO-TeLC-P2A-dTomato, custom made at https://www.wzbio.com/) with a cre-dependent FLPo-expressing AAV5 in LHA (AAV5-pEF1a-DIO-FLPo-WPRE-hGHpA; Addgene viral pep # 87306-AAV5; https://www.addgene.org/87306/). To combine Esr1:TeLC silencing with optogenetic stimulation of the entire LHA-LHb pathway (Figure 3) we injected cre-independent ChR2 in LHA (AAV5-pAAV-hSyn-hChR2(H134R)-EYFP; Addgene viral pep # 26793-AAV5; https://www.addgene.org/26973/) together with the intersectional TeLC approach: we injected retrograde FLPo-dependent TeLC in LHb (ssAAV-retro/2-hSyn-fDIO-TeLC-P2A-dTomato, custom made at https://www.wzbio.com/) with a cre-dependent FLPo-expressing AAV5 in LHA (AAV5-pEF1a-DIO-FLPo-WPRE-hGHpA).

For electrophysiological validation of the TeLC silencing intersectional approach we co-injected retrograde FLPo-dependent TeLC (ssAAV-retro/2-hSyn-fDIO-TeLC-P2A-dTomato, custom made at https://www.wzbio.com/) and retrograde creON/flpON ChR in LHb (pAAV-hSyn Con/Fon hChR2(H134R)-EYFP; Addgene viral pep # 55645-AAVrg; https://www.addgene.org/55645/) and a cre-dependent FLPo-expressing AAV5 in LHA (AAV5-pEF1a-DIO-FLPo-WPRE-hGHpA; Addgene viral pep # 87306-AAV5; https://www.addgene.org/87306/). As positive control experiments electrophysiological validation of the TeLC silencing intersectional approach we injected retrograde creON/flpON ChR in LHb (pAAV-hSyn Con/Fon hChR2(H134R)-EYFP; Addgene viral pep # 55645-AAVrg; https://www.addgene.org/55645/) and a cre-dependent FLPo-expressing AAV5 in LHA (AAV5-pEF1a-DIO-FLPo-WPRE-hGHpA; Addgene viral pep # 87306-AAV5; https://www.addgene.org/87306/). As additional positive control we injected in LHA cre-inducible ChR (pAAV-Ef1a-DIO hChR2-EYFP; Addgene viral pep #35509-AAV5; https://www.addgene.org/35509/). For electrophysiological characterization of LHA-LHb neurons in Esr1-cre, Npy-cre and Pv-cre mice at baseline as well as for electrophysiological characterization of Esr1+ LHA-LHb neurons after stress we injected retrograde cre-dependent tdTomato (pAAV-FLEX-tdTomato; Addgene viral prep #28306-AAVrg; http://www.addgene.org/28306/) in the LHb. The helper virus TVA-V5-RG (AAV5-EF1a-DIO-TVA-V5-t2A-Rabies G) and Rabies-EGFP virus were cloned and produced in the Meletis laboratory.

### Stereotaxic injections

All stereotaxic injections were performed in 8-10 weeks old mice. For injections of red retrobeads (Lumafluor) in the LHb, wild-type mice were anesthetized with 5% isoflurane and mounted in a stereotaxic apparatus (Harvard Apparatus). 50 nL of retrobeads were injected at 100 nL/minute unilaterally using a glass micropipette with a Quintessential Stereotaxic Injector (Stoelting). The LHb was targeted using the following coordinates: 1.65 mm caudal and 0.35 mm lateral to bregma, and 2.55 mm deep from the dura. Mice were killed 4-7 days post-injection for slice electrophysiology or patch-seq. Viral retrograde labelling of cell-type specific projection neurons was achieved by injecting 150 nL of AAVretro-EF1α-DIO-hChR2(H134R)-mCherry or AAVretro-FLEX-tdTomato (for anatomical mapping, optogenetic manipulation and *in vivo* Neuropixels recording) or 150 nL of AAVretro-FLEX-tdTomato (for electrophysiological recording after stress induction protocol) or 150 nL of AAVretro-hSyn-fDIO-TeLC-P2A-dTomato or AAVretro-hSyn1-dlox-TeTxLC-2A-NLS-dTomato (for TeLC silencing) bilaterally at 50 nL/minute. The LHb was targeted using the same coordinates reported above for retrobead injections. For anterograde labelling of LHA neurons, 200 nL of AAV5-EF1α-DIO-hChR2(H134R)-mCherry or AAV5-FLEX-tdTomato or 200 nL of AAV5-DIO-FLPo was injected at 50 nL/minute in LHA unilaterally (for mapping input topography of the LHb) or bilateral (for behavioral experiments upon optogenetic manipulation or TeLC silencing). The LHA was targeted at the peak coordinates of each cell-type using the following coordinates: Pv-cre: 1 mm; Esr1-cre: 1.25 mm; Npy-cre: 1.45 mm; Gal-cre: 1.55 mm; Vglut2-cre and C57BL/6J: 1.4 mm caudal from bregma. LHA injections in all mice were performed 1 mm lateral from bregma, and 4.65mm deep from the dura. For cell-type projection-specific retrograde tracing using Rabies virus 100 uL of TVA-V5-RG (AAV5-EF1a-DIO-TVA-V5-t2A-Rabies G) was injected in the LHA and 2 weeks after 100 uL of Rabies-EGFP was injected into the LHb. For all stereotaxic injections, the glass pipette was left in place for 15 minutes after the injection and before removal from the brain. Optogenetic experiments were performed 15 days post-injection. TeLC experiments were performed 6 weeks post-injection. For anatomy, mice were killed 15-20 days post-injection. For Rabies retrograde tracing experiments, mice were killed 10 days after the Rabies virus injection.

### Implant surgery

Adult mice were anesthetized with isoflurane (3 % for induction then 1-2 %). Buprenorphine (0.1 mg/kg s.c.), carprofen (5 mg/Kg s.c.) and lidocaine (4 mg/kg s.c.) was administered. The body temperature was maintained at 37°C by a heating pad. An ocular ointment (Viscotears, Alcon) was applied over the eyes. The head of the mouse was fixed in a stereotaxic apparatus (Kopf). Lidocaine 2% was injected locally before skin incision. The skin overlying the cortex was removed, the skull was cleaned with Chlorhexidine and the bone gently cleaned. A thin layer of glue was applied on the exposed skull. A light-weight metal head-post was fixed with light curing dental adhesive (OptiBond FL, Kerr) and cement (Tetric EvoFlow, Invoclar Vivadent, Schaan, Liechtenstein). For optogenetics, two small craniotomies (∼300-500 µm in diameter) are performed to allow the insertion of the two fibers that are slowly lowered vertically into the brain at the depth for photoactivation (LHb: +1.55 mm AP; 0.95 mm ML from bregma and 2.2 mm depth with 10° angle from the dura, LHA: +1.1 mm ML; 4.25 mm depth from the dura; the AP coordinate was adjusted according to peak coordinates in each genotype: Pv-cre: 0.9 mm, Esr1-cre: 1.1 mm, Npy-cre: 1.2 mm, Vglut2-cre and C57BL/6J: 1.1 mm) and cemented to the skull. For extracellular recordings, a chamber was made by building a wall with dental cement along the coronal suture and the front of the skull. Brain regions were targeted using stereotaxic coordinates. After the surgery, the animal is returned to its home cage and carprofen (5 mg/Kg s.c.) was provided for postoperative pain relief 24 hours following surgery.

### Preparation of acute brain slices

All experiments were performed in 250 µm-thick coronal slices prepared on a VT1200S vibratome (Leica, Germany) in ice-cold artificial cerebrospinal fluid containing (in mM): 90 NMDG, 2.5 KCl, 1.25 Na_2_HPO_4_, 0.5 CaCl_2_, 8 MgSO_4_, 26 NaHCO_3_, 20 D-glucose, 10 4-(2-hydroxyethyl)-1-piperazineethanesulfonic acid (HEPES), 3 Na-pyruvate, 5 Na-ascorbate (pH 7.4). Brain slices were then incubated at 22-24 °C for 60 minutes in a recording solution containing (in mM): 124 NaCl, 2.5 KCl, 1.25 Na_2_HPO_4_, 2 CaCl_2_, 2 MgSO_4_, 26 NaHCO_3_, 10 D-glucose (pH 7.4). All constituents were from Sigma-Aldrich. Both solutions were aerated with carbogen (5% CO_2_/95% O_2_). Temperature in the recording chamber was set to 33 °C. Brain slices were superfused with the recording solution at a rate of 4-6 ml/min.

### Patch-clamp electrophysiology

Neurons were visualized by differential interference contrast (DIC) microscopy on an Olympus BX51WI microscope. Whole-cell recordings were carried out using borosilicate glass electrodes (Hilgenberg, Germany) of 5-8 MΩ pulled on a P-1000 instrument (Sutter). Electrodes were filled with an intracellular solution containing (in mM): 130 K-gluconate, 3 KCl, 4 ATP-Na_2_, 0.35 GTP-Na_2_, 8 phosphocreatine-Na_2_, 10 HEPES, 0.5 ethyleneglycol-bis(2-aminoethylether)-*N*,*N*,*N*′,*N*′-tetraacetate (EGTA), (pH 7.2 set with KOH), with Cl-reversal potential ∼-95 mV. Patch-clamp recordings were carried out on a Multiclamp 700B amplifier (Axon, USA) controlled by DigiData 1550B digitizer. Current clamp recordings were not corrected for −9.99 ± 0.38 mV liquid junction potential between the intracellular and recording solutions, as measured against a 3M KCl-electrode. Membrane input resistance (“Mem resistance” in MΩ) was calculated using linear regression established between electrotonic voltage responses to current injections (±20 pA from -70 mV holding) of 500 ms. Membrane time constant (t, ms) was averaged from 20 successive electrotonic voltage responses to hyperpolarizing (-40 pA) current steps and fitted with a single exponential. “AP threshold” (mV) was defined as the voltage point where the upstroke’s slope trajectory first reached 10 mV/ms. AP characteristics were measured on the rheobasic AP elicited by a 1-s-long current step. “AP rise time” (ms) was the time from the AP threshold to the AP’s peak. “Maximum AP up- and AP down-stroke” was determined as the maximum and minimum of the geometrical differential of the AP (mV/ms), respectively. “AP half-width” (ms) was measured at half the maximal amplitude of the AP. Afterpotentials were in certain neuronal types afterdepolarizations (ADP in mV) measured from the most negative fast repolarization of the AP to the most maximal voltage of the ADP. The time to reach this maximal ADP amplitude defined the “ADP rise time” (ms). Afterpotentials could also had been afterhyperpolarizations (AHP in mV) which could either have simple “convex” shape without ADP (AP convex AHP amplitude (mV)) or more complex “concave” AHP which followed an ADP. AHP amplitude was in mV and rise times in ms. Firing at rheobase either occurred in the first “Firing Rheo 1st half APs” or the second “Firing Rheo 2nd half APs” half of the 1-s-long current injection, we measured the number of APs in these time windows. “Firing 2X” labeled parameters derived from the two-times rheobasic current injection responses, “Firing MAX” labeled parameters derived from the voltage response with highest firing frequency of a neuron. Interspike intervals (ISIs) were measured as frequencies in Hz at the first or second ISI on Firing 2X and Firing MAX traces. Adaptation ratio was calculated as the ratio of the last 2 interspike interval relative to the first “Adapt 1-last” and second “Adapt 2-last” interspike intervals. Firing frequency (Hz) was determined at double-threshold current injections producing spike trains. “Firing post step” measured the mean of membrane voltage changes in the 150 ms after the 1-s-long current injection step. All parameters were measured by procedures custom-written in Matlab (MathWorks).

### LHA-LHb postsynaptic response

We recorded LHb neurons in voltage clamp mode found within axon terminals from labeled LHA-LHb neurons. The membrane potential was held at -60mV during optogenetic stimulation protocols. Light stimulation was delivered through the microscope lense with blue (473 nm) LED light (SPECTRA X, Lumenchor). Light intensity measured on the slice was 2.5 mW. The illumination lasted for 1 second (5 ms pulse, 20 Hz). The monosynaptic nature of the evoked postsynaptic currents was confirmed with the combined application of TTX (1 µM; Tocris) and 4-AP (100 µM; Sigma-Aldrich) in the aCSF.

### Patch-seq: Cell harvesting for single-cell RNA sequencing

At the end of each patch-clamp routine, we retracted the recording pipette and approached the recorded soma with another pipette (1.8-2.5 MΩ) containing (in mM): 90 KCl, 20 MgCl2. The soma of each recorded neuron was aspirated into the micropipette within a few seconds by applying mild negative pressure (-50 mPa). This procedure allowed us to decrease time of harvesting thus to minimize RNA loss. When we broke contact, the recording pipette was pulled out from the recording chamber and then carefully rotated over an expelling 0.2 ml tube, where its content (∼0.5 µl) was ejected onto a 4 µl drop of lysis buffer consisting of 0.15% Triton X-100, 1 U/µl TaKaRa RNase inhibitor, 2.5 mM dNTP, 17.5 mM DTT and Smart-seq2 oligo-dT primers (5′-AAGCAGTGGTATCAACGCAGAGTACT30VN-3′) (2.5 uM) pre-placed in the end of the 0.2 ml tight-lock tube (TubeOne). The resultant sample (∼4.5 µl) was briefly centrifuged (10 s), placed to dry ice and stored at -80 °C, until subjected to in-tube reverse transcription (RT).

### Patch-seq single-cell RNA-seq library preparation and sequencing

Before reverse transcription, samples were thawed and subjected to 72 °C for 3 min to facilitate complete cell lysis, then cooled to 4 °C. Immediately following the lysis step, 5.5 µl RT mix was added (containing 0.5 µl Superscript II (200 U/µl), 2 µl Superscript II buffer (5X), 0.5 µl DTT (100 mM), 2 µl betaine (5 mM), 0.1 µl MgCl_2_ (100 mM), 0.25 µl TaKaRa RNase inhibitor (40 U/µl), 0.1 Smart-seq2 TSO (5′-AAGCAGTGGTATCAACGCAGAGTACATrGrG+G-3′), and 0.05 µl water, per reaction) resulting in ∼10 µl reaction volume, and RT was performed at 42 °C for 90 min, followed by 10 cycles of 42°C for 2 min and 50°C for 2 min, and finally 72°C for 10 min. Following RT, 15 µl of PCR mix was added (containing 12.5 µl KAPA HiFi HotStart Ready Mix (2x), 0.2 µl ISPCR primers (5′-AAGCAGTGGTATCAACGCAGAGT-3′) (100 mM), and 2.3 µl water, per reaction) and cDNA amplification was performed at 98°C for 3 min, followed by 22 cycles of 98°C for 20 sec, 67°C for 15 s, and 72°C for 6 min, and a final incubation at 72°C for 5 min. Subsequently, cDNA libraries were purified using AMPure-XP beads (0.8:1 ratio; Beckman Coulter) and quantified by Qubit (Life Technologies) and inspected on an Agilent bioanalyzer 2100 (HS dsDNA kit). Tagmentation and subsequent indexing were done using a custom Tn5 as described earlier (*38*). Paired-end (50bp) sequencing was performed on an Illumina Hiseq2500 instrument, obtaining ∼1.5 million reads per cell.

### Extracellular electrophysiological head-fixed recordings

For acute mPFC recordings, two small craniotomies (300-500 µm in diameter) were opened a few hours (> 3 h) before the experiment to access the pre-marked targeted cortical region (+1.95mm AP; 0.3mm and -0.3mm ML). The mice were anesthetized with isoflurane (3 % for induction then 1-2 %). Buprenorphine (0.1 mg/kg s.c.), carprofen (5 mg/Kg s.c.) and lidocaine (4 mg/kg s.c.) was administered. The open craniotomy was covered with Silicone sealant (Kwik-Cast, WPI) and the mouse was returned to its home cage for recovery. Extracellular spikes in the mPFC were recorded using Neuropixels probes (Phase 3B Option 1, IMEC, Leuven, Belgium) with 383 recording sites along a single shank covering 3800 µm in depth. The probe was lowered gradually (speed ∼20 µ.s^-1^) in the left hemisphere with a micromanipulator (uMp-4, Sensapex, Oulu, Finland) until the tip reached a depth of ∼3800-4200 µm under the surface of the pia. The probe was coated with CM-DiI (1,1’-dioctadecyl-3,3,3’3’-tetramethylindocarbocyanine perchlorate, Thermo Fisher, USA) a fixable lipophilic dye for post-hoc recovery of the recording location. The coating was achieved by holding a drop of CM-DiI at the end of a micropipette and repeatedly painting the probe shank with the drop, letting it dry, after which the probe appeared pink. The electrode reference was then connected to a silver wire positioned over the pia in a second craniotomy with a second micromanipulator. The probe was allowed to sit in the brain for 20-30 minutes before the recordings started.

For optotagging experiments, a previously DiD coated (DiD, Vybrant™ DiD Cell-Labeling Solution, Thermo Fisher, USA) optic fiber was gradually inserted at a 25° angle caudally of the Neuropixels probe targeting the LHA, coated with DiO (DiO, Vybrant™ DiO Cell-Labeling Solution, Thermo Fisher, USA). At the end of the recording, a 473 nm DPSS laser (∼ 1 - 2 mW; Cobolt, Solna, Sweden) was used to optogenetically identify LHb-projecting LHA units (> 100 pulses at 1 - 10 ms length were applied for different laser powers).

The signals were filtered between 0.3Hz and 10 kHz and amplified. The data was digitized with a sampling frequency of 30 kHz with gain 500. The digitized signal was transferred to our data acquisition system (a PXIe acquisition module (PXI-Express chassis: PXIe-1071 and MXI-Express interface: PCIe-8381 and PXIe-8381), National Instruments), written to disk using SpikeGLX (Bill Karsh, Janelia) and stored on local server for future analysis.

### 3D whole brain recorded neuron mapping

We defined the D-V and M-L positions as X and Y coordinates respectively by comparing our patch pipette position to the Allen Reference coronal atlas using pictures taken with a 4X objective in DIC optics after the recordings. The precision of the estimations was ∼0.25 mm. A-P position for Z coordinates were estimated using the closest landmarks namely the fornix and the mammillary tract (Extended Data Fig. 2). The XYZ coordinates were then plotted into the Allen reference atlas with a 0.5% jitter to decrease overlap from lateral or dorsal view. The mapping was done by custom-written protocols in Matlab.

### Neuron filling, reconstruction and morphometry analysis

Brain slices containing biocytin-filled neurons were post-fixed in 4% paraformaldehyde in phosphate-buffer (PB, 0.1M, pH 7.8) at 4 °C overnight. Slices were repeatedly washed in PB and cleared using “CUBIC reagent 1” (25 wt% urea, 25 wt% *N,N,N’,N’*-tetrakis(2-hydroxypropyl) ethylenediamine and 15 wt% polyethylene glycol mono-*p*-isooctylphenyl ether/Triton X-100) for 2 days. After repeated washes in PB, biocytin localization was visualized using Alexa Fluor 647-conjugated streptavidin. Slices were then re-washed in PB and submerged in “CUBIC reagent 2” (50 wt% sucrose, 25 wt% urea, 10 wt% 2,20,20’-nitrilotriethanol and 0.1% v/v% Triton X-100) for further clearing. Slices were mounted on Superfrost glass (Thermo Scientific) using CUBIC2 solution and covered with 1.5 mm cover glasses. *Post-hoc* dendritic morphology reconstruction and measurements (simple neurite tracing plugin, Fiji), sholl analysis (sholl analysis plugin, Fiji) and somatic measurements were performed in Fiji after z-stacks images acquired by a confocal microscope (Zeiss 880). Somatic morphometry was performed on orthogonal projection of z-stack images. Two major perpendicular axes of the cells (X, Y, 90 degree angle) were used to generate the elliptical (E) score (X/Y, if <1.8 round or triangular soma shape, if >1.8 triangular or elongated soma shape). 25% and 75% perpendicular axes to X (Y1= major and Y2 = minor) were used to generate the triangular (T) score (Ymajor/Yminor, if < 1.2 round or elongated soma shape, if > 1.2 triangular soma shape). t-SNE distribution of the LHA-LHb neuronal types based on somatic morphometry was performed using 7 somatic parameters (area, X,Y, Ymajor, Yminor, E score, T score; as in Extended data 2d).

### Immunohistochemistry

Brain sections were cut on a vibratome at 50 µm thickness (Leica VT1000, Leica Microsystems GmbH). Sections were rinsed in phosphate buffered saline (PB), blocked for 2 hours in 10% normal donkey serum with 0.5% Triton X-100, and then incubated in primary antibody overnight at 4°C. Primary antibodies were as follows: rabbit anti-Esr1a (1:1000; Santa cruz biotechnology, Cat#sc-542), rabbit anti-Galanin (1:1000; Gift from E. Theodorsson, AB_2314521); rabbit anti-Npy (1:1000; Peninsula Laboratories International, Cat#T-4070); guinea pig anti-Orexin A (1:1000 Synaptic system, Cat#389 004); chicken anti-V5 (V5; 1:500 dilution; Abcam; ab9113); rabbit anti-cFos (cFos; 1:500 dilution, Santa Cruz Biotechnology; sc-52); Cy5 anti-Streptavidin (1:1000; Cat#016-170-084). After extensive washing with PB, immunoreactivities were revealed using Cy5-conjugated secondary antibodies (1:500, Jackson ImmunoResearch, Cat#711-175-152 or Cat# 706-175-148) and DAPI (1:50000) for 2 hours at room temperature. All antibodies were diluted in carrier solution consisting of PB with 1% BSA, 1% normal goat serum, and 0.5% Triton X-100. Sections were then extensively rinsed in PB, mounted on slides, and coverslipped.

### Neuropixels probe track reconstruction

Brain sections were cut on a vibratome at 400 µm thickness (Leica VT1000, Leica Microsystems GmbH). Slices were repeatedly washed in PB and cleared using “CUBIC reagent 1” (25 wt% urea, 25 wt% *N,N,N’,N’*-tetrakis(2-hydroxypropyl) ethylenediamine and 15 wt% polyethylene glycol mono-*p*-isooctylphenyl ether/Triton X-100) for two days. After repeated washes in PB, slices were incubated with DAPI (1:50000) for one day at room temperature. Slices were then re-washed in PB and submerged in “CUBIC reagent 2” (50 wt% sucrose, 25 wt% urea, 10 wt% 2,20,20’-nitrilotriethanol and 0.1% v/v% Triton X-100) for further clearing. Slices were mounted on customized 400 µm thick slides using CUBIC2 solution and covered with 1.5 mm cover glasses. The blue and red channels were imaged at 4x using a Zeiss 880 confocal microscope. For each brain section 6 to 7 z-stacks spaced by 50µm were obtained and down sampled to a 10 µm resolution. The z-stacks containing the probe red fluorescent signal were further registered in the Allen CCFv3 and the probe position was estimated using the available ‘SHARP-Track’ pipeline (https://github.com/cortex-lab/allenCCF). As the electrodes are spaced 20 µm apart (center to center) on the shank of the probe, the location along the electrode was transformed into the CCFv3 space based on the orientation and position of the probe track. Unit locations were assigned based on the location of the electrode where that unit had the highest waveform amplitude.

### Image acquisition and analysis

All confocal images were taken using a Zeiss 880 confocal microscope. CUBIC cleared sections after slice electrophysiology and biocytin staining were acquired as z-stacks using a Plan-Apochromat 20x/0.8 M27 objective (imaging settings: frame size 1024x1024, pinhole 1AU, Bit depth 16-bit, speed 6, averaging 2). For viral expression overview Plan-Apochromat 10x/0.45 M27 objective was used (imaging settings: frame size 1024x1024, pinhole 1AU, bit depth 8-bit, speed 7, averaging 2). For viral expression and immunohistochemistry close up Plan-Apochromat 20x/0.8 M27 objective was used (imaging settings: frame size 1024x1024, pinhole 1AU, Bit depth 16-bit, speed 6, averaging 2). Projection of axon terminals in LHb were imaged using a Plan-Apochromat 63x/1.40 Oil DIC M27 objective was used (imaging settings: frame size 1024x1024, pinhole 1AU, Bit depth 16-bit, speed 6, averaging 4).

### Input specific 3D habenula map

Acquisition settings of confocal microscope images were kept identical for all sections within the experiment. The LHb was first delineated by manual drawing of contours on each coronal section. Sections from the four different genotypes were aligned at each unique coordinate along the anteroposterior axis by translation and rescaling before defining a common outline. For each section, pixels within the LHb were labelled in a binary manner as either “terminal” or “not terminal” by thresholding fluorescence level. The LHb was then binned into 4.1 µm² squares and the terminal density of these spatial bins was computed using a two-dimensional gaussian kernel, resulting in a heatmap of terminal density across spatial bins for each section and each genotype. To account for differences in baseline fluorescence between genotypes, the density was normalized. For visualization purposes, the expression was linearly rescaled between 0 and the maximum density across all sections of a genotype. For further analysis, we used another normalization to better spread the values within the entire dynamic range. Spatial bins were sorted by increasing terminal density across all sections of a genotype and partitioned into 100 sets of equal cardinalities - each set corresponding to one percentile of the data. Then, the LHb was subdivided by assigning each spatial bin to the genotype with the highest percentile density when this value was above the 50th percentile; otherwise, the region was left unassigned. For three-dimensional (3D) visualization, we used the 3D meshes from the Common Coordinate Framework v3 (CCFv3) 2017 of the Allen Brain Atlas to define the LHb delimitations. We performed the above-mentioned analysis to assign each voxel from a 3D grid to one of the four genotypes, without the 50th percentile threshold for smoother visualization. We then extracted a 3D volume definition for each genotype using the marching cubes algorithm implementation from the *Rvcg* R package. Visualizations were rendered using the OpenGL implementation from the *rgl* R package.

### *In vivo* optogenetics

In optogenetic experiments, mice were bilaterally implanted with optical fibers aimed above the LHb terminals (+1.55 mm AP; 0.95 mm ML from bregma and 2.2 mm depth with 10° angle from the dura) or the cell bodies in the LHA (+1.1 mm ML; 4.25 mm from the dura; the AP coordinate was adjusted according to peak coordinates in each genotype: Pv-cre: 0.9 mm, Esr1-cre: 1.1 mm, Npy-cre: 1.2 mm, Vglut2-cre and C57BL/6J: 1.1 mm). Optical fibers were home made using an optic fiber of 200 µm diameter (FG200UEA, Thorlabs) and matching ferrules (CFLC230; Thorlabs). Mice were coupled via a ferrule patch cord (D204-2128, Doric Lenses) to a ferrule on the head of the mouse using a split sleeve (ADAL1-5, Thorlabs). The optical fiber was connected to a laser (MLL-III-447-200mW laser) via a fiber-optic rotary joint (FRJ_1x1_FC-FC, Doric Lenses) to avoid twisting of the cable caused by the animal’s movement. After a testing session, animals were uncoupled from the fiber-optic cable and returned to their home cage. The frequency and duration of photostimulation were controlled using custom-written Arduino script. Laser power was controlled by dialing an analog knob on the power supply of the laser sources. Light power was set to 8 mW measured at the tip of the ferrule in the patch cord before each experiment, using an optical power and energy meter and a photodiode power sensor (Thorlabs). Animals showing no detectable viral expression in the target region or ectopic fiber placement were excluded from the analysis. Fiber placement of each mouse included in the analysis is reported in Extended Data Fig. 4.

### Behavioral tests

Mice were handled for three consecutive days prior behavioral experiments. Mice were acclimated to the behavioral room for 1 hour before testing. Animal behavior was recorded with a CCD camera with infrared filter interfaced with a Biobserve software (Biobserve GmbH). Red and infrared light were the only light sources present during all behavioral experiments. When multiple behavioral tests were performed within the same subject, one week was left between two experiments. For behavioral phenotyping after shock exposure, mice underwent three consecutive tests within the same day, then combined in the stress index.

### Real time place aversion test

Mice were placed in a custom-made two compartment behavioral arena separated by a wall with an opening in the middle (50 × 25 × 25 cm black plexiglass) for 10 minutes. The behavioral arena was placed on a transparent plexiglass. The animal behavior was recorded with a camera placed below the arena. The mouse performance was evaluated under three different conditions during two consecutive days. The first day optical fibers were connected to the animal but there was no light stimulation, the second day one of the compartments was paired with light stimulation (40 Hz, 5 ms pulse, 473 nm laser, see Video 1). Right after the test, the stimulation paired side was switched. Mice were tracked using DeepLabCut (*39*). We manually labeled the base of the tail base, left and right hindlimbs and forelimbs in 1000 frames, sampled from all sessions. After running DLC on every video, we transformed the video coordinates (in pixels) to world coordinates (in cm) using a perspective transform matching the four corners of the box. We based our movement analysis on the tail base point tracked by DLC.

### Conditioned place aversion test

Mice were placed in a custom-made two compartment behavioral arena (50 × 25 × 25 cm black plexiglass, paired compartment walls had white on black stripes) and the behavior was recorded as in the real time place aversion. The first day optical fibers were connected to the animal but there was no light stimulation, the second day the animal was forced to spend time in the compartments paired with light stimulation (40 Hz, 5 ms pulse, 1 second on, one second off, 473 nm laser). The third day the optical fibers were connected to the animal but there was no light stimulation, and the animal was free to explore both compartments for 10 minutes.

### State induction test by optogenetic stimulation

We adapted a conditioned place aversion (CPA) assay to test the presence and persistence of a behavioral state induced by LHA-LHb pathway stimulation. Mice were placed in a custom-made two compartment behavioral arena (50 × 25 × 25 cm black plexiglass, paired compartment walls had white on black stripes) and the behavior was recorded as in the real time place aversion. The first day optical fibers were connected to the animal but there was no light stimulation. Immediately after the habituation session, the animal was forced to spend time in the compartments paired with light stimulation (40 Hz, 5 ms pulse, 1 second on, one second off, 473 nm laser) for 10 minutes (state induction, Figure 3). Immediately after the state induction session, the animal was free to explore both compartments for 10 minutes (immediate state). The second day (24 h post induction) the animal was placed back to the same arena free to explore both compartments for 10 minutes (sustained state). For the habituation session, the immediate state test and sustained state test the optical fibers were connected to the animal but there was no light stimulation. We manually labeled the base of the tail, left and right hindlimbs and forelimbs in 1000 frames, sampled from all sessions. We based our movement analysis on the tail base point tracked by DLC. After running DLC on every video, we transformed the video coordinates (in pixels) to world coordinates (in cm) using a perspective transform matching the four corners of the box. During the 10 minutes state induction, discrete events of stop-backwards, sharp turns, digging and free rearing (number of events of standing on the hind limbs, far from the arena wall) where manually scored for each second during the stimulated and unstimulated epochs.

### Open field test

Mice were placed in a custom-made open field (49×49 cm black plexiglass) for 10 minutes (TeLC silencing experiment, Extended Data Fig. 6ab) or 20 minutes (optogenetic stimulation experiment, Extended Data Fig. 6r). The behavioral arena was placed on a transparent plexiglass. The animal behavior was recorded with a camera placed below the arena. For the optogenetic experiment, the mouse performance was evaluated under alternating laser off/on epochs, starting with 5 minutes off (stimulated epoch: 40 Hz, 5 ms pulse, 1 second on 500 ms off for 5 minutes, 473 nm laser). We manually labeled the base of the tail, left and right hindlimbs and forelimbs in 1000 frames, sampled from all sessions. After running DLC on every video, we transformed the video coordinates (in pixels) to world coordinates (in cm) using a perspective transform matching the four corners of the box. The speed (cm/s) and stationary time were analyzed on the base of the tail tracked by DLC. For the optogenetic experiment, discrete events of wall rearing (number of events of standing on the hind limbs, with forepaws on the arena wall), free rearing (number of events of standing on the hind limbs, far from the arena wall) and discrete grooming events (number of events of mice in sitting position with licking of the fur, grooming with the forepaws, or scratching with any limb) where manually scored for each second during the stimulated and unstimulated epochs (See Extended Data Video 2).

### Free-Access Caloric Consumption Assay

Mice were food-restricted to 85 to 90% of their initial body weight by administering one daily feeding of ∼2.5 to 3.0 g of standard grain-based chow (immediately following behavioral experiment, if performed). Water was provided ad libitum. All feeding-related behavioral experiments were conducted at the same time in the middle of the animals’ dark cycle (at approximately 14:00). Food restricted mice were placed in a custom made 15 × 15 × 20 cm operant chamber with free access to a bottle containing a 15% sucrose reward for 40 min. Each lick (lick response) was detected, and reward was delivered (rewarded lick, 3 µL reward) with a 1 second timeout. Mice were habituated to the operant chamber and connected to the fibers with no light stimulation. Once stable licking was achieved with light-off sessions (three consecutive days of daily average lick responses within ± 10%), sucrose consumption was monitored for two consecutive days of light-off and two consecutive days of light-on stimulation (40 Hz, 5 ms pulse, 1 second on, one second off, 473 nm laser).

### Quantification of cFOS in LHb

In order to evaluate the recruitment of LHb neurons we performed a cFOS IHC and quantified the cFOS+ neurons with the lateral habenula area, after confocal imaging of the LHb throughout the anterior-posterior axes. For optogenetic experiments (Extended Data Fig. 6o-p), mice were placed in a custom-made open field (49×49 cm black plexiglass) where they received 10 minutes of simulation protocol as in state induction test (40 Hz, 5 ms pulse, 1 second on, one second off, 473 nm laser). Mice were sacrificed 30 minutes after the start of the stimulation. For optogenetic experiments, the control group was implanted with optic fibers, placed in the same open field for 10 minutes and optical fibers were connected to the animal but there was no light stimulation. For TeLC experiments, mice were placed in a sound isolated fear-conditioning chamber where they received 5 inescapable, uncontrollable electric foot shocks, at 0.3 mA over 10 minutes with random shock duration ranging from 1 to 3 seconds and unpredictable inter-shock intervals (ITIs), from 1 to 15 seconds. Mice were sacrificed 30 minutes after the start of the foot shock protocol.

### Stress induction protocol

In order to induce stress-like state in mice for electrophysiological recordings and behavioral tests, mice were exposed to a stress induction ‘training protocol’ for 3 days. Stress induction protocol was modified to ‘mild foot shock’ from what previously described in order to maximize the chance of detecting sexually dimorphic effect and avoiding ceiling effect. Mice were placed in a sound isolated fear-conditioning chamber where they received 360 inescapable, uncontrollable electric foot shocks at 0.3 mA over 1hour with random shock duration ranging from 1 to 3 seconds and unpredictable inter-shock intervals (ITIs), from 1 to 15 seconds. When possible, experiments were performed on pairs of littermates previously housed in the same cage. Control animals were placed in the shocking chamber for 1 h, without being shocked. 24 h after the last shocking protocol mice were assigned to slice electrophysiology or behavioral phenotyping. Animal identity was blinded to the researcher who performed the electrophysiological recordings and the scoring of the behavioral tests.

### Stress Index

All behavioral experiments were consistently performed between 9 and 12 AM. The researcher who performed the test was blinded to the mice cohort. In order to build a reproducible index of stress level in mice we build a stress index, combining three behavioral tests. Tests were performed with an interval of 30 minutes, in an increasing level of aversiveness: all animals were first tested in the marble burying test (MBT), then in the looming test (LST) and finally in the forced swim test (FST). The stress index was built combining one parameter for each test using the Euclidean distance of z-scored normalized values. Parameters used: buried marbles (n, in the MBT), time spent in behaviors classified as aversive (s, in the LST), time spent immobile (s, in the FST).

### Marble burying test (MBT)

The marble burying test was performed as previously described (*40*). Briefly, 20 clean glass marbles of diameter 1.5 cm and homogeneous color were disposed of in a 5x4 matrix on a 5 cm deep sawdust without food and water. One by one, animals were placed in the cage for 30 minutes. At the end of the due time, mice were returned to their cage, and the number of buried marbles was scored by two blinded experimenters. Marbles were considered buried if covered by sawdust by at least two-third. Before starting the testing of a new mouse, marbles were cleaned with 70% ethanol and placed in a newly prepared cage.

### Looming stimulus test (LST)

30 minutes after the MBT, stressed and control mice performed a modified version of the looming test (*41*) in a custom designed 8-shaped arena. The looming arena, classically composed of one only squared arena, was here modified to an eight-shaped field, obtained by merging two round arenas, 30 cm diameter; 23.5 height, with black matt walls to prevent reflection of the stimulus. An opening between the two circular chambers allowed free exploration. No shelter was provided, as this arena design allowed for escape to the opposite compartment as a defensive strategy, where the animal behavior was monitored. A monitor was placed on the ceiling of the arena, providing dim lighting from the gray screen of the monitor. As for the rest of the behavioral test, infrared illumination, invisible to the mouse, was provided for video recording. The arena was placed on a transparent plexiglass. The animal behavior was recorded with a camera placed below the arena. Thanks to these modifications, the looming stimulus was reliably repeated three times. The looming stimuli was triggered by the experimenter once the animal was in the center of one of the arenas, as previously described (*41*). In brief, the stimulus was repeated 15 times with increasing diameter (from 2 to 20 degrees of visual angle) for the first 250 seconds, and then stable at 20 degrees for the remaining 250 ms. The next stimulus was presented when the animal was in the center of any of the two arenas, with a minimum of one-minute interval. Behaviors were scored from video recordings by two blind experimenters. A post-stimuli epoch of 60 seconds after each stimulus was analyzed. Each second was assigned to aversive (escape to opposite compartment, freezing, tail rattling, immobility, periphery) and non-aversive (normal walking, grooming, sniffing) behavior and reported as cumulative time spent in aversive behavior over the three trials.

### Forced swim test (FST)

30 minutes after performing the looming test, stressed and control mice were individually placed in a transparent acrylic cylinder (height: 60 cm, diameter: 14 cm) containing 2 L of clear water at 25 ± 1° C for 6 min. The cylinder was placed on a transparent plexiglass. The animal behavior was recorded with a camera placed below the arena. Water was changed between subjects. Delay to immobility (delay in seconds to the first immobility) and time spent immobile (seconds spent floating passively in the water) were manually scored by a researcher blind to the animals’ cohort.

### Fear conditioning

Mice were placed into a sound isolated TSE Multi Conditioning System where they received five tones followed by mild foot shocks (5 second tone, 0.3 mA foot shock during the last 2 s of the tone) with randomized inter-shock intervals. The following day, conditioning to the tone was tested in the same TSE Multi Conditioning System, placing the mouse in a new context (arena shape and smell) where they received 10 tones (5 seconds tone) with randomized inter-tone intervals.

### Acoustic startle test

Mice were placed into a sound isolated TSE Multi Conditioning System to habituate for 5 minutes. Afterward the habituation period they were exposed to 10 tones (5 seconds tone, 70 dB, 10 kHz) with randomized inter-tone intervals.

### Head-fixed aversive conditioning protocol

A total of 25 male mice (10 Esr1-cre, 5 Npy-cre, 5 Vglut2-cre and 5 C57BL/6J) were exposed to the head-fixed aversive protocol. The researcher who performed the recordings was blinded to the mice cohort. Mice were first habituated to be head-restrained over a period of three to five days to reduce stress levels. An ambient white noise (70 dB) was played continuously to provide a homogeneous auditory background. Behavioral control and behavioral data collection were carried out with custom-written computer routines using a National Instruments board interfaced through LabView or Matlab (Mathworks). The aversive conditioning protocol was divided into eight blocks. Each block being separated by at least one minute from the others. In the first block, a 10kHz pure tone (sound pressure: 80 dB, duration 200 ms), was presented 50 times with a random 3 s to 9 s inter-trial interval (ITI) coming from a uniform distribution. For the second block, the pure tone was followed by a 500 ms pulse light train with a frequency of 40 Hz. Each light pulse has a duration of 5ms with 1ms sinusoidal ramp in and 1ms sinusoidal ramp out. The light power at the tip of the fibers was 6 mW and was systematically measured and adjusted before every experiment. The delay between the onset of the pure tone and the optogenetic stimulation was 500 ms and 100 trials were presented to the mice with a 6 s to 12 s random ITI. The third block was identical to the first block. For the fourth block, a new sound (blue noise, sound pressure: 80 dB, duration 200 ms) was followed by a 50 ms mild air puff delivered into the face of the mice. The delay between the onset of the blue noise and the air puff was also 500 ms. Block five to eight consisted of 50 trials each of the optogenetic stimulus alone (same as in block 2) with four ranging increasing light powers (0.3 mW, 2 mW, 6 mW and 10 mW). The behavior of the mice was monitored with a Blackfly camera (Teledyne FLIR, USA) during each block with a sampling frequency of 50 Hz. A patch cable connected to a laser (Cobolt MLD 473 nm) controlled by custom behavioral procedures (Matlab) and interfaced through a PCI-6221 card (NI) was used for light delivery.

## QUANTIFICATION AND STATISTICAL ANALYSIS

### Clustering of cell types by intrinsic electrophysiology

For unbiased clustering of neurons based on their electrophysiological properties, we applied successive transformations to normalize the dataset. First, we shifted each parameter positive and starting at zero through a translation by the minimum feature value. Then values for each parameter were divided by the sum of the corresponding parameter across the entire dataset and scaled by a factor 10000. We then applied the log(1+x) transformation and scaled the values for each feature by applying z-score. We used PCA for dimensionality reduction and selected the first 4 principal components. Two clustering algorithms were then applied to unbiasedly identify six clusters, i.e. the same number of cell types as defined by the expert classification. First, we applied hierarchical clustering with Ward’s method. Then we performed graph-based clustering using the *Seurat* package implementation (*k* parameter was set to 20 and resolution to 1.5). As each clustering method generates a slightly different partition of the dataset, we combined them by a consensus clustering: only cells assigned to the same cluster by both algorithms were kept, others were discarded (n = 19). We rendered two-dimensional visualizations by applying the t-SNE algorithm to the 4 principal components with a perplexity of 30. The accuracy of clustering methods was defined as the percentage of cells assigned to the same cluster with the unbiased clustering and the expert-based annotation. The accuracy which can be expected by chance was estimated by randomly shuffling the unbiased cluster assignment 10000 times and averaging the corresponding accuracy values.

### Patch-seq analysis

Raw data was adapter- and quality trimmed with fastp v0.20.0, using default settings, then pseudo-aligned to GRCm38.p6 protein-coding transcripts from GENCODE vM22 using Salmon v0.14.1 (-- libType IU --validateMappings). Shell commands were run in parallel using GNU parallel. Transcript-level estimates were collapsed to gene-level using tximport v1.12.3 then converted to a SingleCellExperiment object using scater v1.12. Outliers were detected batch-wise based on 3 median absolute deviations (MADs) from median based on (log10) total read counts, (log10) genes detected and percentage mitochondrial counts (lower tail, lower tail and upper tail, respectively) using scater. Next, gene-specific variances were decomposed into biological and technical components as implemented in scran v1.12.1 using experimental batches as a blocking factor. We defined highly variable genes (HVGs) as the top 500 variable genes, ordered by biological variance component and FDR (FDR < 0.05). Dimensionality reduction using UMAP or t-SNE was performed on the first 50 principal components for HVGs as implemented in scater. Graph-based Louvain clustering was performed on the first 50 principal components for HVGs based on Spearman’s rank correlation cell-cell distances as implemented in scran. For discrete groups, we extracted molecular markers of the electrophysiological cell types in a pairwise combination comparison of the 6 distinct neuronal types by calculating a log2-fold gene expression difference as the ratio of the TPMs/gene across the 6 types. For visualization of the markers, we filtered the gene expression to exclude single cells belonging to a cluster where a the marker expression was below 20% of the cells. To determine whether the gene expression across two cell types were statistically significant we modeled TPMs according to a negative binomial distribution and specified the type of linkage between the variance and the mean as a locally regressed non-parametric smooth function of the mean. This comparison delivered the P-values which had been adjusted to account for the multiple testing problem using the Benjamini-Hochberg adjustment with a threshold of 0.05 for the adjusted P-values (Q-values), equivalent to consider a 5% false positive as acceptable, and identify the genes that are significantly expressed by considering all the genes with adjusted P-values below this threshold.

### Statistics on sex-difference of intrinsic electrophysiology in stress

Same intrinsic parameters were extracted for the analysis as for the classification and clustering of LHA-LHb neuronal types. Parameters were compared across cell types and sexes using Data Analysis with Bootstrap-coupled ESTimation (DABEST), a permutation based graphical estimation which steps away from dichotomic null-hypothesis significance testing (*42*). In short, the analysis bootstraps the pairwise comparison of means of randomly shuffled groups, producing a Gardner-Altman plot, which visualizes effect size as the distribution of mean differences which also defines a 95% confidence interval (CI). P-value reflects the proportion of group comparisons which resulted in a mean difference outside the CI.

### *In vivo* optotagged LHA-LHb units

To identify optotagged LHA-LHb units (light responses in > 10 % of trials in 10 ms bin after laser pulse), the stimulus-associated spike latency test (SALT; α < 1 %) was used. To compare light-evoked with spontaneous spike-waveforms, Pearson’s correlation coefficient (r > 0.9) was used.

### Neuroanatomical analysis

For ChR2-mCherry neuron quantification, following brain sectioning, endogenous signal of mCherry expressing neurons was used by semi-automatic counting using Imaris software. For axon terminals quantification in the input specific 3D habenula map, confocal images were acquired maintaining consistent settings within the genotypes. Orthogonal projections were then processed in ImageJ (Fiji) software, where a consistent threshold was applied to transform the fluorescence signal into binary data, therefore rescaled, registered and mapped into a common reference atlas, as detailed. Data are represented as mean ± SEM or as box plots. Single data point where overlaid to the box plots, to identify individual neurons. For dendritic and somatic morphometry, we to test pairwise differences between groups we used one-way ANOVA and reported Tukey’s multiple comparisons p-values in Supplementary Table 3.

### Optogenetic and behavioral experiments

In all experiments, significance levels of the data were determined using custom written Matlab script. Two-tailed unpaired or paired Student’s t test when comparing two independent or dependent groups, respectively. For the comparison between more than two groups, we used one-way ANOVA. To further test pairwise differences between groups we used Tukey’s Multiple comparisons test. Data are represented as mean ± standard deviation or SEM as reported in the figures, or as boxplots. Single data point where overlaid to the box plots, to identify individual animals. In the optogenetic stimulation experiments, when both groups were performed, open circle single data points represent somatic stimulation of retrogradely labelled neurons in LHA, while solid gray circles represent terminal stimulation in LHb of anterogradely labelled LHA neurons.

### Pupillometry

We tracked the pupil of the mice using DeepLabCut (DLC; (*39*)). We manually labeled the pupil (4 points) and the eye (4 points) in 2000 frames, uniformly sampled from all mice and blocks. After running DLC on every video, we transformed the video coordinates (in pixels) to a pupil and an eye area (ellipse fitting, pixels.^2). Each pupil area value was z-scored to block 1 per mouse. For the heatmap visualization, normalized pupil area values were averaged per genotype. For the bar plots and the scatter plots, normalized pupil area values were averaged per genotype and per trial. Significant change of pupil size between blocks per genotypes was tested on the trial grand average values as observations using a Mann-Whitney U test with Bonferroni correction for multiple comparisons.

### Spike sorting

The high-pass filtered action potential (AP) extracellular potential data were preprocessed using common-average referencing: the channel’s median was subtracted to remove baseline offset fluctuations, then the median across channels was also subtracted from each channel to remove artifacts. The data was then automatically spike sorted with Kilosort 2.0 (https://github.com/MouseLand/Kilosort/releases/tag/v2.0) and then manually curated using the phy2 GUI (https://github.com/cortex-lab/phy). During manual curation, clusters of waveforms showing near-zero amplitudes or non-physiological waveforms were classified as ‘noise’. Clusters with inconsistent waveform shapes and/or refractory period violations were classified as ‘multi-unit’. The remaining units were classified as ‘good’. These potential good units were finally investigated against spatially neighboring clusters. Units showing similar waveforms, clear common refractory periods and putative drift patterns were subjected to a merge attempt. If the resulting cluster was still showing consistent waveforms and a clear refractory period in their auto correlogram, the merge was kept. Splits of the clusters were also performed on a few occasions where the principal features of the waveforms showed distinct clusters and two or more groups of waveforms could be identified. Only the ‘good’ units were kept for the following analysis.

### *In vivo* electrophysiological data analysis

The spike times were corrected for the temporal drifting along the recording (∼10 ms/h) relative to a clock signal registered independently by the PXIe acquisition module and the PCI-6221 card login the behavioral signals. First the temporal drift between the two devices was measured for each recording. Second, a linear regression was applied to correct the timestamps. Then, only units from four PFC subregions (ACAd, PL, ILA, ORBm) with firing rate > 0.1 Hz and ISI violation of less than 1% were selected. Spikes falling into LFP saturation periods were removed with a margin of 1 s before and after saturation. Spike trains were aligned to the trial start and binned at 10 ms resolution for analysis. For the heatmap visualization, the firing rates were averaged per unit across trials and per block, smoothed with a 50 ms gaussian kernel and z-scored to the baseline per block. To consistently average the same number of trials per block (50 trials), one trial over two was removed from the block 2 trial pool. The units were then sorted either by PFC region (ACAd, PL, ILA and ORBm), by the genotype of the line they originated, by their peak response time in the 500 ms window following the 10 kHz auditory pure tone stimulus in block 1 or by a combination of the preceding options. The same sorting order was kept across all blocks.

### Generalized linear model (GLM)

We used a generalized linear model to describe how each single unit firing activity was related to our four task events (sound 1, optogenetic stimulation, sound 2 and air puff). Each single unit rate was binned at 100 ms and modeled as a linear combination of the four regressors also binned at 100 ms (sound 1 and 2; 200ms on after sound onset, optogenetic stimulation; 500ms on after stimulation onset, air puff; 500ms after air puff onset). We used the Matlab function glmfit.m with a Poisson link function. Four beta coefficients, for each regressor, were obtained per fitted unit.

### Tuning scores

For each recording session we created surrogate data by circular shifting the firing rates at a random time point of the recording session. This procedure decreases the risk of false positive errors as only the connection between the regressors and the neural activity is broken. To calculate the tuning score, we calculated the beta coefficients per unit with the GLM fit previously described and compared the obtained values to the distribution of beta coefficients obtained from 1000 iterations of the same GLM fit on circular-shifted surrogate data. The tuning score of one unit per regressor was calculated as the difference of the true beta coefficient form the mean of null distribution (1000 beta coefficients obtained by circular shift) divided by the standard deviation of the null distribution. We obtained four tuning scores per units referred to as its *tuning profile*. Positive and negative tuning scores can be obtained. We refer to the absolute maximum of these four tuning scores as the primary tuning of the unit.

Additionally, we classified units as *positively significantly tuned* to a regressor if the beta coefficient of this unit is larger than the 99.9 percentile of the distribution of the beta coefficients of circular shifted shuffled iterations. Reversely, *negatively significantly tuned* units are obtained when the beta coefficient is less than the 0.01 percentile of the shuffled distribution.

### Principal component analysis (PCA)

Spikes of every unit were binned with a 10 ms non-overlapping window in the interval of [-0.1 s, 3 s] relative to the trial onset. Spiking activity was averaged across trials within each block (50 trials in block 1, three, four; 100 trials in block 2). Spike counts were z-scored and convolved with a Gaussian kernel (sigma = 2). Units were pooled together across mice within each genotype (n = 10 mice for Esr1-Cre, n = 5 mice for other genotypes). To balance the number of units, a random subset of 550 units was drawn for each genotype. We performed the dimensionality reduction using Principal Component Analysis (PCA) in order to plot the trajectories of neural activity within blocks for each genotype. For a single block we built a **N x T** matrix of neuronal activity, where **N** is the number of units pooled together from four different genotypes (4 x 550), **T** is the number of time bins during the trial. We used PCA (python, *scikit-learn 0.23.2*) to project the activity of all genotypes to the common activity space with three principal components, and then projected the activity of each genotype on its corresponding principal component (PC) loadings in order to visualize them independently. The procedure was repeated for each of four blocks. Visual inspection confirmed that randomly selected subsets of 550 units resulted in stable trajectories, therefore Figure S9a shows representative plots.

### Hierarchical clustering

Trial averaged firing rates (10 ms bin) of the GLM fitted units (n = 1945) were smoothed with a gaussian kernel (sigma 0.8) and z-scored across four blocks. A quality control step was performed: units with less than 200 active (firing rate < 0.5 Hz) bins across all blocks (1600 bins in total) were excluded from the analysis (n = 1788 units remaining). PCA was then applied to the activity matrix and the top 20 PC loadings were extracted for each unit. Hierarchical clustering using the PC loadings as features was then applied (Ward’s method). The obtained dendrogram was cut at the depth of 10 children. The resulting 15 activity clusters were given subjective names reflecting their principal primary tuning (‘opto’ for primary tuning to the optogenetic stimulation, and similar for ‘air puff’ and ‘sound’). When a significant modulation (Wilcoxon signed rank test with Bonferroni correction) of the mean 1s baseline activity preceding the auditory stimulus onset was observed across some blocks, the name ‘state’ was used. ‘opto’ and ‘air puff’ clusters are also referred to as ‘aversive signal’ clusters.

### Conditioned units

To determine if a unit was conditioned by the aversive stimuli, a set of four Wilcoxon signed-rank tests was run to detect changes in activity upon the 10 kHz pure tone in block 2, the blue noise in block 4, the optogenetic stimulation and the air puff: Each statistical test was performed by comparing the mean firing rate defined as baseline (500 ms before the auditory stimulus (pure tone or blue noise presentation) and the mean firing rate following the selected event (500 ms time window) for each unit across equivalent number of trials (50). Conditioned units displayed significantly (Wilcoxon signed rank-test, Bonferroni corrected alpha of 0.001/4) increased or decreased response to the sound and optogenetic manipulation in block 2 (top) or to the sound and air puff in block 4 (bottom) over the trials within the respective block.

### Activity modes

From all the task-modulated units, we made an activity matrix where each row is the trial averaged firing rate of each unit (50 ms bin) across four blocks. We next subtracted the per-unit firing rate across all blocks and smoothed each trial averaged firing rate with a gaussian window (sigma 0.8). Then, we computed four activity modes (directions in the activity space that optimally separate different task events and epochs): the ‘sound’ mode, the ‘opto OR air puff’ mode, the ‘opto AND air puff’ mode, and the ‘state’ mode. We used the data from all genotypes to compute the modes and visualize the projections onto them. The ‘sound’ mode was computed as the weights given by a support-vector machine (SVM) network trained to discriminate if a sound occurred or not (in a 200 ms time window including some baseline preceding the sound onset or the activity following the sound onset). This ‘sound’ mode results in a n x 1 vector containing the SVM weights for each unit where n is the number of task-modulated units. The ‘opto OR air puff’ mode, the ‘opto AND air puff’ mode and the ‘state’ mode were constructed similarly. For the ‘opto OR air puff’ and the ‘opto AND air puff’ modes we used the 500 ms time window following either the optogenetic stimulation onset or the air puff onset to train the SVM to discriminate between the two types of aversive stimuli. For the ‘State’ mode we trained the SVM on the 1s baseline activity preceding the sound onset in block 1 vs block 4. The analysis yielded three n x 1 vectors of SVM coefficients (n is the number of units). We applied QR decomposition to these vectors to form **W**, the n x 3 orthogonal matrix (using either the ‘OR’ mode or the ‘AND’ mode).

The trial averaged data per block (**x** = n x 4 x t matrix where t are the time bins), was then projected onto these axes as the dot product **W^T^x**. For the data per genotype **W** and **x** were restricted only to the units belonging to one genotype.

### Decoding (logistic regression)

For decoding, spikes during baseline period of [-2.1 s, -0.1 s] before the trial onset were binned with a non-overlapping window of 100 ms. The activity was concatenated across blocks (N x 4T, where N is the number of units in one genotype, T is the number of time points in one block of trials) and z-scored within each unit. Units with no spikes during baseline were removed. We decoded the binary block identity in block 1 versus block 2, or in block 1 versus block 4, independently within each genotype and mPFC subregion to find out how accurately a neuronal population could decode aversive versus non-aversive state. We randomly selected a different number of units (from 5 to 150) to train the logistic regression (python, scikit-learn 0.23.2) on 50% of time bins and to test it on the remaining 50% and for each subset repeated this procedure 50 times (repeated 2-fold cross-validation). Therefore, each data point depicts mean and standard deviation of 50 repeats. Dashed line shows chance performance (50%). In some areas less than 150 neurons were recorded and in such cases accuracy trace ends at the number of recorded units.

## Acknowledgments

We thank Joanne Bakker and Tomas Hökfelt for sharing Gal-cre mice, and Elvar Theodorsson for sharing the anti-galanin antibody. We thank Ram Yahya and Fredrik Wernstal for help with spike sorting. Funding for the study was provided to K.M. by the Swedish Research Council, StratNeuro, Hjärnfonden; to M.C. by the Wallenberg Scholar programme (Knut and Alice Wallenberg Foundation); to M.C. and K.M by a KAW project grant (Knut and Alice Wallenberg Foundation); postdoctoral funding to D.C. from the Hagelén Foundation and Hjärnfonden; postdoctoral funding to P.L.M. from StratNeuro and a NARSAD Young Investigator Grant (Brain & Behavior Research Foundation).

## Author contributions

D.C. performed neuroanatomical, behavioral, optogenetic experiments, and data analysis.

J.F. performed electrophysiology and Patch-seq, and data analysis.

P.L.M performed in vivo PFC recordings and data analysis.

D.C., J.F., and P.L.M contributed to the study design.

M.S. performed data analysis and visualization of PFC recordings.

F.J. performed in vivo LHA recordings and data analysis.

C.O. contributed with data analysis and visualization of neuroanatomy.

A.L. contributed with data analysis and visualization from Patch-seq experiments.

V.C. contributed to electrophysiology and data analysis.

M.G. contributed to behavioral experiments.

I.N. contributed to behavioral experiments and neuroanatomy.

M.W. contributed with behavioral experiments.

I.La. contributed with neuroanatomical and behavioral experiments.

H.K. contributed with data analysis and visualization of neuroanatomy.

I.Le. contributed with neuroanatomical and behavioral experiments.

H.P. contributed with neuroanatomical and behavioral experiments.

B.R. contributed with RNA sequencing experiments and data analysis.

M.C. supervised the PFC experiments.

K.M. designed the study and supervised the project.

K.M. wrote the manuscript with help from D.C., J.F, P.L.M. and M.C.

All authors discussed and commented the manuscript.

## Data availability

The Neuropixels dataset generated during this study will be available in a NWB standard format at [Dandy Archive].

## Code availability

The code generated during this study to analyze the Neuropixels dataset will be available at: https://github.com/PierreLeMerre/Esr1_NPX_code

**Extended Data Video 1.**

**Related to figure 4. Real time place aversion (rtPA) in Esr1-cre mice.**

Real time place aversion induced by optogenetic stimulation of the Esr1+ LHA-LHb pathway.

**Extended Data Video 2.**

**Open field behavior of Npy-cre mice.**

Rearing events in the open field induced by optogenetic stimulation of the Npy+ LHA-LHb pathway.

**Extended Data Video 3.**

**Neuropixels recordings of the aversive state.**

Head-fixed internal vs external aversive state conditioning during Neuropixels recording in mPFC.

**Extended Data Video 4. Pupil tracking.**

Tracking pupil with DeepLabCut. Same Esr1-cre mouse as in Video 3.

**Extended Data Video 5. Looming stimulus presentation in the 8-shaped arena.**

Behavioral response to looming stimuli in a female mouse 24h after stress induction.

---

**Extended Data Fig. 1.**
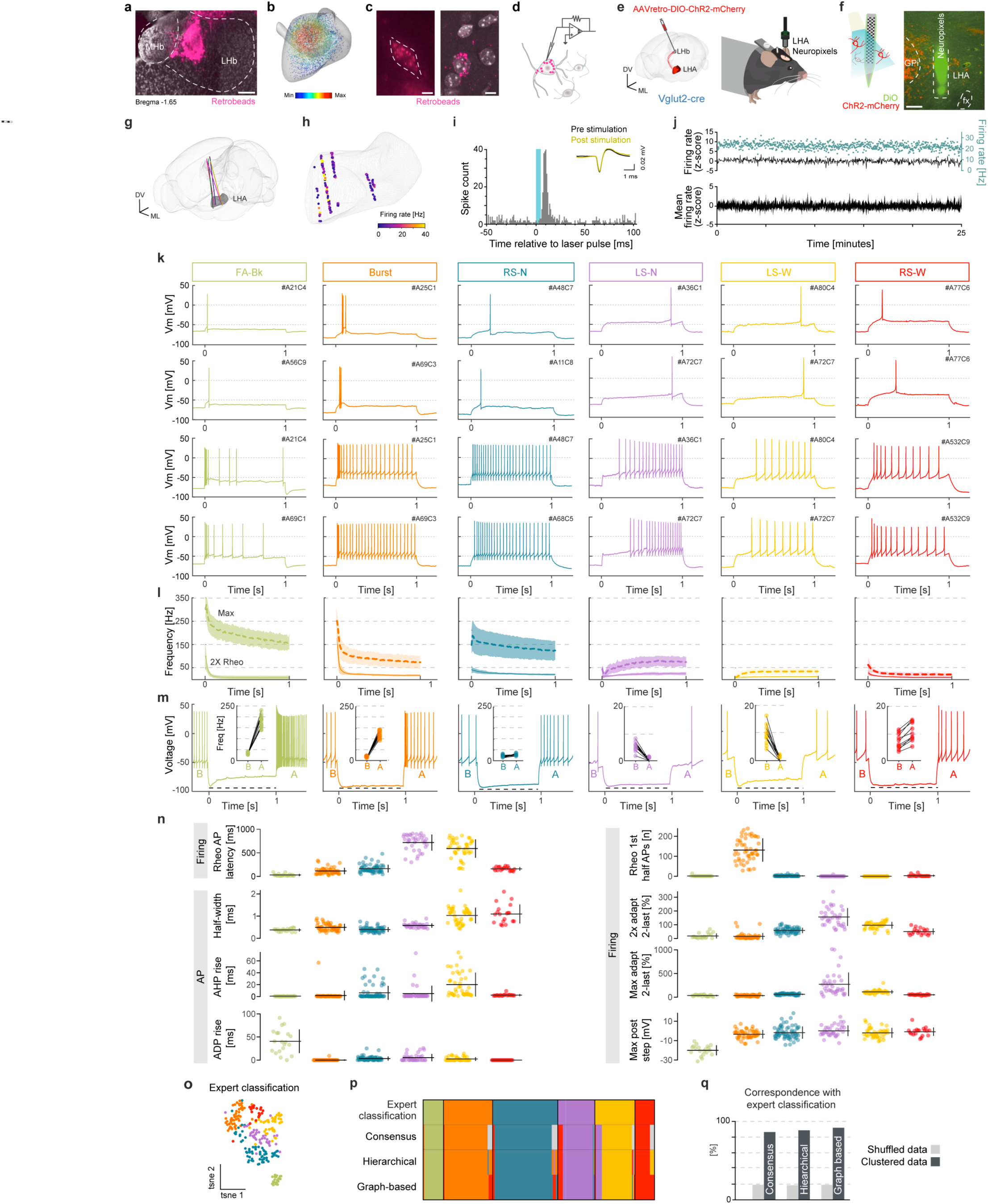
*In vivo* and *ex vivo* electrophysiology of LHA-LHb neurons. **(a)** Representative image of retrobeads injected into the LHb. **(b)** Volume-density distribution of retrobead labelled LHA-LHb neurons (n = 5883, N = 7 wt). **(c)** Representative images of retrobeads (pink) in retrogradely labelled LHA-LHb neurons. Left: epifluorescence (patch-clamp set-up), right: confocal. **(d)** Schematic of patch-clamp recordings of retrobead-labelled LHA-LHb neurons. **(e)** Experimental design for opto-tagging (Neuropixels) of LHA-LHb neurons in awake head-fixed mice. **(f)** Left: schematic illustration of Neuropixels probe dyed with DiO (green), LHA-LHb neurons expressing ChR2-mCherry (red), and light application (light blue). Right: image of DiO labelled probe track (green) and ChR2-mCherry expressing neurons (red). **(g)** 3D rendering of the tracked anatomical position of five Neuropixels probes in the LHA. **(h)** The spontaneous firing frequency (color coded) of recorded LHA-LHb units (n = 165 units; N = 5 Vglut2-cre mice). **(i)** Peri-stimulus histogram for an example light-activated (opto-tagged) LHA-LHb unit relative to light stimulus (blue). Inset: spontaneous (black) and light-evoked (yellow) spike waveforms. **(j)** Top: 25 min recording of an example opto-tagged, tonically active LHA-LHb unit. Turquoise: raw firing rate, black: z-scored firing rate. Bottom: average, z-scored firing rates of opto-tagged, tonically active LHA-LHb units (n = 9, N = 9 Vglut2-cre mice). **(k)** Example firing patterns (current clamp) of the six identified LHA-LHb neuron types. Upper three rows: response at rheobase; lower three rows: response at double rheobase. **(l)** Average of inter-spike interval (ISI) immediate frequencies from whole-cell firing patterns at 2X Rheo and at Max (mean ± SD). Same neurons as in **Fig 1b**. **(m)** Example responses to hyperpolarizing current injections (n = 8 for each neuron type). Insert: firing frequency as the average of the 200 ms firing before (B) and after (A) hyperpolarization. Dashed horizontal line: hyperpolarization activated depolarization or “Sag-potential”. Same neurons as in **Fig 1b**. **(n)** Hive plots of AP- and firing-related parameters for the six identified LHA-LHb neuron types (mean ± SD, n = 230 neurons, N = 46 mice). **(o)** t-SNE plot of LHA-LHb neuron types classified by electrophysiology. Color code: expert classification. **(p)** Comparison of neuronal identity by expert classification, hierarchical clustering, graph-based clustering, and consensus clustering. Gray: lack of agreement. Consensus clustering: the agreement between hierarchical and graph-based unbiased analyses. **(q)** Correspondence of the three clustering approaches (hierarchical clustering, graph-based clustering, and consensus clustering) and the randomly shuffled data of each clustering with the expert classification. Abbreviations: Globus pallidus internus (GPi), fornix (fx), All data acquired in male mice. Scale bars: 100 µm (**a)**, 10 µm (**c**), 150 µm (**f**).

**Extended Data Fig. 2.**
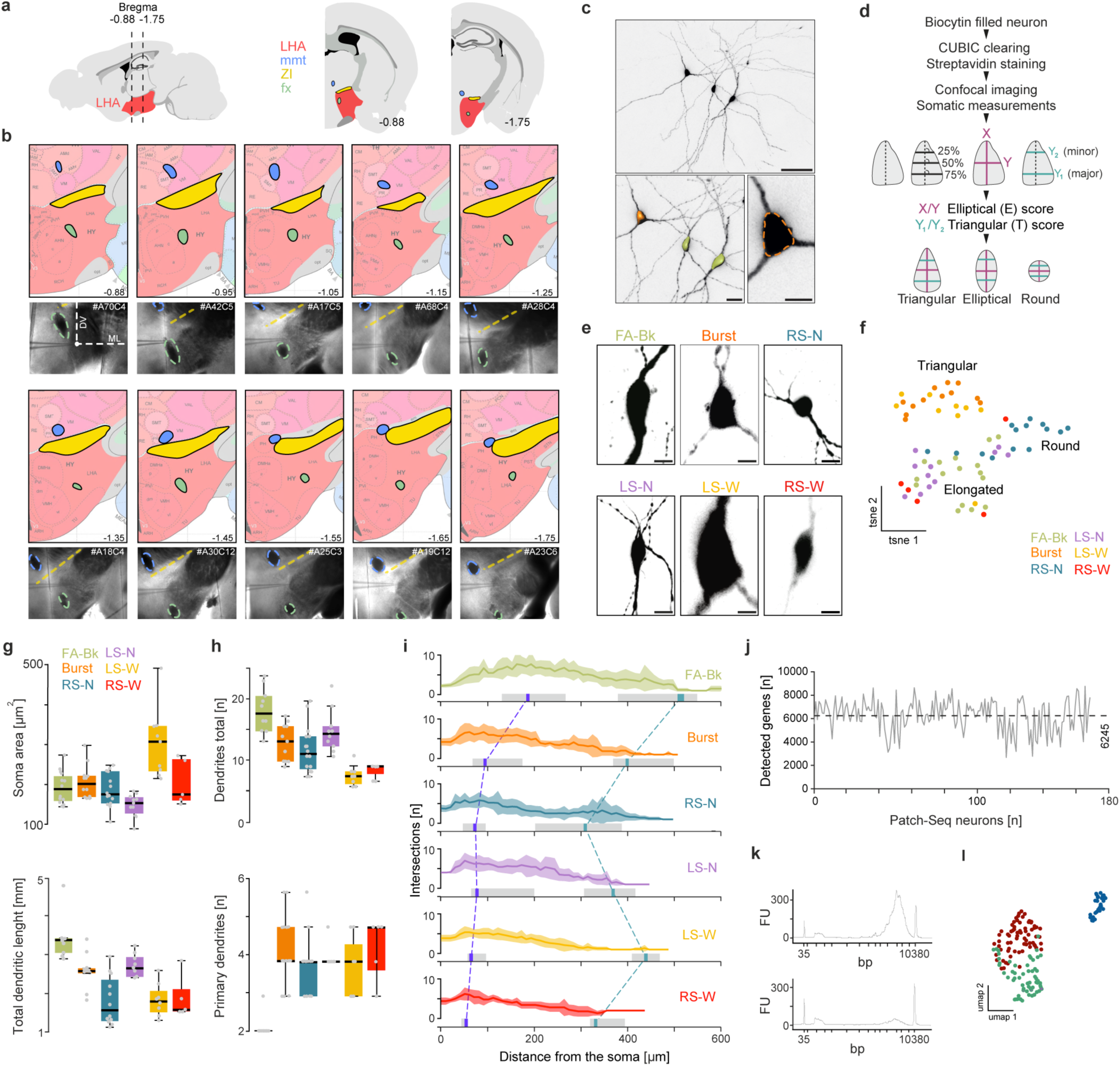
Anatomical, morphological, and genetic characterization of electrophysiologically recorded LHA-LHb neurons. **(a)** Left: AP range of tissue used for whole-cell patch-clamp recordings. Right: landmarks in the vicinity of the LHA used for mapping of the AP origin of coronal sections used for whole-cell patch-clamp recordings of LHA-LHb neurons: mammillary tract (blue), fornix (green), zona incerta (yellow). **(b)** Top: coronal anatomical plates (Allen Brain Atlas CCFv2) at different AP coordinates. Bottom: DIC image of a corresponding *ex vivo* brain slice and a recording electrode. The relative DV and ML position of mmt, fx, and ZI was used to identify the AP position of the brain slice. **(c)** Top: confocal image of three representative biocytin filled LHA-LHb neurons. Bottom left: magnification of the neurons, with pseudo-coloring of the soma. Bottom right: the soma outlines (dashed line) were detected in the orthogonal projection of confocal z-stack. **(d)** Outline of the workflow for soma morphometry. The elliptical score (X/Y ratio, magenta lines) and the triangular score (Y1/Y2 ratio, turquoise lines) were used for classification of soma as triangular, elongated, or round, respectively. **(e)** Representative soma shapes (biocytin filled) of the six identified LHA-LHb neuron types. **(f)** t-SNE plot of the parameters established by soma morphometry. Color code: expert classification. FA-Bk: n = 16, Burst: n = 12, RS-N: n = 14, LS-N: n = 11, LS-W: n = 9, RS-W: n = 5 LHA-LHb neurons, N = 17 wt mice. **(g)** Quantification of the soma area (top, same neurons as in (**f**)) and total dendritic length (bottom, FA-Bk: n = 8, Burst: n = 10, RS-N: n = 12, LS-N: n = 8, LS-W: n = 8, RS-W: n = 5. N = 19 wt mice). **(h)** Quantification of the total number of dendrites (top) and the number of primary dendrites (bottom). FA-Bk: n = 10, Burst: n = 11, RS-N: n = 16, LS-N: n = 10, LS-W: n = 8, RS-W: n = 5. **(i)** Sholl analysis (number of intersections) of reconstructed neurons highlights the cell-type specific complexity of the dendritic trees of LHA-LHb neuron types, mean ± s.e.m. Violet vertical bar: median of the critical radius, turquoise vertical bar: median of the enclosing radius, gray horizontal bar: 95 percentile. Same neurons as in (**g**), bottom. **(j)** Number of detected genes in the Patch-seq LHA-LHb neurons (n = 163, same neurons as in **Fig 2b-d**). Average: 6245 genes/neuron. **(k)** Representative examples of Agilent Bioanalyzer traces of cDNA amplicons of mRNA samples from single Patch-seq harvested LHA-LHb neurons. Top: successful sample (included in data), bottom: failed sample (discarded), ∼10% of samples were discarded. **(l)** UMAP plot showing identity of neurons based on gene expression from Patch-seq, identifying three clusters of LHA-LHb neurons, n = 163, same neurons as in **Fig 2b-d**, color code as in **Fig 2b**, right. All data acquired in male mice. For boxplots (**g**-**h**), data shown as median, box (25th and 75th percentiles), and whiskers (data points that are not outliers). Scale bars: 50 µm (**a**, top), 25 µm (**a**, bottom left), 10 µm (**a**, bottom right). Abbreviations: mammillary tract (mmt), zona incerta (ZI), fornix (fx), #A70C4 (along all S2B) codes for patched cell ID: A-(animal number) and C-(cell number), and Uniform Manifold Approximation and Projection (UMAP)

**Extended Data Fig. 3.**
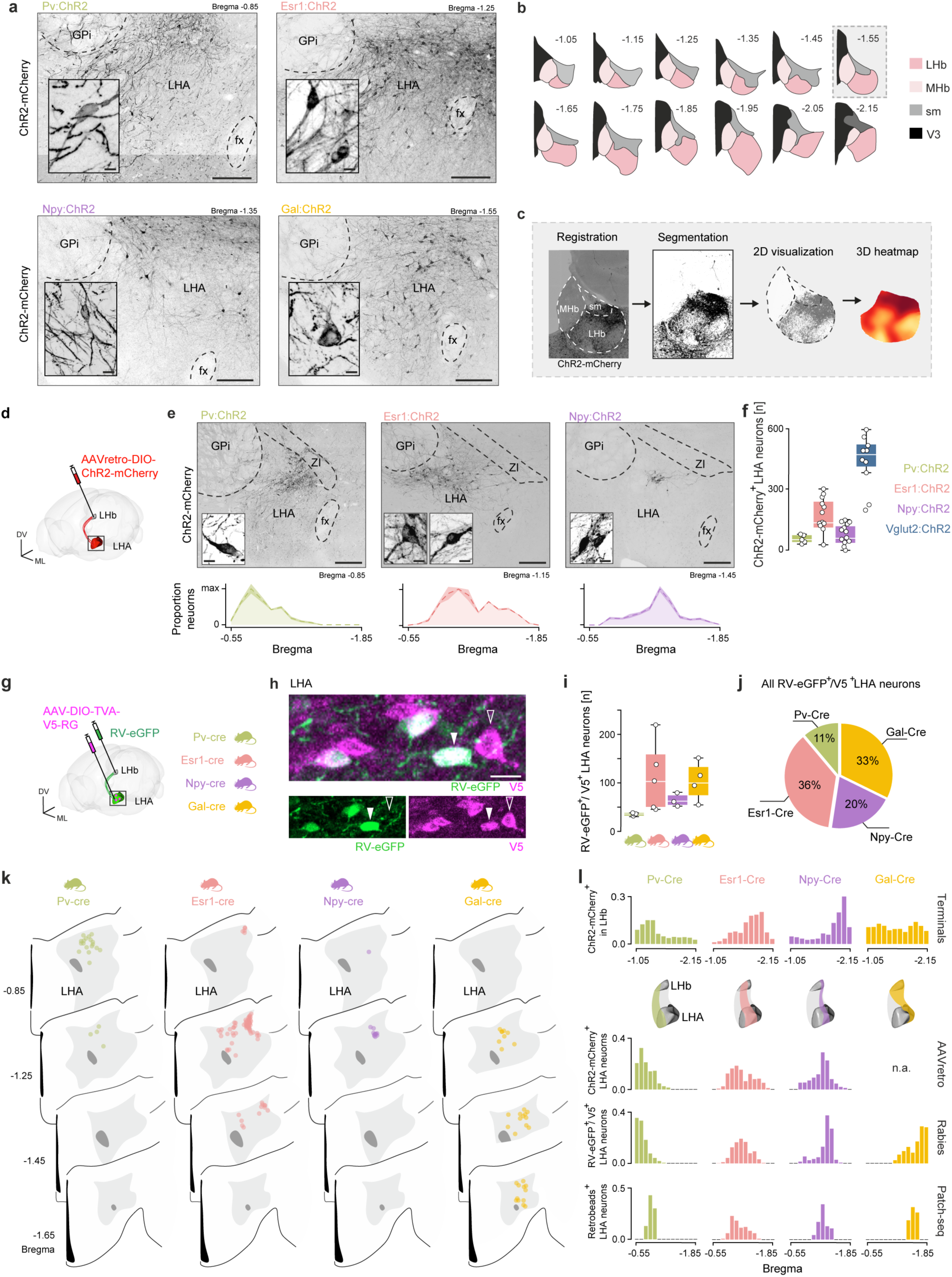
Genetic targeting of LHA-LHb pathways. **(a)** Representative confocal images of cre-dependent anterograde labelling of LHA-LHb neurons in Pv-cre, Esr1-cre, Npy-cre, and Gal-cre mice, respectively. Inserts: representative examples of cell-type specific soma morphologies. **(b)** Anatomical plates (Allen Brain Atlas CCFv2) of the LHb and neighboring structures along the AP range. **(c)** Outline of the workflow for anatomical mapping of axon terminals. **(d)** Experimental design for cell-type specific retrograde labeling of LHA-LHb neurons. **(e)** Top: representative confocal images of retrogradely labelled neurons in the LHA. Insert: representative examples of cell-type specific soma morphologies. Bottom: quantification of the labelled soma along the AP axis (mean ± s.e.m; Pv:ChR2: N = 5, Esr1:ChR2: N = 12, Npy:ChR2: N = 13). **(f)** Quantification of retrogradely labelled LHA-LHb neurons (same mice as in (**e**) + Vglut2:ChR2: N = 10. **(g)** Experimental design for rabies tracing. The helper virus AAV-DIO-TVA-V5-RG was injected in the LHA, and the EnvA-coated rabies virus (RV-eGFP) in the LHb. **(h)** Representative confocal image of LHA-LHb neurons co-expressing RV-eGFP (green) and V5 (magenta; Esr1-cre mouse). Arrow head: RV-eGFP+/V5+ neuron (quantified), empty arrow head: RV-eGFP-/V5+ neuron (not quantified). **(i, j)** Quantification of RV-eGFP+/V5+ LHA-LHb neurons Pv-cre, Gal-cre: N = 4, Esr1-cre: N = 5, Npy-cre: N = 3. (**i**) total number, (**j)** the proportion (average) RV-eGFP+/V5+ LHA neurons in each genotype, of all RV-eGFP+/V5+ labelled LHA neurons detected across the genotypes. Pv-cre, Gal-cre: N = 4; Npy-cre: N = 3; Esr1-cre: N = 5 mice. **(k)** Anatomical plates (Allen Brain Atlas CCFv2) of the LHA along the AP axis with the position of detected RV-eGFP+/V5+ LHA-LHb neurons. One dot = one neuron. Each plate depicts neurons detected in two consecutive brain sections (50 µm) from a representative mouse. **(l)** Topography of the projection field and the soma location of the LHA-LHb pathways along the AP axis. From top to bottom: bar plots with the proportion axon terminals in the LHb (anterograde viral strategy, Pv-cre: N = 3, Esr1-cre: N = 5, Npy-cre: N = 4, Gal-cre: N = 4 mice); soma in the LHA (retrograde viral strategy, Pv-cre: N = 5, Esr1-cre: N = 12, Npy-cre: N = 13; soma in the LHA (rabies tracing, Pv-cre: N = 4, Esr1-cre: N = 5, Npy-cre: N = 3, Gal-cre: N = 4; neurons in the LHA (Patch-seq, FA-Bk: n = 12, Burst + RS-N: n = 77, LS-N: n = 35, LS-W: n = 28). Abbreviations: Globus pallidus internal (GPi), fornix (fx), medial habenula (MHb) and zona incerta (ZI), medial habenula (MHb), stria medullaris (sm), and third ventricle (V3). For boxplots (**f, i**), data shown as median, box (25th and 75th percentiles), and whiskers (data points that are not outliers). Scale bars: 200 µm (**a, e)**, inset 10 µm (**a, e**), 10 µm (**h**).

**Extended Data Fig. 4.**
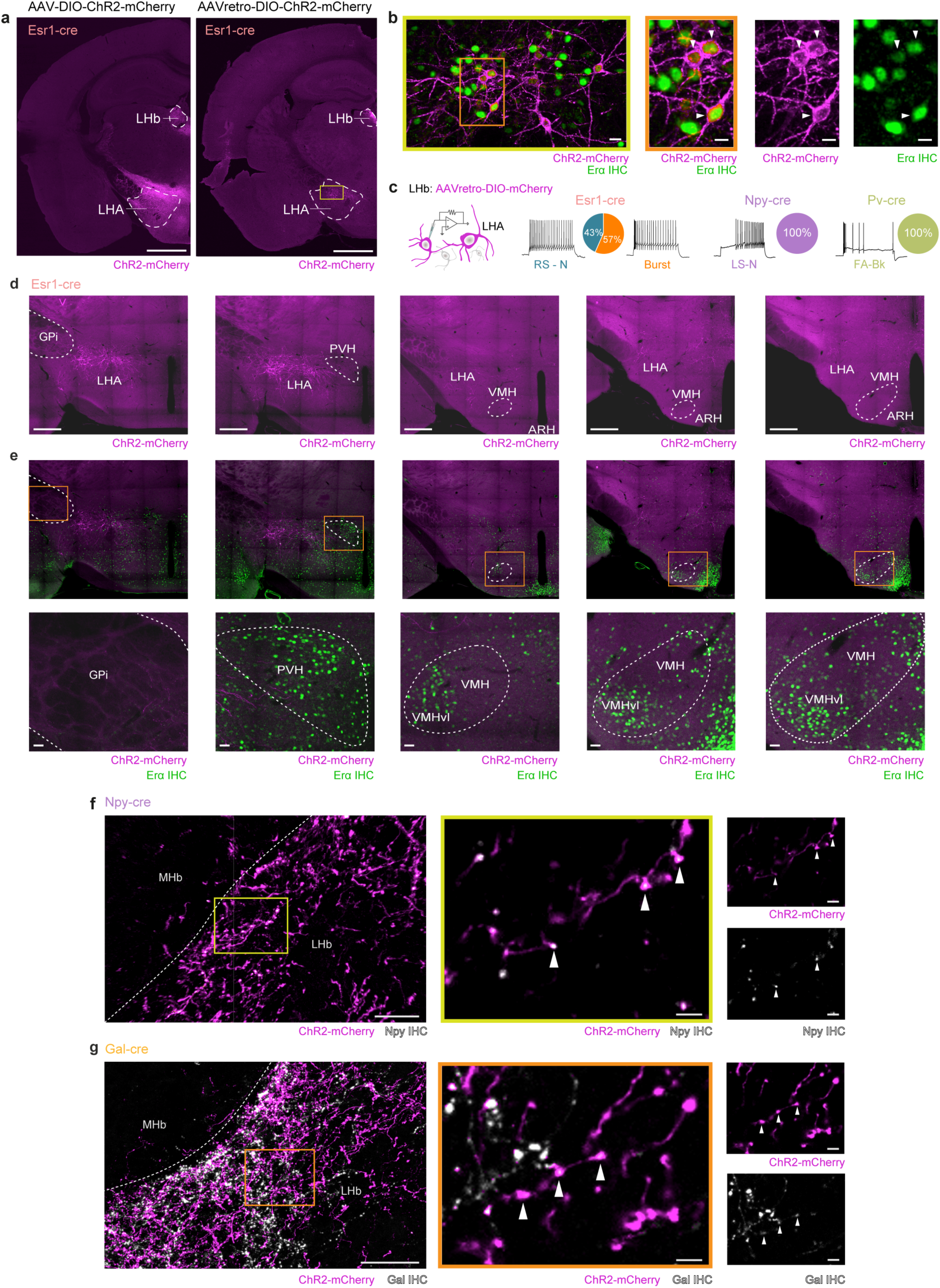
Characterization of distinct LHA-LHb neuron types. **(a)** Anterograde (left) and retrograde (right) viral labeling of the Esr1+ LHA-LHb pathway in the Esr1-cre mice. **(b)** ESR1 protein expression (green) was confirmed by IHC in retrogradely labeled LHA-LHb neurons (magenta) in Esr1-cre mice. Orange box: location of right panel. **(c)** Retrograde viral labeling of Esr1+ LHA-LHb neurons with consecutive whole-cell patch-clamp characterization of labeled neurons exclusively identified RS-N (n = 10) and Burst (n = 13) type neurons, corroborating the detection of Esr1 in the Patch-seq transcriptome of these neuron types. N = 4 Esr1-Cre mice. Retrograde viral labeling of the LHA-LHb pathway in Npy-cre and Pv-cre mice, respectively, with consecutive whole-cell patch-clamp characterization of labeled neurons exclusively identified LS-N type LHA-LHb neurons (n = 19) in Npy-cre mice (N = 3), and FA-Bk type LHA-LHb neurons (n = 12) in PV-cre mice (N = 3), confirming the validity of the experimental strategy for cell type specific targeting of LHA-LHb pathways. **(d)** Representative confocal images along the AP axis (left to right) of the hypothalamic area with retrograde viral labeling (magenta) of the LHA-LHb pathway in Esr1-cre mice. **(e)** Top: same images as in (**d**), here with detection of ESRα protein expression (IHC; green). Orange box: location of panel in bottom row. Bottom: close-up of box in top. **(f-g)** Left: representative confocal images of virally labelled (magenta) LHA-LHb axon terminals in the LHb of a Npy-cre mouse (**f**) and a Gal-cre mouse (**g**), confirming expression of the neuropeptide Y (white) in the LHA-LHb axon terminals in Npy-cre mice, and expression of galanin (white) in the LHA-LHb axon terminals in Gal-cre mice. Yellow/orange box: location of right panel. Right: close-up of box in left. All data aquired in male mice. Scale bars: 1 mm (**a**), 10 µm (**b**, **f**, left), 200 µm (**d**), 50 µm (**e**, bottom), 2 µm (**f**, right). Abbreviations: Globus pallidus internal (GPi), paraventricular hypothalamic nucleus (PVH), ventromedial hypothalamic nucleus (VMH), ventromedial hypothalamic nucleus, ventrolateral part (VMHvl).

**Extended Data Fig. 5.**
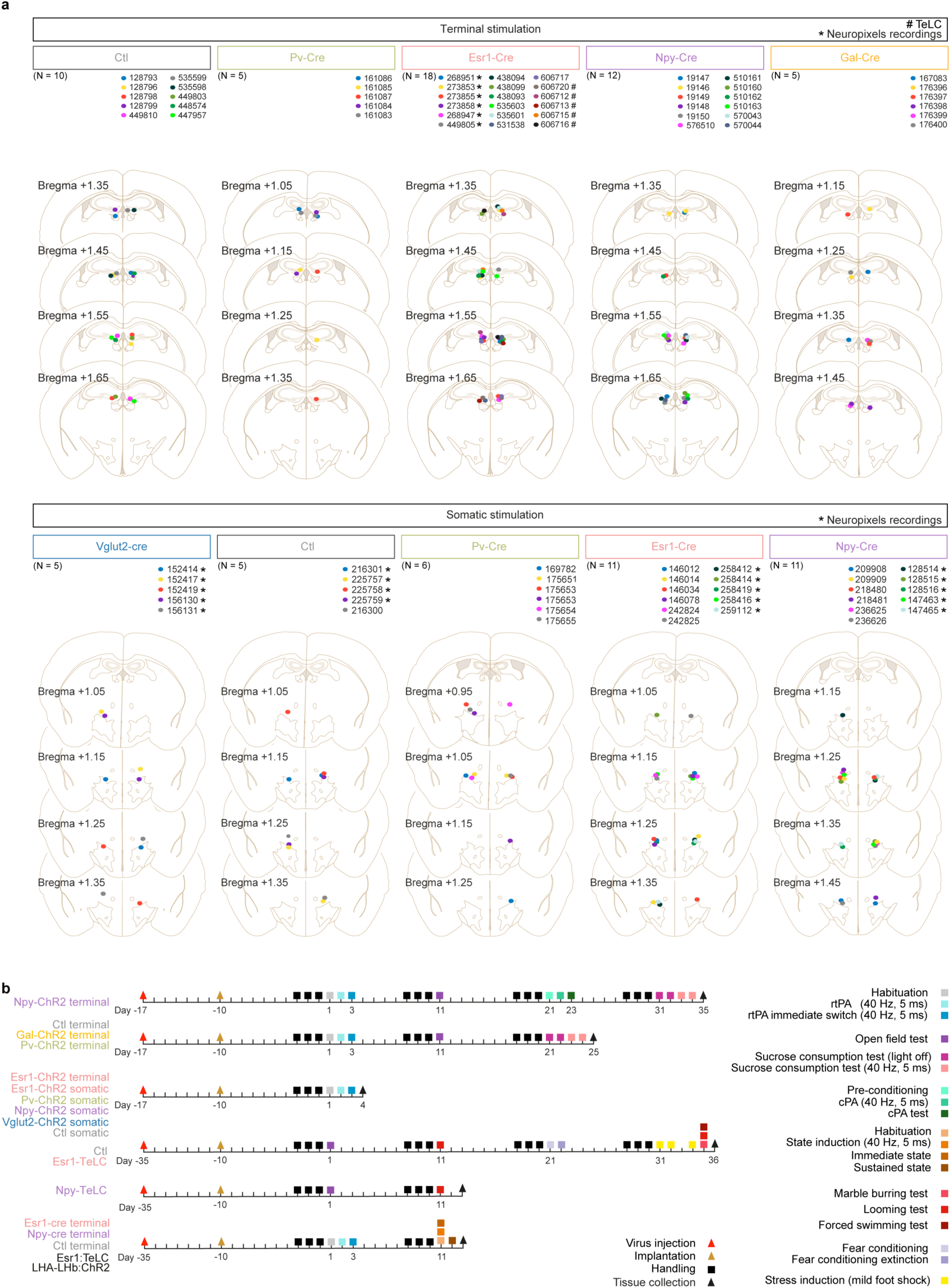
Confirmation of the position of optical fibers, and overview of experimental time lines. **(a)** The location of the tip of the optical fibers across the experiments, visualized across the AP axis of coronal anatomical plates (Allen Brain Atlas CCFv2). Top: position of the tip of optical fibers in experiments with optogenetic stimulation of LHA-LHb axon terminals in the LHb. Bottom: position of the tip of optical fibers in experiments with somatic optogenetic stimulation of LHA-LHb neurons in the LHA. Colored dots: individual mice and their respective bilateral optical fiber location. *) mice subjected to Neuropixels recordings, #) mice subjected to TeLC silencing. **(b)** Schematic overview of the experimental time line for the different cohorts of mice. N = number of mice.

**Extended Data Fig. 6.**
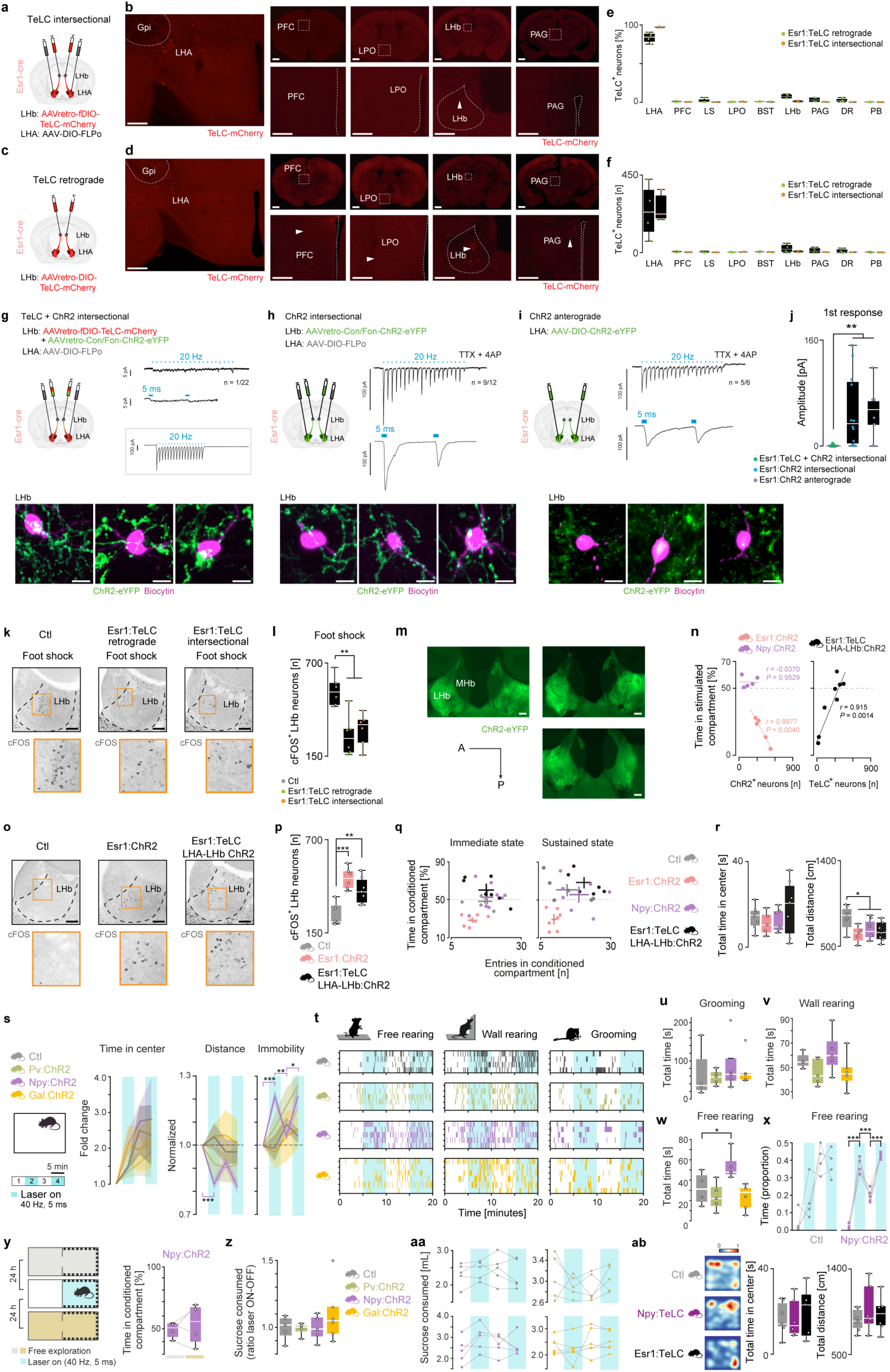
Validation of the TeLC silencing approaches, and optogenetic manipulation or TeLC silencing of LHA-LHb pathways during behavior. **(a)** Experimental design of the intersectional approach for TeLC silencing of LHA-LHb neurons. (**b**) Representative confocal images of TeLC labelled LHb projecting neurons (red) in the LHA and potential off-target brain regions in a Esr1-Cre mouse for the TeLC intersectional approach. (c) Experimental design of the retrograde approach for TeLC silencing of Esr1+ LHA-LHb neurons. (**d**) Representative confocal images of TeLC labelled neuron (red) **(e-f)** Quantification of neurons expressing TeLC-mCherry in the LHA and putative off-target regions in Esr1-cre mice. (**e**) The distribution of TeLC-mCherry+ neurons in different brain regions, plotted as the percentage of the total number of TeLC-mCherry+ neurons in individual mice. (**f**) The number of TeLC-mCherry+ neurons in different brain regions. Green dots: mice with TeLC retrograde approach (N = 4), orange dots: mice with TeLC intersectional approach (N = 3). **(g)** Electrophysiological *ex vivo* validation of the silencing of Esr1+ LHA-LHb neurons. Left: experimental design of the intersectional viral approach for concurrent TeLC silencing and ChR2-expression in Esr1+ LHA-LHb neurons. Right: whole-cell recordings of LHb neurons concurrent with light application (1 s, 5 ms pulses, 20 Hz) in the LHb validated the absence of light-evoked synaptic responses upon TeLC silencing of Esr1+ LHA-LHb neurons. Top: representative light-evoked response in a LHb neuron (synaptic response in 1/22 LHb neurons). Middle: close-up of trace in top, showing the two first responses to light stimulation. Box: Representative light-evoked response of a TeLC+/ChR2+ neuron in the LHA. Bottom images: representative confocal images of whole-cell recorded biocytin (magenta) filled LHb neurons intermingled with ChR2-eYPF+ (green) Esr1+ LHA-LHb axon terminals. **(h)** Left: experimental design of the intersectional viral approach for expression of ChR2 in Esr1+ LHA-LHb axon terminals. Right: whole-cell recordings of LHb neurons concurrent with light application (1 s, 5 ms pulses, 20 Hz) in the LHb confirmed reliable synaptic responses in LHb neurons, validating the experimental strategy. Top: representative light-evoked response in a LHb neuron (synaptic response in 9/12 LHb neurons). Bottom: close-up of trace in top, showing the two first responses to light stimulation. Bottom images: representative confocal images of whole-cell recorded biocytin (magenta) filled LHb neurons intermingled with ChR2-eYPF+ (green) Esr1+ LHA-LHb axon terminals. **(i)** Left: experimental design of the anterograde viral approach for expression of ChR2 in Esr1+ LHA-LHb axon terminals. Right: whole-cell recordings of LHb neurons concurrent with light application (1 s, 5 ms pulses, 20 Hz) in the LHb confirmed reliable synaptic reponses in LHb neurons, validating the experimental strategy. Top: representative light-evoked response in a LHb neuron (synaptic response in 5/6 LHb neurons). Bottom: close-up of trace in top, showing the two first responses to light stimulation. Bottom images: representative confocal images of whole-cell recorded biocytin (magenta) filled LHb neurons intermingled with ChR2-eYPF+ (green) Esr1+ LHA-LHb axon terminals. **(j)** Average amplitude of the first light-evoked synaptic response for all the whole-cell recorded LHb neurons in (**g**-**i)**. TeLC + ChR2 intersectional vs ChR2 intersectional, p < 0.001; TeLC ChR2 intersectional vs ChR2 anterograde, p < 0.001, unpaired *t*-test. Colored dots: viral approach employed. Green dots: Esr1:TeLC + ChR2 intersectional: n = 22, blue dots: Esr1:ChR2 intersectional: n = 12, gray dots: Esr1:ChR2 anterograde: n = 6. **(k-l)** TeLC expression significantly reduced the expression of cFOS. Representative confocal images (**k**) and quantification (**l**) of cFOS+ neurons in the LHb in response to foot shock in control mice and mice with TeLC silencing of Esr1+ LHA-LHb neurons, respectively. Two different approaches were used for TeLC mediated silencing. (**l**) Ctl vs Esr1:TeLC retrograde, p = 0.0014; Ctl vs Esr1:TeLC intersectional, p = 0.0016, Unpaired *t*-test, N = 6 for each the three group of mice. **(m)** Representative confocal images of the LHb region along the AP axis, showing ChR2-eYFP+ (green) LHA-LHb axon terminals in a Esr1-cre mouse subjected to the TeLC + ChR2 intersectional approach. **(n)** The relationship (Pearson correlation coefficient) between the time spent (%) in the stimulated compartment in the real time place aversion test and the number of ChR2+ neurons (left), or the number of TeLC+ neurons (right). **(o-p)** Representative confocal images (**o**) and quantification (**p**) of cFOS+ neurons in the LHb in response to optogenetic stimulation of Esr1+ LHA-LHb axons terminal in mice with TeLC silening (Esr1:TeLC LHA-LHb ChR2 mice) or without TeLC silening (Esr1:ChR2 mice) of Esr+ LHA-LHb neurons. Ctl vs Esr1:ChR2, p < 0.001; Ctl vs Esr1:TeLC LHA-LHb ChR2, p = 0.0086, unpaired *t*-test. Ctl: N = 5, Esr1:ChR2: N = 6, Esr1:TeLC LHA-LHb ChR2: N = 7 mice. **(q)** Optogenetic activation of Esr1+ LHA-LHb axon terminals (Esr:ChR2 mice) reduces both the time spent and the number of entries into the conditioned compartment. Ctl, Esr1:ChR2, Esr1:TeLC LHA-LHb ChR2: N = 6, Npy:ChR2: N = 7 mice, same mice as in **Fig 3j-k**. **(r)** Open field. Continuous optogenetic activation (10 min, 40 Hz, 5 ms, 1 s on–1 s off) of LHA-LHb axon terminals significantly decreased the distance travelled in Esr1:ChR2, Npy:ChR2, and Esr1:TeLC LHA-LHb ChR2 mice compared to Ctl mice (right, Ctl vs Esr1:ChR2, p = 0.0207; Ctl vs Npy:ChR2, p = 0.0186; Ctl vs Esr1:TeLC LHA-LHb ChR2, p = 0.0389, unpaired *t*-test), while the time spent in the center was not affected (left). Same mice as in (**q**). **(s)** Open field. Block wise optogenetic activation (20 min total, 4 x 5 min blocks, 40 Hz, 5 ms) of the axon terminals of the Pv+, Npy+, or Gal+l LHA-LHb pathway, respectively, did not affect the time spent in the center of an open field. The time spent in the center in each block, plotted as fold change compared to block 1 (mean ± s.e.m). Optogenetic activation of the axon terminals of the Npy+ LHA-LHb pathway significantly decreased the distance travelled in the open field (Npy:ChR2 block 1 vs block 2, p < 0.001). This effect was also apparent as significantly increased immobility (block 2 vs1, p < 0.001; block 3 vs 2, p = 0.0043; block 4 vs 3, p = 0.038, t-test pair-wise comparison of each block to the previous). No significant effects on locomotion were observed in responses to activation of the Pv+ and Gal+ LHA-LHb pathways, respectively. Ctl: N = 4, Pv:ChR2, Npy:ChR2: N = 5, Gal:ChR2: N = 6 mice. **(t-w)** Scoring of free rearing, wall rearing, and grooming in the open field in response to block wise optogenetic activation of the axon terminals of the Pv+, Npy+, or Gal+ LHA-LHb pathway, respectively. Same animals as in (**s**). 1 colored coded vertical bar = 1 s in a specific behavior. (**w**) Optogenetic activation of the axon terminals of the Npy+ LHA-LHb pathway significantly increased the total time spent free rearing (Npy:ChR2 vs Ctl, p = 0.0487, same mice as in **Fig 3k**. **(x)** Quantification of proportion time free rearing in blocks with (blue) and without (white) light activation, respectively. Optogenetic activation of the axon terminals of the Npy+ LHA-LHb pathway significantly increased the proportion time spent free rearing (block 2 vs 1, p < 0.001; block 3 vs 2, p < 0.001; block 4 vs 3, p < 0.001, t-test pair-wise comparison of each block to the previous). Same mice as in **Fig 3k**. **(y)** Left: schematic outline of the experimental design for conditioned place aversion. Optogenetic activation (10 min) of the axon terminals of the Npy+ LHA-LHb pathway did not induce avoidance of the conditioned compartment (right). Gray horizontal bar: before conditioning, gold horizontal bar: after conditioning. N = 4 Npy:ChR2 mice. **(z-aa)** Sucrose consumption test, with block wise optogenetic activation (as in (**s**)). Optogenetic activation of the axon terminals of the Pv+, Npy+, or Gal+ LHA-LHb pathway, respectively, did not alter sucrose consumption. (**z**) The ratio of the sucrose consumed in blocks with (ON) vs blocks without (OFF) light application. (**aa**) Sucrose consumed in the individual blocks. Ctl, Pv:ChR2, Npy:ChR2, Gal:ChR2: N = 5 mice. **(ab)** TeLC silencing of Npy+ LHA-LHb neurons or Esr1+ LHA-LHb neurons did not alter behavior in the open field (TeLC intersectional approach, see panel (**a**)). Left: heatmaps of example locomotion (10 min). Right: quantification of total time in center, and total distance, respectively. Ctl: N = 5, Npy:TeLC, Esr1:TeLC: N = 6 mice. Abbreviations: real-time place aversion (rtPA), conditioned place aversion (cPA). All data acquired in male mice. For boxplots (**e-f**, **j**, **l**, **p**, **r**, **u-w**, **z**, **ab**), data shown as median, box (25th and 75th percentiles), and whiskers (data points that are not outliers). Scale bars: 1 mm (**b**, top, **d,** top), 200 µm (**b**, bottom, **d** bottom), 100 µm (**k**, **m**, **o**), 10 µm (**g**-**i**), * *p* < 0.05, \*\**p* < 0.01, \*\*\**p* < 0.001.

**Extended Data Fig. 7.**
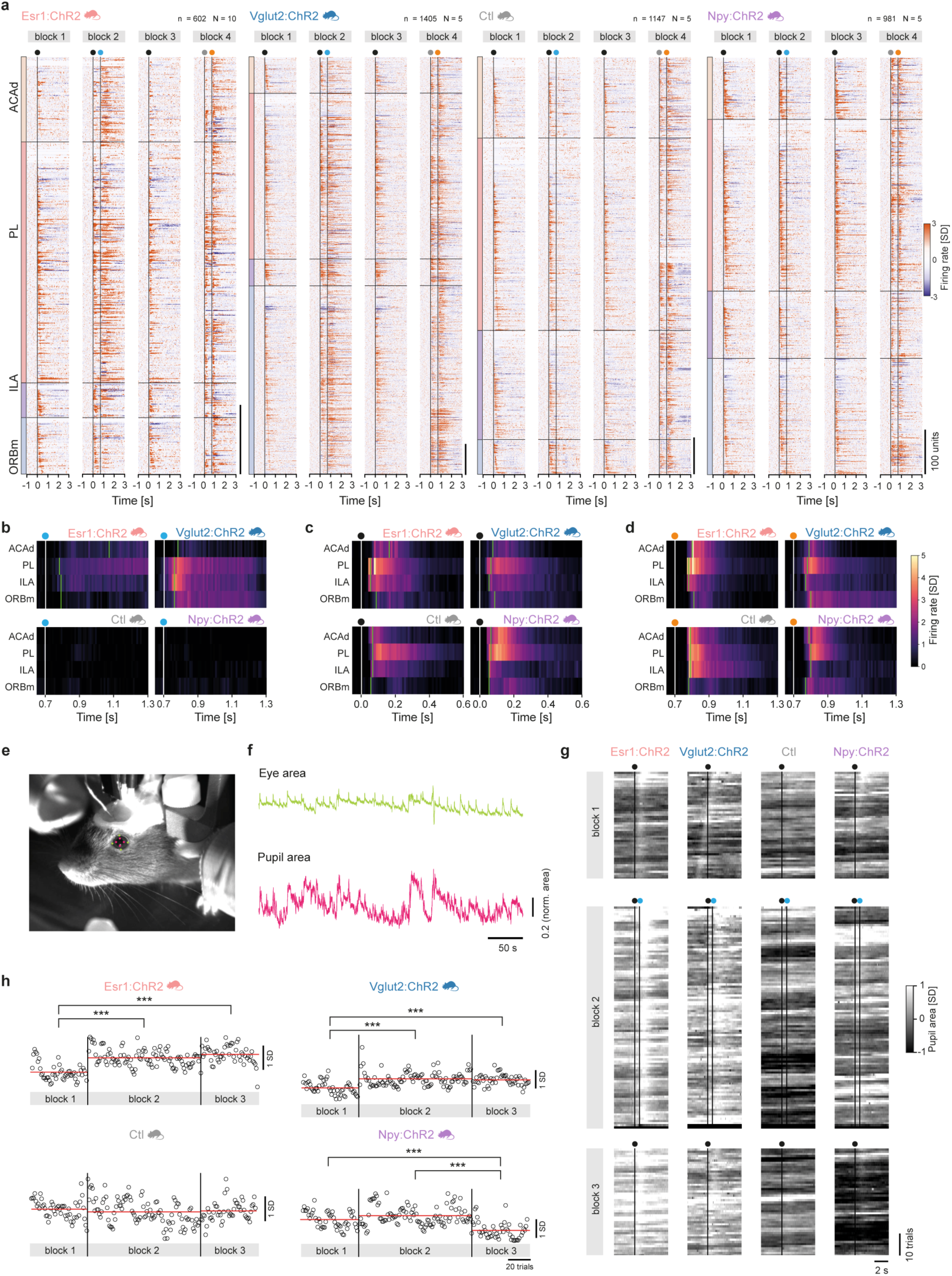
Pupil tracking, and state induction in the mPFC. **(a)** Trial-averaged firing rate (z-scored) for all individual units recorded across the mPFC subregions; Ctl: n = 1147, N = 5; Vglut2:ChR2: n = 1405, N = 5; Npy:ChR2: n = 981, N = 5, Esr1:ChR2: n = 602, N = 10). Colored dots: block specific events; black: sound 1, blue: optogenetic stimulation, gray: sound 2, orange: air puffs. The units are sorted based on their location within a specific mPFC subregion (outlined by horizontal black lines), and their respective peak response latency in block 1. **(b)** Average population firing across the mPFC subregions in (part of) block 2. The detected latency of the onset of the optogenetic responses is indicated with green vertical lines. White vertical line; onset of optogenetic stimulation. Bin size: 10 ms. **(c)** As (b), but detected latency of the onset of the sound response in block 1. White vertical line; sound onset. **(d)** As (b), but detected latency of the onset of the air puff response in block 4. White vertical line; air puff onset. **(e)** Example pupil tracking using DeepLabCut, light green dots: eye markers, pink dots: pupil markers. **(f)** Extracted traces of normalized eye area (top) and normalized pupil area (bottom), colors as in (e). **(g)** Heatmaps of the mean pupil area (z-scored to block 1) across the individual trials in block 1-3, Ctl, Vlut2:ChR2, Npy:ChR2: N = 5, Esr1:ChR2: N = 10 mice. Colored dots: block specific events; black: sound 1, blue: optogenetic stimulation. **(h)** Mean trial-averaged pupil area (z-scored to block 1) for all individual trials from block 1-3, same trials and mice as in (g). One circle = one trial, block1: n = 50, 2: n = 100, 3: n = 50 trials. Black vertical lines: block limits. Red horizontal lines: mean pupil area in the block. Mann-Whitney U test with Bonferroni correction. Abbreviations: medial prefrontal cortex (mPFC), anterior cingulate area, dorsal part (ACAd), prelimbic area (PL), infralimbic area (ILA) and orbitofrontal area, medial part (ORBm), peri-stimulus time histogram (PSTH), standard deviation (SD). n = number of units, N = number of animals. All data acquired in male mice. *** p < 0.001.

**Extended Data Fig. 8.**
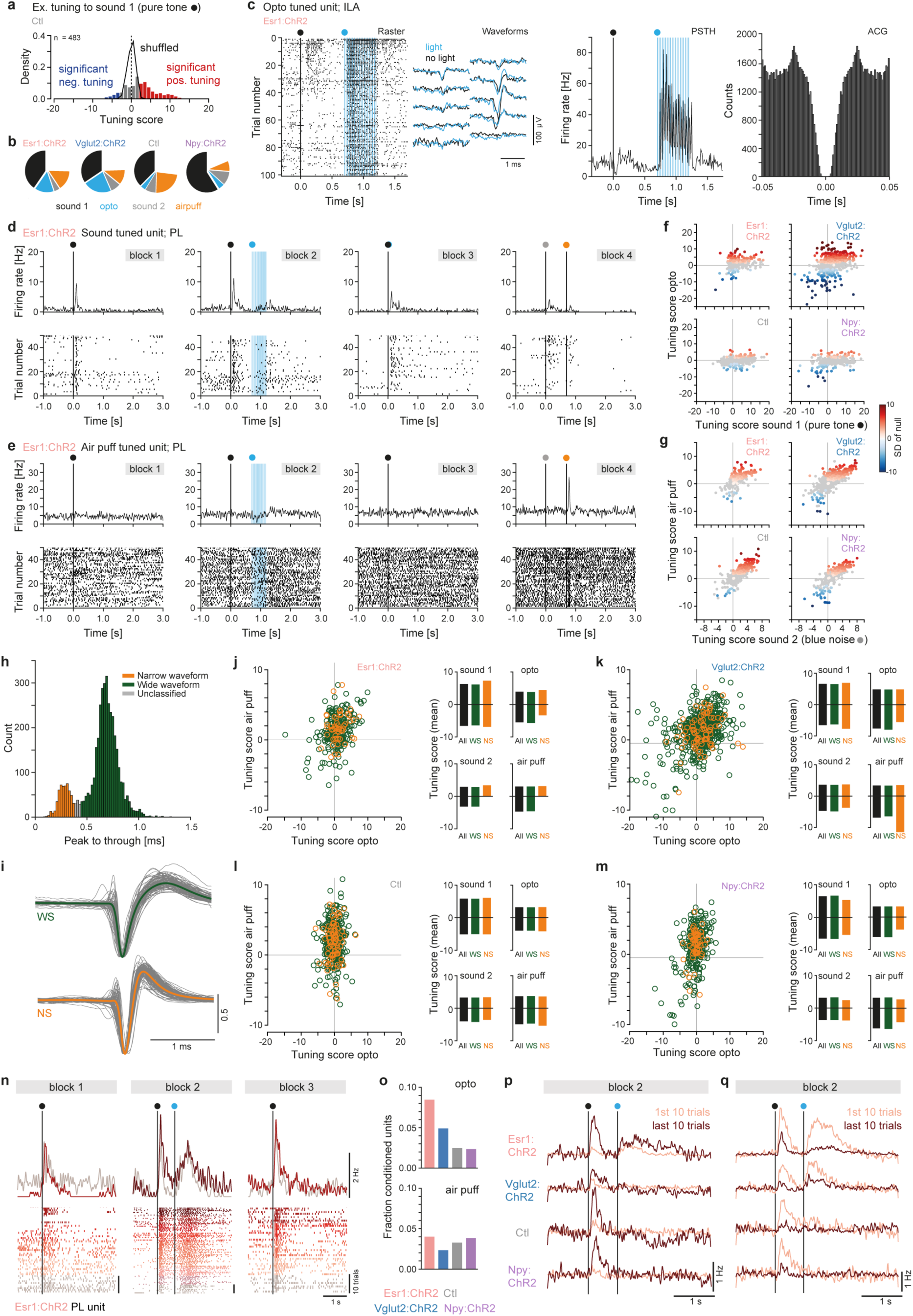
Tuning of mPFC units to sound, LHA-LHb pathway stimulation, and air puffs. (**a**) Tuning scores for all units in Ctl mice (N = 5) fitted with the GLM (n = 482). The tuning to sound 1 (pure tone) is shown. Black curve: tuning scores expected by chance. (**b**) Pie charts of the proportions of primary tuning for all units fitted with the GLM. Ctl: n = 482; Vglut2:ChR2: n = 738; Npy:ChR2: n = 406, Esr1:ChR2: n = 319 units. (**c**) Example ILA unit in an Esr1:ChR2 mouse, significantly tuned to the optogenetic stimulation. From left to right: raster plot aligned to the onset of the sound; mean waveform during (blue) or outside of (black) light application on 10 neighboring recording channels; peri-stimulus-time-histogram (PSTH) aligned to the onset of the sound; auto-correlogram (ACG; bin size: 1 ms). Colored dots: block specific events; black: sound 1, blue: optogenetic stimulation, blue vertical lines: light pulses (5 ms). (**d**) Example PL unit in an Esr1:ChR2 mouse, significantly tuned to sound 1 (pure tone; block 1-3). Top: PSTHs, bottom: raster plots of 50 trials. Colored dots: block specific events; black: sound 1, blue: optogenetic stimulation, gray: sound 2, orange: air puffs. (**e**) As **(d**) but for an example PL unit in an Esr1:ChR2 mouse significantly tuned to air puffs (block 4). (**f**) Scatterplots of the tuning of all units fitted with the GLM (n = 1954) in response to optogenetic stimulation (y axis) vs sound 1 (pure tone; y axis). One dot = one unit. Units significantly tuned to optogenetic stimulation are color coded (standard deviation from the null distribution). Gray units: non-significant tuning. (**g**) As **(f**) but for sound 2 (blue noise) vs air puffs. Units significantly tuned to air puffs are color coded (standard deviation from the null distribution). Gray dots: units without significant tuning. **(h**) Distribution of the peak to through duration for all recorded mPFC units (n = 4135, N = 25). Units with narrow waveform (peak to through < 0.38 ms) were classified as narrow spiking units (NS, orange) and units with wide waveform (peak to through duration > 0.44 ms) as wide spiking (WS, green) units. (**i**) Mean waveform of the WS (green trace) and NS (orange trace) units, respectively. Gray traces: waveforms of 100 randomly picked units/cell type. All waveforms are peak normalized. (**j-m**) Left: scatterplots of the tuning of all WS (green) and NS (orange) units fitted with the GLM (**Table 7** and **8**) in response to optogenetic stimulation (x axis) vs air puffs (y axis) for Esr1:ChR2 (**j**), Vglu2:ChR2 (**k**), Ctl (**l**), and Npy:ChR2 mice (**m**), respectively. One circle = one unit. Right: The mean tuning scores of the significantly tuned units, showing the mean negative and mean positive tuning scores, respectively. (**n**) Response profile of example PL unit in an Esr1:ChR2 mouse, across block 1-3. Top: PSTH, bottom: raster, both with color-coding of the trial progression within the blocks (lighter to darker, 10 trails/color). Colored dots: block specific events; black: sound 1, blue: optogenetic stimulation; vertical black lines: event onset. Bin size: 10 ms. (**o**) The fraction of conditioned units. Conditioned units displayed significantly (Wilcoxon signed rank-test) increased or decreased response to the sound and optogenetic manipulation in block 2 (top) or to the sound and air puffs in block 4 (bottom) over the trials within the respective block. (**p**) Mean response profile of the conditioned units that over the trials displayed significantly increased response to the sound in block 2. Mean of the first (light color) and last (dark color) 10 trials in block 2 are shown. Colored dots: block specific events; black: sound 1, blue: optogenetic stimulation, vertical black lines: event onset. Bin size: 10 ms. (**q**) As (**p**) but for the units that over the trials displayed significantly decreased response to the sound in block 2. n = number of units, Ctl, Vlut2:ChR2, Npy:ChR2: N = 5, Esr1:ChR2: N = 10 mice in all panels with group data. All data acquired in male mice.

**Extended Data Fig. 9.**
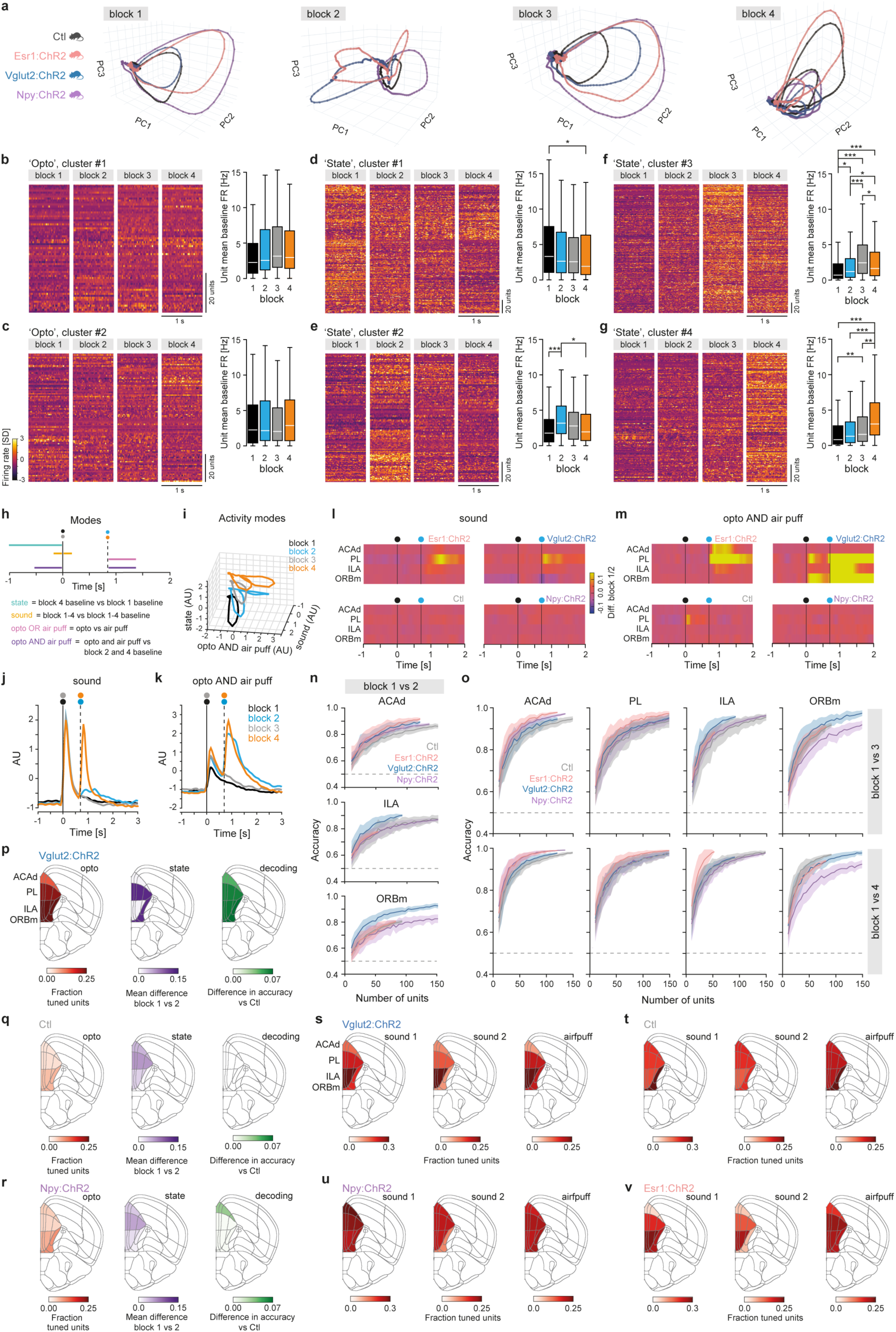
Prefrontal population dynamics in response to air puffs or optogenetic stimulation of distinct LHA-LHb pathways. (**a**) Trajectories of the neuronal population activity (trial averaged) in block 1-4; the first three principal components (PC) are shown. (**b-g**) Left: Trial-averaged baseline (the 1 s preceeding sound onset, n = 50 trials) firing rate (z-scored) in block 1-4 for all individual units in specific clusters (bin size 10 ms, see also **Fig 4h**). The units are sorted as in **Fig 4h**. Right: mean baseline firing rate in block 1-4, respectively. SD (error bar) and significance level (Wilcoxon signed-ranked test with Bonferroni correction). (**b**), ‘Opto’ cluster #1 (n = 65 units), (**c**) ‘Opto’ cluster #2 (n = 93 units), (**d**) ‘State’ cluster #1 (n = 195 units), (**e**) ‘State’ cluster #2 (n = 128 units), (b) ‘State’ cluster #3 (n = 253 units), (**g**) ‘State’ cluster #4 (n = 150 units). **h.** Schematic outline of the task events used to calculate the activity modes. **i.** mPFC neuronal population activity in the four blocks projected in 3D onto the three activity modes (state, sound, and aversive AND signal; N = 25 mice). **j.** The mPFC neuronal population activity in the four blocks (1-4, color coded, bin size: 10 ms) projected in 1D onto the Sound mode. Colored dots: block specific events; black: sound 1, blue: optogenetic stimulation, gray: sound 2, orange: air puffs. Vertical lines: event onset. See also **Fig 4h**. (k) as **(j**) but with projection on the ‘opto AND air puff’ mode. (l) Detailed investigation of the sound mode; involvement of mPFC subregions and LHA-LHb pathways. Difference of neuronal population activity between block 1 and 2, projected onto the sound mode; bin size: 50 ms. Colored dots: block specific events; black: sound 1, blue: optogenetic stimulation; Vertical lines: event onset. (m) as (**l**) but with projection onte the ‘aversive AND signal’ mode. (n) Decoding (average prediction accuracy) of block identity (block 1 vs 2) using baseline activity in the ACAd, ILA, and ORBm. Accuracy is plotted as a function of the number of units included in the logistic regression. Mean ± SD: 50 repeated cross-validations. Dashed line: 50% chance performance. (o) Top: decoding (average prediction accuracy) of block identity (block 1 vs 3) using baseline activity in the ACAd, PL, ILA, and ORBm. Accuracy is plotted as a function of the number of units included in the logistic regression. Mean ± SD: 50 repeated cross-validations. Dashed line: 50% chance performance. Bottom: as top, but block 1 vs 4. (p) Fraction of significantly tuned units across the mPFC subregions in response to optogenetic stimulation in Vlut2:ChR2 mice (left). Mean difference of the projection of the ‘state mode’ between block 1 vs block 2 across the mPFC subregions (middle). Difference in decoding (average prediction accuracy) of block identity (block 1 vs 2) in Vlut2:ChR2 vs Ctl mice (right). AP = 1.90 mm, Allen reference atlas CCFv3. (**q**-**r**) As (**p**), but for Ctl (**q**), and Npy:ChR2 mice (**r**). (**s**) Fraction of units across the mPFC subregions in Vglut2-cre mice with significant tuning to sound 1(left), sound 2 (middle), and air puffs (right) (**t-v**) As **(s**), but for Ctl (**t**), Npy:ChR2 (**u**), and Esr1:ChR2 (**v**) mice. n = number of units, N = number of animals. All data acquired in male mice. * *p* < 0.05, \*\**p* < 0.01, \*\*\**p* < 0.001.

**Extended Data Fig. 10.**
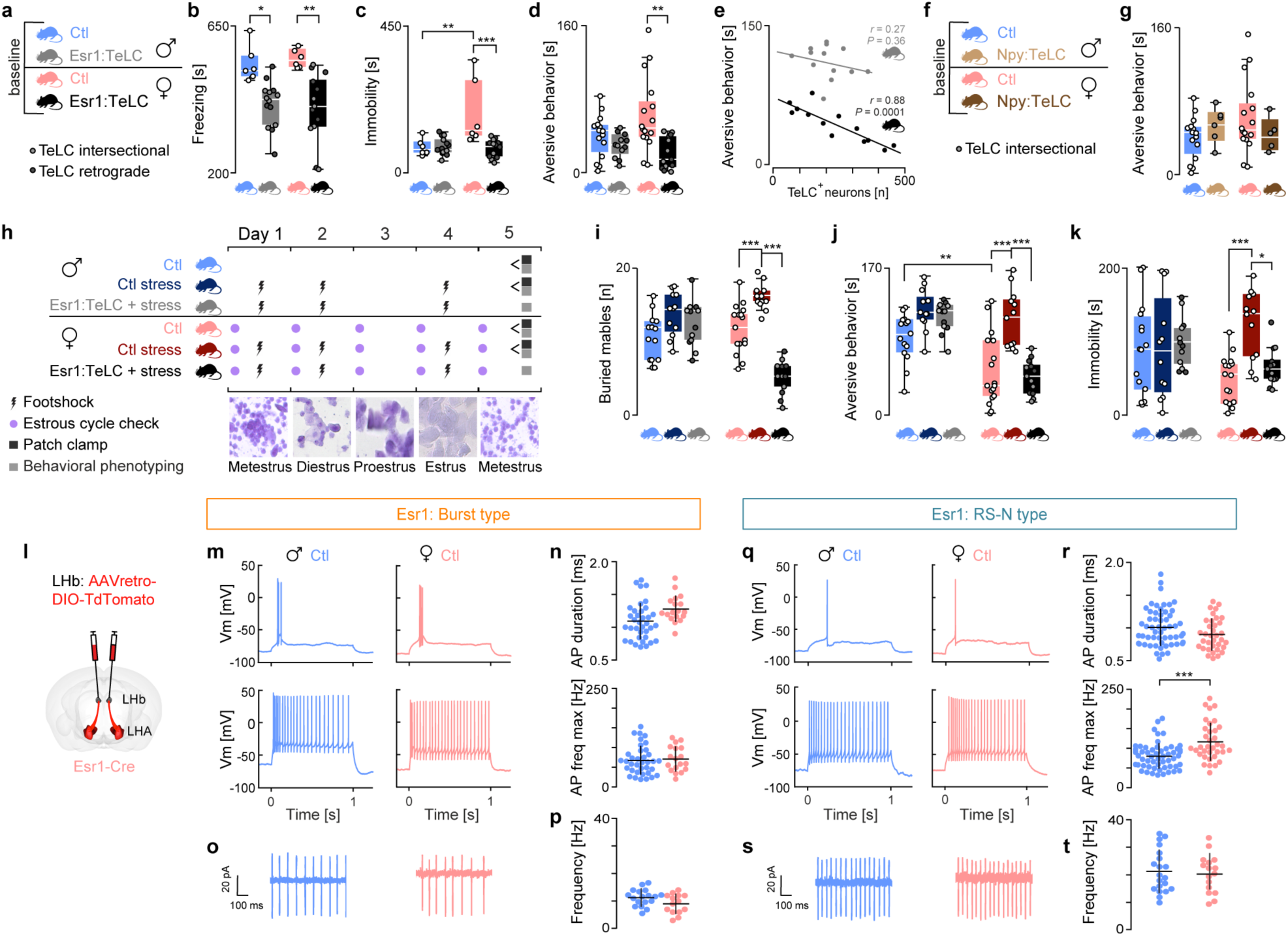
Aversive behaviors, behavioral stress response, and *ex vivo* electrophysiological characterization of baseline properties of Esr1 + LHA-LHb neurons in male and female mice. (**a**) Animal cohorts used in behavioral phenotyping (top), and the viral strategy for TeLC silencing of Esr1+ LHA-LHb neurons (bottom). (**b**) Fear conditioning. TeLC silencing of Esr1+ LHA-LHb neurons significantly reduced the total time freezing during fear conditioning extinction both in male and female mice (male Esr1:TeLC vs Ctl male, p = 0.0163; Esr1:TeLC female vs female Ctl, p = 0.0028, one-way ANOVA with Tukey’s Multiple comparisons test. Ctl: N = 6 male, 6 female, Esr1:TeLC: N = 14 male, 13 female. (**c**) Acoustic startle test. TeLC silencing of Esr1+ LHA-LHb neurons significantly reduced the immobility induced by tone presentation in female mice (one-way ANOVA with Tukey’s Multiple comparisons test, Ctl male vs Ctl female, p = 0.0029; Esr1:TeLC female vs Ctl female, p < 0.001. Same mice as in (**b**). (**d**) Looming stimuli test. TeLC silencing of Esr1+ LHA-LHb neurons in female mice significantly reduced the time spent in aversive behaviors (Ctl female vs Esr1:TeLC female, p = 0.0013, one-way ANOVA with Tukey’s Multiple comparisons test. Ctl: N = 15 male, 16 female, Esr1:TeLC: N = 14 male, 13 female. (**e**) The relationship (Pearson correlation coefficient) between the total time spent in aversive behavior in the looming stimuli test and the number of TeLC+ neurons. Same mice as in (**d**). (**f**) Animal cohorts used for baseline phenotyping in the looming stimuli test (top), and the viral strategy for TeLC silencing of Npy+ LHA-LHb neurons (bottom). (**g**) TeLC silencing of Npy+ LHA-LHb neurons did not influence the time spent in aversive behaviors in the looming stimuli test. Npy:TeLC: N = 6 male, 5 female, Ctl: N = 15 males, 16 female (same Ctl mice as in (**d**)). (**h**) Experimental design and animal cohorts used for behavioral phenotyping and *ex vivo* electrophysiological recordings. **(i-k)** Behavioral phenotyping 24 h after ending of the stress paradigm (see also **Fig 5c**). Ctl: N = 15 male, 16 female, Ctl stress: N = 12 male, 12 female; Esr1:TeLC stress: N = 12 male, 13 female; all mice same as in **Fig 5c-f**. One-way ANOVA with Tukey’s Multiple comparisons for all statistics. (**i**) Marble burying test: Ctl female vs Ctl stress female: p < 0.001; Ctl stress female vs Esr1:TeLC stress female: p < 0.001. (**j**) Looming stimuli test: Ctl male vs Ctl female: p = 0.0062; Ctl stress female vs Ctl female: p < 0.001; Ctl stress female vs Esr1:TeLC stress female: p < 0.001. (**k**) Forced swim test: Ctl stress female vs Ctl female: p < 0.001; Ctl stress female vs Esr1:TeLC stress female: p = 0.0254. (**l**) Experimental design for retrograde labelling Esr+ LHA-LHb neurons used for ex vivo electrophysiological recordings. (**m**) Representative baseline whole-cell firing traces at rheobase (top), whole-cell firing patterns at maximal depolarization (bottom) of Burst type Esr1+ LHA-LHb neurons in male (blue; n = 37 neurons) and female (pink; n = 18 neurons). (**n**) No significance difference in AP duration and maximal firing frequency of Burst type neurons were found between male (n = 37) and female (n = 18) mice. (**o**) Representative cell-attached baseline tonic firing traces of Burst type Esr1+ LHA-LHb neurons in male (blue) and female (pink) mice. (**p**) No significance difference in tonic firing frequency of Burst type neurons were found between male (n = 18) and female (n = 14) mice. (**q**) Representative baseline whole-cell firing traces at rheobase (top), whole-cell firing patterns at maximal depolarization (bottom) of RS-N type Esr1+ LHA-LHb neurons in male (blue) and female (pink) mice. (**r**) No significance difference in AP duration of RS-N type neurons were found between male and female mice, while neurons in female mice displayed a significant higher maximal firing frequency than male mice (female vs male mice, p < 0.001, DABEST test). Female mice: n = 57, male mice: n = 34. (**s**) Representative cell-attached baseline tonic firing traces of RS-N type Esr1+ LHA-LHb neurons in male (blue) and female (pink) mice. (**t**) No significance difference in tonic firing frequency of RS-N type neurons were found between male (n = 20) and female (n = 16) mice. n = number of neurons, N = number of animals. The data acquired in male and female mice as indicated in panel. For boxplots (**b-d**, **g**, **i-k**), data shown as median, box (25th and 75th percentiles), and whiskers (data points that are not outliers). \**p* < 0.05, \*\**p* < 0.01, \*\*\**p* < 0.001.

**Extended Data Table 1.** Intrinsic parameters of characterized LHA-LHb cell types.

**Supplementary Table 1, Related to Figure 1**
|  | FA-Bk |  | Burst |  | RS-N |  | LS-N |  | LS-W |  | RS-W |  |
| --- | --- | --- | --- | --- | --- | --- | --- | --- | --- | --- | --- | --- |
| n | 20 |  | 49 |  | 65 |  | 37 |  | 40 |  | 19 |  |
| Mean/std | Mean | Std | Mean | Std | Mean | Std | Mean | Std | Mean | Std | Mean | Std |
| Rm (MΩ) | 166,18 | 110,57 | 732,75 | 320,36 | 630,30 | 332,85 | 906,63 | 262,15 | 756,54 | 355,74 | 721,03 | 273,89 |
| Spike latency (ms) | 30,0 | 24,0 | 114,9 | 68,9 | 164,8 | 80,6 | 717,1 | 171,3 | 591,2 | 194,9 | 161,2 | 38,0 |
| Rheo first half freq | 1,65 | 2,91 | 135,50 | 69,04 | 2,25 | 2,61 | 0,29 | 1,61 | 0,00 | 0,00 | 3,30 | 5,02 |
| Rheo second half freq | 0,00 | 0,00 | 0,00 | 0,00 | 0,00 | 0,00 | 1,93 | 2,49 | 2,79 | 3,23 | 0,00 | 0,00 |
| AP thr (mV) | -47,45 | 7,35 | -39,09 | 3,96 | -39,40 | 3,85 | -38,54 | 3,62 | -32,41 | 6,00 | -31,36 | 5,29 |
| Max upstroke (dt/dV) | 510,6 | 116,6 | 386,6 | 103,6 | 447,2 | 91,7 | 408,0 | 89,2 | 272,5 | 109,8 | 222,4 | 81,9 |
| AP half-width (ms) | 0,38 | 0,06 | 0,50 | 0,16 | 0,40 | 0,11 | 0,57 | 0,10 | 1,03 | 0,35 | 1,10 | 0,43 |
| Max downstroke (dt/dV) | -248,8 | 56,8 | -194,4 | 69,3 | -247,9 | 73,1 | -164,1 | 33,6 | -92,3 | 32,9 | -77,5 | 24,4 |
| Conv AHP ampl | 0,63 | 2,82 | 15,94 | 5,13 | 4,61 | 8,63 | 9,85 | 10,16 | 15,38 | 11,76 | 22,11 | 5,95 |
| ADP rise (mV) | 40,63 | 25,59 | 0,19 | 1,31 | 3,48 | 4,45 | 5,39 | 7,70 | 2,45 | 3,98 | 0,00 | 0,00 |
| AHP rise (ms) | 2,54 | 9,28 | 5,00 | 23,67 | 6,28 | 11,74 | 7,03 | 18,21 | 24,64 | 35,77 | 2,43 | 1,75 |
| Freq ISI 1 (Hz) 2xRheo | 107,7 | 59,6 | 186,9 | 63,7 | 42,3 | 26,0 | 18,7 | 18,9 | 12,8 | 11,5 | 33,5 | 14,7 |
| Adaptation ISI1 last 2xRheo | 15,8 | 16,6 | 9,5 | 7,5 | 58,1 | 24,2 | 225,4 | 169,1 | 171,6 | 166,3 | 36,1 | 22,7 |
| Adaptation ISI2 last 2xRheo | 19,0 | 18,6 | 15,9 | 23,7 | 58,8 | 19,6 | 156,4 | 64,7 | 96,7 | 24,1 | 51,3 | 22,1 |
| Freq MAX (Hz) | 164,40 | 52,45 | 88,27 | 47,45 | 128,43 | 60,52 | 63,32 | 23,66 | 34,35 | 13,70 | 30,79 | 11,61 |
| Freq ISI 1 (Hz) Max | 279,9 | 78,2 | 238,0 | 75,7 | 163,0 | 69,4 | 55,4 | 50,7 | 13,9 | 13,4 | 61,9 | 20,8 |
| Freq ISI 2 (Hz) Max | 298,4 | 82,5 | 232,3 | 81,9 | 184,5 | 74,8 | 43,9 | 34,6 | 36,5 | 14,3 | 53,4 | 14,6 |
| Adaptation ISI1 Last Max | 36,2 | 18,0 | 30,6 | 14,5 | 71,0 | 25,1 | 375,8 | 381,8 | 404,6 | 235,0 | 50,2 | 23,5 |
| Adaptation ISI2 Last | 33,7 | 16,3 | 31,7 | 14,8 | 60,3 | 14,0 | 271,5 | 248,9 | 110,5 | 38,6 | 49,4 | 10,3 |
| Post step potential (mV) | -20,04 | 5,22 | -3,45 | 4,42 | -1,78 | 6,32 | 0,14 | 5,53 | -1,98 | 5,07 | -0,81 | 4,33 |

**Extended Data Table 2.** Dendritic morphology, dendritic sholl analysis and somatic morphometry parameters of biocytin filled LHA-LHb neurons by type.

**Supplementary Table 2, Related to Figure 1**
|  | FA-Bk |  | Burst |  | RS-N |  | LS-N |  | LS-W |  | RS-W |  |
| --- | --- | --- | --- | --- | --- | --- | --- | --- | --- | --- | --- | --- |
| Mean/std | Mean | Std | Mean | Std | Mean | Std | Mean | Std | Mean | Std | Mean | Std |
| Dendritic morphology |  |  |  |  |  |  |  |  |  |  |  |  |
| Total length [mm] | 3,395 | 0,523 | 2,575 | 0,452 | 1,802 | 0,624 | 2,709 | 0,292 | 1,820 | 0,427 | 1,863 | 0,564 |
| Total branches [n] | 19,10 | 4,40 | 12,72 | 3,69 | 11,31 | 4,60 | 15,00 | 4,24 | 6,87 | 2,16 | 7,60 | 1,81 |
| Primary branches [n] | 2,07 | 0,27 | 4,31 | 0,94 | 3,81 | 0,83 | 4,09 | 0,30 | 3,87 | 0,83 | 4,40 | 0,89 |
| Dendritic sholl analysis |  |  |  |  |  |  |  |  |  |  |  |  |
| Ending radius | 495,00 | 123,04 | 413,00 | 92,02 | 303,84 | 124,73 | 365,55 | 59,81 | 435,00 | 41,05 | 374,00 | 104,30 |
| Sum intersections [n] | 233,60 | 36,94 | 172,20 | 53,64 | 121,00 | 56,73 | 158,66 | 38,50 | 125,12 | 25,05 | 126,80 | 39,97 |
| Mean intersection [n] | 4,84 | 0,70 | 4,10 | 1,08 | 4,06 | 1,29 | 4,33 | 1,30 | 2,91 | 0,77 | 3,37 | 0,29 |
| Median intersection [n] | 5,00 | 0,81 | 4,00 | 1,05 | 3,84 | 1,72 | 4,66 | 1,87 | 2,56 | 1,23 | 3,30 | 0,83 |
| Max intersections [n] | 10,50 | 2,67 | 8,40 | 2,27 | 8,30 | 2,05 | 8,44 | 1,23 | 6,50 | 1,41 | 7,41 | 0,54 |
| Max intersection radius | 189,00 | 114,35 | 155,00 | 119,65 | 113,84 | 113,91 | 101,11 | 74,06 | 71,25 | 39,07 | 46,00 | 5,47 |
| Ramification index | 8,66 | 3,39 | 6,20 | 3,36 | 6,00 | 1,92 | 5,22 | 2,81 | 5,47 | 2,12 | 4,96 | 2,41 |
| Enclosing radius | 495,00 | 123,04 | 423,00 | 92,02 | 303,84 | 124,73 | 376,66 | 80,93 | 435,00 | 41,05 | 374,00 | 104,30 |
| Critical value | 8,23 | 1,70 | 6,85 | 1,77 | 6,66 | 1,20 | 7,07 | 1,60 | 5,51 | 1,14 | 5,95 | 0,84 |
| Critical radius | 204,32 | 98,41 | 136,28 | 98,03 | 97,37 | 84,37 | 126,66 | 71,96 | 75,04 | 27,99 | 49,47 | 10,19 |
| Mean value | 4,93 | 0,73 | 4,16 | 1,11 | 4,17 | 1,32 | 4,42 | 1,36 | 2,96 | 0,80 | 3,43 | 0,30 |
| Somatic morphometry |  |  |  |  |  |  |  |  |  |  |  |  |
| Area [µm<sup>2</sup>] | 191,98 | 38,96 | 206,25 | 41,46 | 185,52 | 42,33 | 147,69 | 28,30 | 303,96 | 87,87 | 204,39 | 57,37 |
| X [µm] | 22,22 | 3,88 | 19,30 | 2,45 | 18,04 | 2,96 | 20,60 | 2,59 | 26,21 | 3,87 | 23,99 | 1,89 |
| Y1 major [µm] | 8,38 | 1,14 | 12,38 | 2,92 | 10,42 | 1,93 | 8,76 | 2,30 | 14,77 | 3,20 | 9,13 | 3,76 |
| Y [µm] | 10,71 | 1,31 | 12,86 | 2,87 | 12,67 | 2,01 | 10,59 | 2,13 | 14,69 | 2,32 | 10,22 | 1,35 |
| Y2 minor [µm] | 7,92 | 1,00 | 8,54 | 1,85 | 9,70 | 1,61 | 7,97 | 2,08 | 9,40 | 1,76 | 7,76 | 2,61 |
| T score [Y1/Y2] | 1,05 | 1,05 | 1,44 | 0,10 | 1,07 | 0,04 | 1,09 | 0,04 | 1,59 | 0,32 | 1,15 | 0,09 |
| E score [X/Y] | 2,08 | 0,32 | 1,55 | 0,33 | 1,44 | 0,24 | 1,98 | 0,28 | 1,82 | 0,39 | 2,18 | 0,41 |

**Extended Data Table 3.** P-values of pairwise Tukey’s multiple comparisons within One-Way ANOVA test performed on dendritic morphology and somatic morphometry displayed parameters.

|  |  |  |  |  |  |  |  |
| --- | --- | --- | --- | --- | --- | --- | --- |
| Dendrites total length | FA-Bk | Burst | 0,0071 | Soma area [um2] | FA-Bk | Burst | 0,9732 |
|  | FA-Bk | RS-N | 0,0455 |  | FA-Bk | RS-N | 0,2117 |
|  | FA-Bk | LS-N | 0,0000 |  | FA-Bk | LS-N | 0,9992 |
|  | FA-Bk | LS-W | 0,0000 |  | FA-Bk | LS-W | 0,0000 |
|  | FA-Bk | RS-W | 0,0000 |  | FA-Bk | RS-W | 0,9963 |
|  | Burst | RS-N | 0,9914 |  | Burst | RS-N | 0,0631 |
|  | Burst | LS-N | 0,0084 |  | Burst | LS-N | 0,8913 |
|  | Burst | LS-W | 0,0277 |  | Burst | LS-W | 0,0004 |
|  | Burst | RS-W | 0,1133 |  | Burst | RS-W | 1,0000 |
|  | RS-N | LS-N | 0,0018 |  | RS-N | LS-N | 0,4082 |
|  | RS-N | LS-W | 0,0073 |  | RS-N | LS-W | 0,0000 |
|  | RS-N | RS-W | 0,0406 |  | RS-N | RS-W | 0,2838 |
|  | LS-N | LS-W | 1,0000 |  | LS-N | LS-W | 0,0000 |
|  | LS-N | RS-W | 0,9999 |  | LS-N | RS-W | 0,9767 |
|  | LS-W | RS-W | 1,0000 |  | LS-W | RS-W | 0,0075 |
| Primary dendrites [n] | FA-Bk | Burst | 0,0000 | Soma T score | FA-Bk | Burst | 0,0000 |
|  | FA-Bk | RS-N | 0,0000 |  | FA-Bk | RS-N | 0,9643 |
|  | FA-Bk | LS-N | 0,0000 |  | FA-Bk | LS-N | 0,9996 |
|  | FA-Bk | LS-W | 0,0000 |  | FA-Bk | LS-W | 0,0000 |
|  | FA-Bk | RS-W | 0,0000 |  | FA-Bk | RS-W | 0,7314 |
|  | Burst | RS-N | 0,9711 |  | Burst | RS-N | 0,0000 |
|  | Burst | LS-N | 0,3933 |  | Burst | LS-N | 0,0000 |
|  | Burst | LS-W | 0,7388 |  | Burst | LS-W | 0,1548 |
|  | Burst | RS-W | 0,9999 |  | Burst | RS-W | 0,0013 |
|  | RS-N | LS-N | 0,9255 |  | RS-N | LS-N | 0,9958 |
|  | RS-N | LS-W | 0,9880 |  | RS-N | LS-W | 0,0000 |
|  | RS-N | RS-W | 0,9695 |  | RS-N | RS-W | 0,9770 |
|  | LS-N | LS-W | 1,0000 |  | LS-N | LS-W | 0,0000 |
|  | LS-N | RS-W | 0,6236 |  | LS-N | RS-W | 0,8588 |
|  | LS-W | RS-W | 0,8065 |  | LS-W | RS-W | 0,0000 |
| Total dendrites [n] | FA-Bk | Burst | 0,0053 | Soma E score | FA-Bk | Burst | 0,0008 |
|  | FA-Bk | RS-N | 0,1874 |  | FA-Bk | RS-N | 0,9697 |
|  | FA-Bk | LS-N | 0,0001 |  | FA-Bk | LS-N | 0,0000 |
|  | FA-Bk | LS-W | 0,0000 |  | FA-Bk | LS-W | 0,3989 |
|  | FA-Bk | RS-W | 0,0000 |  | FA-Bk | RS-W | 0,9870 |
|  | Burst | RS-N | 0,7609 |  | Burst | RS-N | 0,0241 |
|  | Burst | LS-N | 0,9366 |  | Burst | LS-N | 0,9448 |
|  | Burst | LS-W | 0,0047 |  | Burst | LS-W | 0,3973 |
|  | Burst | RS-W | 0,1046 |  | Burst | RS-W | 0,0056 |
|  | RS-N | LS-N | 0,1894 |  | RS-N | LS-N | 0,0011 |
|  | RS-N | LS-W | 0,0001 |  | RS-N | LS-W | 0,8789 |
|  | RS-N | RS-W | 0,0070 |  | RS-N | RS-W | 0,8449 |
|  | LS-N | LS-W | 0,0239 |  | LS-N | LS-W | 0,0683 |
|  | LS-N | RS-W | 0,3183 |  | LS-N | RS-W | 0,0005 |
|  | LS-W | RS-W | 0,9906 |  | LS-W | RS-W | 0,3389 |

**Extended Data Table 4.** Summary of quantified behavior upon optogenetic manipulation of LHA-LHb pathway in Esr1-cre, Npy-cre, Pv-cre, Gal-cre, Vglut2-cre and C57BL6/J mice.

| Test | Quantified | Figure | Mouse line |  |  |  |  |  |
| --- | --- | --- | --- | --- | --- | --- | --- | --- |
|  |  |  | Esr1-ChR2 | Npy-ChR2 | Pv-ChR2 | Gal-ChR2 | Vglut2-ChR2 | control |
| rtPA | Aversion | 3b | ↑ | = | = | = | ↑ | = |
| State Induction | Aversive behaviors | 3g-h | ↑ | = | n.p. | n.p. | n.p. | = |
| State Induction | Rearing behavior | 3i, | = | ↑ | n.p. | n.p. | n.p. | = |
| Immediate and sustained | cPA | 3l-m, s6q | ↑ | = | n.p. | n.p. | n.p. | = |
| Open field | Distance | S6r,s | ↓ | ↓ | = | = | n.p. | = |
| Open field | Free rearing | 3j, S6t-w | n.p. | ↑ | = | = | n.p. | = |
| Open field | Wall raring | S6t-w | n.p. | = | = | = | n.p. | = |
| Open field | Grooming | S6t-w | n.p. | = | = | = | n.p. | = |
| Open field | Center | S6r | = | = | = | = | n.p. | = |
| Sucrose consumption | Sucrose consumption | S6ä | n.p. | = | = | = | n.p. | = |

**Extended Data Table 5.** Mean, SD and p-values of Mann-Whitney (two-tailed) test with Bonferroni correction performed on the mean block1-zscored pupil area (number of trials indicated in each column) for each genotype (Vglut2-cre, Npy-cre and Esr1-cre) vs control.

|  | Block 1 (n = 50 trials) |  | Block 2 (n = 100 trials) |  |  | Block 3 (n = 50 trials) |  |  |  |
| --- | --- | --- | --- | --- | --- | --- | --- | --- | --- |
| Mann-Whitney (two-tailed) | Mean | S.d | Mean | S.d. | P value (bonf. Corr) | Mean | S.d. | P value (bonf. Corr) | P value (bonf. Corr) |
| C57Bl/6J (N = 5) | -0.018 | 0.380 | -0.130 | 0.516 | 0.352 | -0.090 | 0.400 | 0.698 | 1.781 |
| Vglut2 (N = 5) | 0.023 | 0.353 | 0.5222 | 0.386 | 4.641e-11 | 0.463 | 0.288 | 3.443e-08 | 2.0498 |
| Npy (N = 5) | 0.029 | 0.417 | 0.223 | 0.495 | 0.105 | -0.526 | 0.435 | 5.721e-08 | 2.894e-14 |
| Esr (N = 10) | 0.016 | 0.352 | 0.611 | 0.347 | 9.282e-14 | 0.751 | 0.369 | 4.355e-12 | 0.021 |

**Extended Data Table 6.** Mean, SD and p-values of Mann-Whitney (one-tailed) test with Bonferroni correction performed on the mean tuning scores (absolute value, number of mice indicated in each column) for each genotype (Vglut2-cre, Npy-cre and Esr1-cre) vs control (C57Bl/6J).

|  | C57Bl/6J |  | Vglut2 |  |  | Npy |  |  | Esr1 |  |  |
| --- | --- | --- | --- | --- | --- | --- | --- | --- | --- | --- | --- |
| N (mice) | 5 |  | 5 |  |  | 5 |  |  | 10 |  |  |
| Mann-Whitney<br>(one-tailed) | Mean | S.d | Mean | S.d. | P value<br>(bonf.<br>Corr) | Mean | S.d. | P value<br>(bonf.<br>Corr) | Mean | S.d. | P value<br>(bonf.<br>Corr) |
| Pure tone | 4.147 | 0.973 | 4.079 | 1.904 | 1.738 | 5.138 | 0.927 | 0.286 | 3.559 | 1.959 | 2.205 |
| Optogenetics | 1.176 | 0.258 | 3.337 | 0.409 | 0.012 | 1.126 | 0.599 | 1.905 | 1.896 | 0.612 | 0.045 |
| Blue noise | 2.253 | 0.461 | 1.903 | 0.863 | 2.536 | 2.254 | 0.304 | 1.643 | 1.296 | 0.782 | 2.955 |
| Air puff | 2.713 | 0.377 | 2.246 | 0.331 | 2.917 | 2.222 | 0.442 | 2.833 | 1.713 | 0.708 | 2.992 |

**Extended Data Table 7.** Mean, SD and p-values of Mann-Whitney (one-tailed) test with Bonferroni correction performed on the positive tuning scores (number of units indicated in each column) for each unit type, for each genotype (C57Bl/6J, Vglut2-cre, Npy-cre and Esr1-cre).

| Positive tunings |  |  |  |  |  |  |  |  |  |  |  |  |
| --- | --- | --- | --- | --- | --- | --- | --- | --- | --- | --- | --- | --- |
| C56Bl6/J |  |  |  |  |  |  |  |  |  |  |  |  |
|  | All units |  |  | Wide-spiking waveform units |  |  |  | Narrow-spiking waveform units |  |  |  |  |
| N (mice) | 5 |  |  | 5 |  |  |  | 5 |  |  |  |  |
| Mann-Whitney (one-tailed) | Mean | n | S.d | Mean | n | S.d. | P value (bonf. Corr) | Mean | n | S.d. | P value (bonf. Corr) | P value (bonf. Corr) |
| Pure tone | 5.814 | 218 | 3.063 | 5.767 | 176 | 3.031 | 2.678 | 6.067 | 35 | 3.236 | 1.851 | 1.716 |
| Optogenetics | 3.165 | 29 | 1.157 | 3.138 | 22 | 1.091 | 2.886 | 3.203 | 6 | 1.571 | 2.032 | 2.024 |
| Blue noise | 3.201 | 190 | 1.413 | 3.201 | 150 | 1.387 | 2.768 | 3.298 | 34 | 1.522 | 1.897 | 1.789 |
| Air puff | 3.598 | 247 | 1.621 | 3.626 | 196 | 1.619 | 2.662 | 3.547 | 44 | 1.704 | 2.891 | 2.705 |
| VGlut2-cre |  |  |  |  |  |  |  |  |  |  |  |  |
|  | All units |  |  | Wide-spiking waveform units |  |  |  | Narrow-spiking waveform units |  |  |  |  |
| N (mice) | 5 |  |  | 5 |  |  |  | 5 |  |  |  |  |
| Mann-Whitney (one-tailed) | Mean | n | S.d | Mean | n | S.d. | P value (bonf. Corr) | Mean | n | S.d. | P value (bonf. Corr) | P value (bonf. Corr) |
| Pure tone | 6.536 | 285 | 3.386 | 6.452 | 244 | 3.328 | 2.483 | 6.847 | 33 | 3.555 | 2.004 | 1.790 |
| Optogenetics | 4.764 | 293 | 2.398 | 4.717 | 253 | 2.319 | 2.787 | 4.733 | 33 | 2.742 | 2.024 | 2.106 |
| Blue noise | 3.505 | 196 | 1.667 | 3.451 | 171 | 1.647 | 2.255 | 3.569 | 21 | 1.710 | 2.445 | 2.106 |
| Air puff | 3.247 | 292 | 1.438 | 3.223 | 256 | 1.424 | 2.203 | 3.398 | 31 | 1.620 | 1.035 | 0.8400 |
| NPY-cre |  |  |  |  |  |  |  |  |  |  |  |  |
|  | All units |  |  | Wide-spiking waveform units |  |  |  | Narrow-spiking waveform units |  |  |  |  |
| N (mice) | 5 |  |  | 5 |  |  |  | 5 |  |  |  |  |
| Mann-Whitney (one-tailed) | Mean | n | S.d | Mean | n | S.d. | P value (bonf. Corr) | Mean | n | S.d. | P value (bonf. Corr) | P value (bonf. Corr) |
| Pure tone | 6.399 | 234 | 3.278 | 6.545 | 200 | 3.364 | 2.028 | 5.259 | 32 | 2.318 | 0.261 | 0.171 |
| Optogenetics | 3.195 | 31 | 1.178 | 3.194 | 30 | 1.199 | 3.000 | 3.217 | 1 | 0 | NaN | NaN |
| Blue noise | 3.047 | 180 | 1.358 | 3.185 | 151 | 1.339 | 1.432 | 2.341 | 27 | 1.300 | 0.178 | 0.073 |
| Air puff | 3070 | 176 | 1.269 | 3.166 | 148 | 1.237 | 1.919 | 2.594 | 27 | 1.359 | 0.702 | 0.458 |
| Esr-cre |  |  |  |  |  |  |  |  |  |  |  |  |
|  | All units |  |  | Wide-spiking waveform units |  |  |  | Narrow-spiking waveform units |  |  |  |  |
| N (mice) | 5 |  |  | 5 |  |  |  | 5 |  |  |  |  |
| Mann-Whitney (one-tailed) | Mean | n | S.d | Mean | n | S.d. | P value (bonf. Corr) | Mean | n | S.d. | P value (bonf. Corr) | P value (bonf. Corr) |
| Pure tone | 6.309 | 123 | 3.823 | 6.14 | 101 | 3.682 | 2.442 | 7.301 | 21 | 4.415 | 1.100 | 0.917 |
| Optogenetics | 3.828 | 92 | 1.820 | 3.745 | 77 | 1.873 | 1.921 | 4.344 | 13 | 1.505 | 0.433 | 0.281 |
| Blue noise | 2.838 | 78 | 1.853 | 2.735 | 64 | 1.879 | 2.187 | 3.309 | 14 | 1.713 | 1.075 | 0.814 |
| Air puff | 2.959 | 134 | 1.608 | 2.948 | 106 | 1.612 | 2.743 | 3.349 | 27 | 1.633 | 2.827 | 2.673 |

**Extended Data Table 8.** Mean, SD and p-values of Mann-Whitney (one-tailed) test with Bonferroni correction performed on the negative tuning scores (number of units indicated in each column) for each unit type, for each genotype (C57Bl/6J, Vglut2-cre, Npy-cre and Esr1-cre).

| Negative tunings |  |  |  |  |  |  |  |  |  |  |  |  |
| --- | --- | --- | --- | --- | --- | --- | --- | --- | --- | --- | --- | --- |
| C56Bl6/J |  |  |  |  |  |  |  |  |  |  |  |  |
|  | All units |  |  | Wide-spiking waveform units |  |  |  | Narrow-spiking waveform units |  |  |  |  |
| N (mice) | 5 |  |  | 5 |  |  |  | 5 |  |  |  |  |
| Mann-Whitney (one-tailed) | Mean | n | S.d | Mean | n | S.d. | P value (bonf. Corr) | Mean | n | S.d. | P value (bonf. Corr) | P value (bonf. Corr) |
| Pure tone | -4.989 | 32 | 1.663 | -4.961 | 24 | 1.880 | 2.240 | -5.020 | 7 | 0.8370 | 1.749 | 1.392 |
| Optogenetics | -3.889 | 24 | 1.207 | -3.887 | 19 | 1.239 | 3 | -4.112 | 4 | 1.280 | 2.303 | 2.329 |
| Blue noise | -3.990 | 4 | 0.4590 | -4.160 | 3 | 0.3780 | 2.571 | -3.480 | 1 | 0 | 2.400 | 1.500 |
| Air puff | -4.874 | 9 | 0.9550 | -4.649 | 6 | 0.9270 | 2.052 | -5.324 | 3 | 1.022 | 1.636 | 1.143 |
| VGlut2-cre |  |  |  |  |  |  |  |  |  |  |  |  |
|  | All units |  |  | Wide-spiking waveform units |  |  |  | Narrow-spiking waveform units |  |  |  |  |
| N (mice) | 5 |  |  | 5 |  |  |  | 5 |  |  |  |  |
| Mann-Whitney (one-tailed) | Mean | n | S.d | Mean | n | S.d. | P value (bonf. Corr) | Mean | n | S.d. | P value (bonf. Corr) | P value (bonf. Corr) |
| Pure tone | -6.451 | 44 | 3.395 | -6.271 | 37 | 3.022 | 2.796 | -7.723 | 6 | 5.567 | 2.330 | 2.217 |
| Optogenetics | -7.564 | 84 | 4.446 | -7.940 | 72 | 4.565 | 1.676 | -5.478 | 11 | 2.938 | 0.3410 | 0.1890 |
| Blue noise | -4.682 | 10 | 1.086 | -4.784 | 9 | 1.100 | 2.590 | -3.770 | 1 | 0 | 2.182 | 1.800 |
| Air puff | -6.788 | 24 | 3.862 | -6.342 | 22 | 3.525 | 2.147 | -11.465 | 231 | 5.629 | 0.3610 | 0.2720 |
| NPY-cre |  |  |  |  |  |  |  |  |  |  |  |  |
|  | All units |  |  | Wide-spiking waveform units |  |  |  | Narrow-spiking waveform units |  |  |  |  |
| N (mice) | 5 |  |  | 5 |  |  |  | 5 |  |  |  |  |
| Mann-Whitney (one-tailed) | Mean | n | S.d | Mean | n | S.d. | P value (bonf. Corr) | Mean | n | S.d. | P value (bonf. Corr) | P value (bonf. Corr) |
| Pure tone | -6.551 | 31 | 4.363 | -6.689 | 28 | 4.427 | 2.771 | -5.336 | 3 | 2.690 | 2.121 | 2.035 |
| Optogenetics | -6.059 | 31 | 2.973 | -6.177 | 26 | 3.059 | 2.439 | -3.745 | 3 | 0.5090 | 0.272 | 0.274 |
| Blue noise | -3.632 | 3 | 0.2830 | -3.586 | 2 | 0.3830 | 3 | -3.725 | 1 | 0 | 3 | 3 |
| Air puff | -6.153 | 27 | 2.634 | -6.382 | 22 | 2.680 | 2.312 | -4.236 | 4 | 1.499 | 0.4460 | 0.3290 |
| Esr-cre |  |  |  |  |  |  |  |  |  |  |  |  |
|  | All units |  |  | Wide-spiking waveform units |  |  |  | Narrow-spiking waveform units |  |  |  |  |
| N (mice) | 10 |  |  | 10 |  |  |  | 10 |  |  |  |  |
| Mann-Whitney (one-tailed) | Mean | n | S.d | Mean | n | S.d. | P value (bonf. Corr) | Mean | n | S.d. | P value (bonf. Corr) | P value (bonf. Corr) |
| Pure tone | -6.5910 | 3 | 0.848 | -6.4330 | 2 | 1.1350 | 3 | -6.9080 | 1 | 0 | 3 | 3 |
| Optogenetics | -5.4010 | 21 | 2.9710 | -5.6280 | 19 | 3.0270 | 2.2660 | -3.2470 | 2 | 1.1630 | 0.628 | 0.505 |
| Blue noise | -3.0620 | 1 | 0 | -3.0620 | 1 | 0 | 3 | NaN | 0 | NaN | NaN | NaN |
| Air puff | -4.6060 | 9 | 1.1530 | -4.5060 | 8 | 1.1910 | 2.6340 | NaN | 0 | NaN | NaN | NaN |

**Extended Data Table 9.** Mean, SD and p-values of Wilcoxon signed rank (two-tailed) test with Bonferroni correction performed on the mean baseline firing rate (number of units indicated in each row) across blocks for each activity cluster identified in Fig. 3g.

|  | Block1 |  | Block 2 |  |  | Block 3 |  |  |  | Block 4 |  |  |  |  |
| --- | --- | --- | --- | --- | --- | --- | --- | --- | --- | --- | --- | --- | --- | --- |
| Wilcoxon (two-tailed) | Mean | S.d | Mean | S.d. | P value (bonf. Corr) | Mean | S.d. | P value (bonf. Corr) | P value (bonf. Corr) | Mean | S.d. | P value (bonf. Corr) | P value (bonf. Corr) | P value (bonf. Corr) |
| 'Opto' #1<br>(n = 65 units) | 3,752 | 5,183 | 4,990 | 5,841 | 1,079 | 5,474 | 6,047 | 0,125 | 0,184 | 5,270 | 5,858 | 2,046 | 2,671 | 4,982 |
| 'Opto' #2<br>(n = 93 units) | 4,312 | 5,311 | 4,598 | 5,245 | 3,207 | 4,938 | 5,851 | 2,059 | 1,273 | 5,106 | 5,793 | 4,788 | 3,606 | 4,756 |
| 'Air puff' #1<br>(n = 110 units) | 2,829 | 3,670 | 3,856 | 4,290 | 0,048 | 4,550 | 4,929 | 0,002 | 4,008E-05 | 5,039 | 5,258 | 1,688 | 0,227 | 2,085 |
| 'Air puff' #2<br>(n = 65 units) | 2,731 | 3,441 | 2,810 | 3,672 | 3,467 | 3,323 | 3,978 | 0,996 | 0,050 | 4,426 | 5,100 | 1,887 | 0,129 | 1,079 |
| 'State' #4<br>(n = 150 units) | 1,976 | 2,793 | 2,362 | 2,788 | 0,109 | 2,886 | 3,409 | 0,001 | 1,132E-12 | 4,673 | 5,166 | 0,705 | 6,759E-08 | 1,186E-04 |
| 'State' #3<br>(n = 253 units) | 1,964 | 3,106 | 2,413 | 3,294 | 0,014 | 3,974 | 4,817 | 1,652E-15 | 6,013E-08 | 3,272 | 4,582 | 1,869E-07 | 0,021 | 0,048 |
| 'State' #2<br>(n = 128 units) | 3,306 | 4,658 | 4,621 | 5,078 | 3,222E-04 | 4,002 | 5,056 | 0,120 | 1,767 | 3,645 | 4,970 | 0,391 | 0,008 | 1,493 |
| 'State' #1<br>(n = 195 units) | 6,317 | 9,328 | 5,446 | 8,473 | 1,302 | 5,139 | 8,186 | 0,461 | 0,047 | 4,919 | 8,291 | 3,731 | 0,846 | 2,095 |
| 'State' #5<br>(n = 87 units) | 2,273 | 3,351 | 2,824 | 3,844 | 0,229 | 3,847 | 4,639 | 1,912E-04 | 2,209E-04 | 4,137 | 5,661 | 0,057 | 0,048 | 5,211 |
| 'Sound' #1<br>(n = 72 units) | 4,836 | 5,189 | 5,450 | 5,561 | 2,080 | 5,987 | 5,875 | 0,447 | 2,773 | 5,314 | 5,433 | 2,322 | 5,210 | 1,821 |
| 'Sound' #2<br>(n = 72 units) | 3,474 | 4,423 | 4,033 | 4,414 | 0,823 | 4,760 | 4,901 | 0,059 | 0,034 | 5,084 | 5,065 | 1,320 | 0,798 | 3,910 |
| 'Sound' #3<br>(n = 49 units) | 5,449 | 7,031 | 6,042 | 7,164 | 3,163 | 7,100 | 8,165 | 1,076 | 0,883 | 7,695 | 8,903 | 2,864 | 2,025 | 5,087 |
| 'Sound' #4<br>(n = 33 units) | 11,336 | 7,604 | 10,753 | 6,620 | 5,025 | 12,342 | 6,823 | 2,341 | 1,171 | 13,263 | 7,135 | 1,866 | 0,842 | 3,178 |
| 'Sound' #5<br>(n = 189 units) | 6,460 | 7,075 | 6,533 | 7,583 | 3,926 | 7,047 | 8,571 | 4,687 | 2,589 | 7,779 | 8,983 | 5,204 | 1,515 | 1,908 |
| 'Sound' #6<br>(n = 227 units) | 3,075 | 3,765 | 3,904 | 4,477 | 0,059 | 4,860 | 5,232 | 1,088E-05 | 2,267E-08 | 5,722 | 6,862 | 0,115 | 0,003 | 1,456 |

**Extended Data Table 10.**
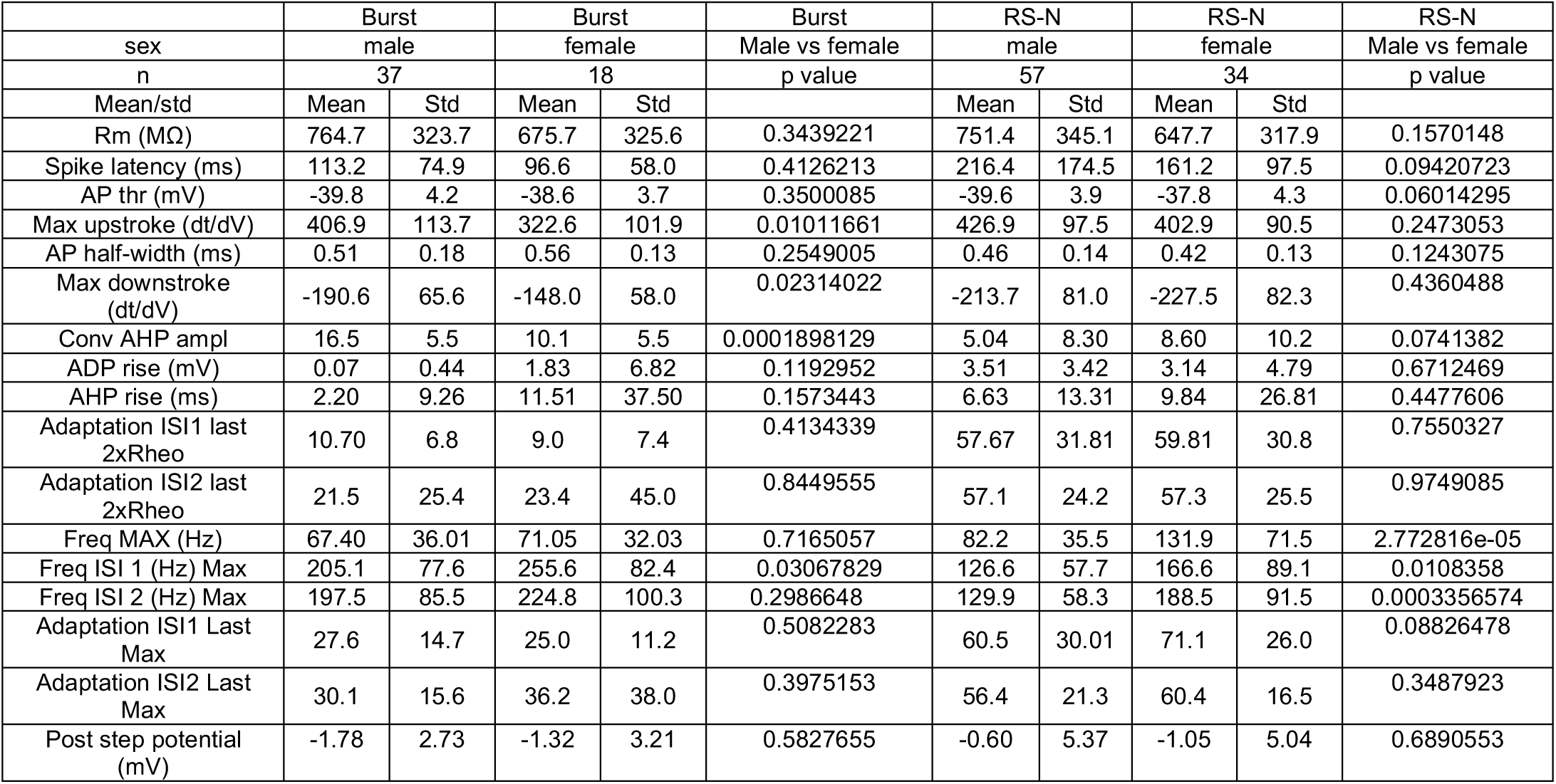
Intrinsic parameters of characterized Burst-type and RS-N type Esr1+ LHA-LHb in male and female mice at baseline.

**Supplementary Table 10, Related to Figure 10**
|  | Burst |  | Burst |  | Burst | RS-N |  | RS-N |  | RS-N |
| --- | --- | --- | --- | --- | --- | --- | --- | --- | --- | --- |
| sex | male |  | female |  | Male vs female | male |  | female |  | Male vs female |
| n | 37 |  | 18 |  | p value | 57 |  | 34 |  | p value |
| Mean/std | Mean | Std | Mean | Std |  | Mean | Std | Mean | Std |  |
| Rm (MΩ) | 764.7 | 323.7 | 675.7 | 325.6 | 0.3439221 | 751.4 | 345.1 | 647.7 | 317.9 | 0.1570148 |
| Spike latency (ms) | 113.2 | 74.9 | 96.6 | 58.0 | 0.4126213 | 216.4 | 174.5 | 161.2 | 97.5 | 0.09420723 |
| AP thr (mV) | -39.8 | 4.2 | -38.6 | 3.7 | 0.3500085 | -39.6 | 3.9 | -37.8 | 4.3 | 0.06014295 |
| Max upstroke (dt/dV) | 406.9 | 113.7 | 322.6 | 101.9 | 0.01011661 | 426.9 | 97.5 | 402.9 | 90.5 | 0.2473053 |
| AP half-width (ms) | 0.51 | 0.18 | 0.56 | 0.13 | 0.2549005 | 0.46 | 0.14 | 0.42 | 0.13 | 0.1243075 |
| Max downstroke (dt/dV) | -190.6 | 65.6 | -148.0 | 58.0 | 0.02314022 | -213.7 | 81.0 | -227.5 | 82.3 | 0.4360488 |
| Conv AHP ampl | 16.5 | 5.5 | 10.1 | 5.5 | 0.0001898129 | 5.04 | 8.30 | 8.60 | 10.2 | 0.0741382 |
| ADP rise (mV) | 0.07 | 0.44 | 1.83 | 6.82 | 0.1192952 | 3.51 | 3.42 | 3.14 | 4.79 | 0.6712469 |
| AHP rise (ms) | 2.20 | 9.26 | 11.51 | 37.50 | 0.1573443 | 6.63 | 13.31 | 9.84 | 26.81 | 0.4477606 |
| Adaptation ISI1 last 2xRheo | 10.70 | 6.8 | 9.0 | 7.4 | 0.4134339 | 57.67 | 31.81 | 59.81 | 30.8 | 0.7550327 |
| Adaptation ISI2 last 2xRheo | 21.5 | 25.4 | 23.4 | 45.0 | 0.8449555 | 57.1 | 24.2 | 57.3 | 25.5 | 0.9749085 |
| Freq MAX (Hz) | 67.40 | 36.01 | 71.05 | 32.03 | 0.7165057 | 82.2 | 35.5 | 131.9 | 71.5 | 2.772816e-05 |
| Freq ISI 1 (Hz) Max | 205.1 | 77.6 | 255.6 | 82.4 | 0.03067829 | 126.6 | 57.7 | 166.6 | 89.1 | 0.0108358 |
| Freq ISI 2 (Hz) Max | 197.5 | 85.5 | 224.8 | 100.3 | 0.2986648 | 129.9 | 58.3 | 188.5 | 91.5 | 0.0003356574 |
| Adaptation ISI1 Last Max | 27.6 | 14.7 | 25.0 | 11.2 | 0.5082283 | 60.5 | 30.01 | 71.1 | 26.0 | 0.08826478 |
| Adaptation ISI2 Last Max | 30.1 | 15.6 | 36.2 | 38.0 | 0.3975153 | 56.4 | 21.3 | 60.4 | 16.5 | 0.3487923 |
| Post step potential (mV) | -1.78 | 2.73 | -1.32 | 3.21 | 0.5827655 | -0.60 | 5.37 | -1.05 | 5.04 | 0.6890553 |

